# Engineering immunogens that select for specific mutations in HIV broadly neutralizing antibodies

**DOI:** 10.1101/2023.12.15.571700

**Authors:** Rory Henderson, Kara Anasti, Kartik Manne, Victoria Stalls, Carrie Saunders, Yishak Bililign, Ashliegh Williams, Pimthada Bubphamala, Maya Montani, Sangita Kachhap, Jingjing Li, Chuancang Jaing, Amanda Newman, Derek Cain, Xiaozhi Lu, Sravani Venkatayogi, Madison Berry, Kshitij Wagh, Bette Korber, Kevin O. Saunders, Ming Tian, Fred Alt, Kevin Wiehe, Priyamvada Acharya, S. Munir Alam, Barton F. Haynes

## Abstract

Vaccine development targeting rapidly evolving pathogens such as HIV-1 requires induction of broadly neutralizing antibodies (bnAbs) with conserved paratopes and mutations, and, in some cases, the same Ig-heavy chains. The current trial-and-error search for immunogen modifications that improve selection for specific bnAb mutations is imprecise. To precisely engineer bnAb boosting immunogens, we used molecular dynamics simulations to examine encounter states that form when antibodies collide with the HIV-1 Envelope (Env). By mapping how bnAbs use encounter states to find their bound states, we identified Env mutations that were predicted to select for specific antibody mutations in two HIV-1 bnAb B cell lineages. The Env mutations encoded antibody affinity gains and selected for desired antibody mutations *in vivo*. These results demonstrate proof-of-concept that Env immunogens can be designed to directly select for specific antibody mutations at residue-level precision by vaccination, thus demonstrating the feasibility of sequential bnAb-inducing HIV-1 vaccine design.

## Introduction

The discovery of effective strategies for rapid design of immunogens to induce HIV-1 broadly neutralizing antibodies (bnAbs) is critical to vaccine design for HIV-1 and for other highly variable viruses (*1, 2*). A wealth of structural and B cell clonal lineage information has demonstrated that bnAb properties make them especially difficult to induce through vaccination. These include the presence of highly improbable functional amino acid mutations (*3*), autoreactivity (*4*), and rare bnAb B cell precursors (*1*). Mutation-guided B cell lineage immunogen design is a strategy to overcome immunogen subdominance of bnAb B cell precursors by sequential immunogen designs aimed at selection of B cells with B cell receptors (BCRs) containing functional yet rare somatic mutations (*5*). In bnAb unmutated common ancestor (UCA) knock-in mouse models that guarantee the presence of bnAb precursors (*6, 7*), induction of bnAb breadth and potency has nonetheless proved challenging. (*8*).

Antibodies mature by acquiring affinity-enhancing amino acid substitutions by somatic hypermutation (*9, 10*). By cycling through antigen recognition and somatic hypermutation, BCRs evolve to develop enhanced affinity (*10*). While a repertoire of low to high-affinity BCRs to target antigens survive during these cycles, higher affinity correlates with enhanced survival (*9, 11*). For HIV-1 vaccine development, the goal is to design bnAb-inducing immunogens such that the affinity for the bnAb or its intermediate forms is greater than the affinity for the bnAb precursor (*2, 12*). Such an affinity gradient could, in principle, enhance the survival of B cells bearing desired HIV-1 bnAb mutations. Many well-defined bnAb clonal lineages targeting conserved Env epitopes exist providing the data sets needed for this approach.

For some HIV-1 bnAb lineages such as those targeting the envelope (Env) V3-glycan bnAb site, variability in heavy and light chain gene usage presents a hurdle for consistent vaccine induction of heterologous neutralizing responses (*13, 14*). Though similarities among V3-glycan bnAbs exist, the heterogeneous nature of individual antibody contacts and their variable bound state orientations complicate vaccine immunogen development. Multiple bnAbs targeting the N332-glycan supersite have been identified from infected HIV-1 individuals including DH270(*15*), PGT128(*16*), PGT135(*14*), PGT121(*17*), and BG18(*18*), which utilize distinct pairings of variable heavy (V_H_) and variable light (V_L_) gene segments(*14, 15, 19*).

Moreover, V3-glycan bnAbs require long HCDR3 regions to bind to Env necessitating the utilization of rare bnAb B cell precursors (*1*). Thus, design of immunogens that can induce many types of V3-glycan bnAb B cell lineages will be necessary to induce a sufficient number of V3-glycan bnAb precursors to result in a robust bnAb response to that epitope.

In contrast with the varied germline gene usage by the V3-glycan bnAbs, the CH235 CD4-mimicking CD4 binding site (bs) VH1-46-utilizing class of bnAbs exclusively uses the VH1-46 gene and has a conserved mode of Env binding that is similar to that of CD4 (*20*). Many high-resolution crystal structures of these antibodies and medium-resolution cryo-EM structures of their bound state complexes with stabilized, soluble Envs, have been obtained for attempts at structure-based design (*8, 14–17, 21–25*). Designs aimed at eliciting bnAb responses based on these structures have yet to elicit mature bnAb responses with neutralization potency and breadth (*8, 13, 14*). Thus, what is lacking is a directed immunogen design process capable of systematically selecting for specific HIV-1 bnAb BCR mutations in a B cell lineage.

Here, we used molecular dynamics simulation combined with Markov state modeling (MSM) to probe, in atomic detail and with high temporal resolution, the encounter states that make up the association process. We examined the association pathways for the mature DH270 V3-glycan and CH235 CD4bs VH1-46-class of bnAbs. Each bnAb mutation can contribute to both association and dissociation depending on its role in reaching and leaving the bound state. We reasoned that knowledge of bnAb association and dissociation mechanisms would enable us to modify our designs to amplify the effects key antibody mutations have on bnAb lineage antibody affinity.

Our results showed that the antibodies reach the bound state from collisions across a large area of the Env near the target epitope. Epitope recognition was accomplished primarily through rotations about the Env surface to bring the paratope into position and was facilitated by specific contacts between Env and antibody. This information was combined with cryo-EM bnAb-Env structures to clarify dissociation mechanisms in the context of the defined association mechanisms and to identify immunogen modifications that enhance affinity for specific antibody mutations. These designs enhanced the selection of targeted antibody mutations *in vivo*.

Thus, these data demonstrated a process whereby a detailed knowledge of each antibody residue’s role in facilitating encounters with the immunogen leading to a stable interaction can enable precise Env designs capable of selecting for specific antibody mutations in human B cell lineages.

## Results

### Molecular dynamics simulation locates a V3-glycan bnAb antibody-antigen bound pose

The DH270 V3-glycan antibody lineage is a promising target due to its relatively limited somatic mutation compared to other HIV bnAbs and the detailed knowledge of Env sequence diversity available from longitudinal viral sequencing of the HIV-1-infected person as the lineage developed (*15*). An autologous Env selected for a critical, improbable DH270 clonal lineage intermediate G57R mutation that confers heterologous neutralization in a DH270 UCA knockin mouse model. This was accomplished via the elimination of two potential N-linked glycosylation sites (PNGS) in the V1 loop that, when glycosylated, sterically hindered DH270 antibody access to the epitope (*8*). Several of the twelve lineage mutations needed to confer 90% of the mature DH270.6 (*26*) were observed individually in a vaccination study, but co-occurrence was relatively rare (*8*), suggesting weaker selection of those residues. To track the role that key mutations play in the association process, we employed a molecular dynamics (MD)-based search for collision-induced encounter states (Figure 1A).

**Figure 1.**
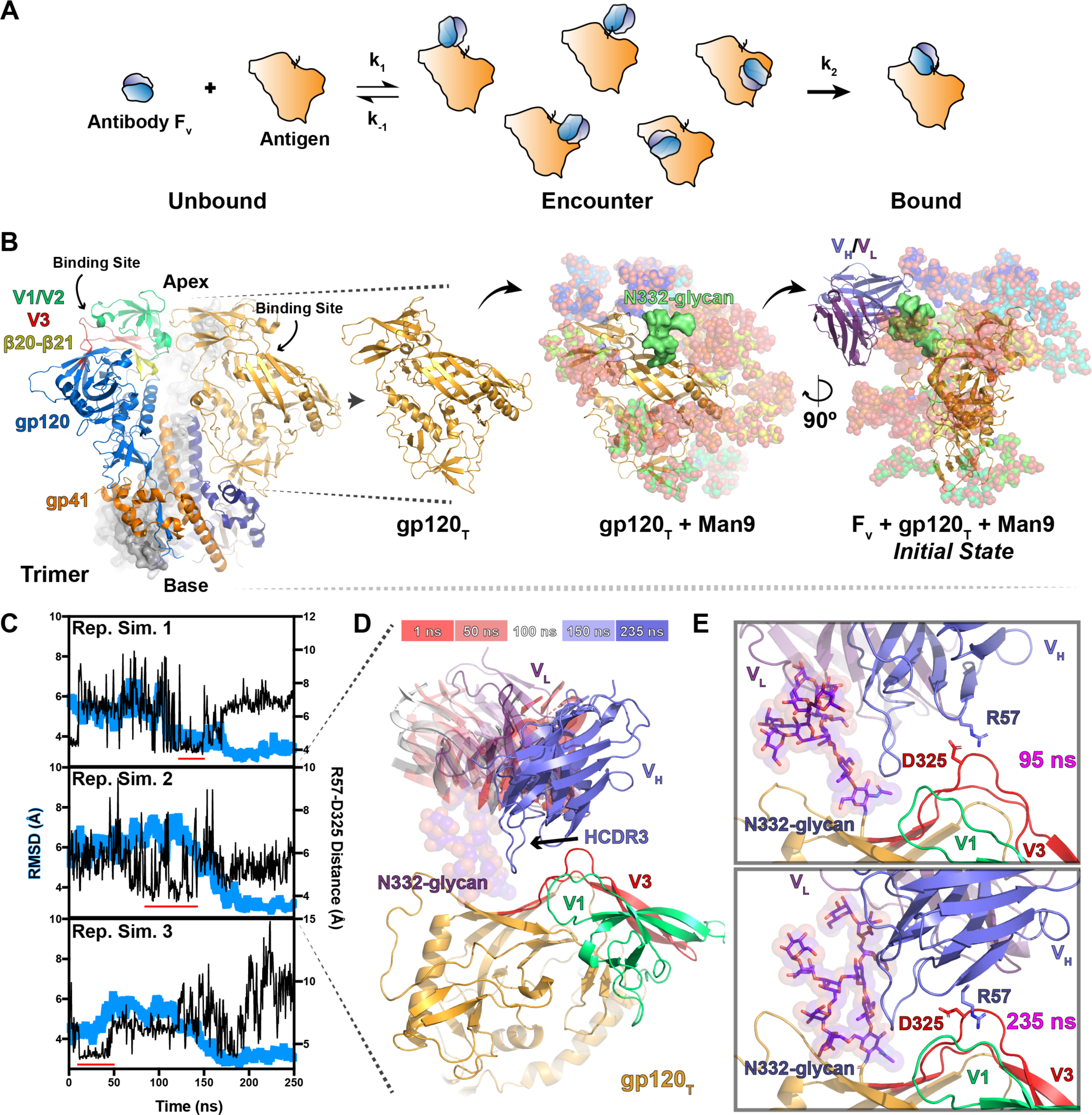
Molecular dynamics simulation of DH270.6 association with a truncated CH848 HIV-1 Env gp120. **A)** Schematic representation of an antibody Fv and gp120 antigen associating through an intermediate encounter ensemble. Transition rates k1 and k-1 represent the ensemble rate of forming a collision-induced encounter state and dissociating to the unbound conformation, respectively, while k2 describes the conversion rate from an encounter intermediate to the bound complex. **B)** (left) An HIV-1 Env trimer ectodomain structure. The protomer on the left highlights gp120 (blue) allosteric elements V1/V2 (green), V3 (red), β20- β21 (yellow), as well as gp41 (orange). The protomer to the right highlights the gp120 region that is extracted (gp120T) from the prefusion closed state structure (orange) and gp41 (purple). The third protomer of the trimer is represented in a grey surface. (middle) The man9 glycosylated gp120_T_ highlights the position of the N332-glycan (green surface). (right) The man9 glycosylated gp120_T_ with the DH270.6 antibody Fv positioned near the N332-glycan. **C)** RMSD trajectories (light blue) of three representative transitions from encounter states to the bound state. Distances between DH270.6 heavy chain R57 and gp120 D325 (black) show close interactions (<3.0 Å, red line) before reaching the bound state. **D)** Representative encounter to bound transition showing rotation of the DH270.6 Fv (fade from red to white to blue) around the N332-glycan (purple/red spheres). **E)** Frames from the representative simulations in an R57 to D325 interactive encounter transition (top) and the final, non-interactive R57 to D325 bound state.

The V3-glycan targeting DH270.6 bnAb interacts with the CH848 HIV-1 viral variant Env ectodomain through interaction with the V3 GDI(K/R) motif and an adjacent N-linked glycan at position N332. Since the antibody-Env interaction does not involve quaternary trimer interactions, we first asked whether the simulation of a single truncated Env gp120 (gp120_T_) extracted from a prefusion closed-state trimer structure would present a stable V3-glycan epitope (Figure 1B). We used an Env sequence isolated from the CH848-infected individual at 949 days post-infection (CH848.d949.10.17 (*15*); referred to here as CH848-d949) (*8*). Five hundred independent, 1.1-microsecond simulations of a Man9 glycosylated gp120_T_ revealed minimal (average RMSD 2.22 Å) overall structural differences from the closed-state Env conformation (Supplemental Figure 1). The most prominent changes occurred at the interprotomer contact loops and variable loops V1, V2, V4, and V5. As the variable loops are flexible, and the interprotomer contact regions are distal from the DH270.6 epitope, we concluded that this flexibility would not significantly affect relevant encounter state formation. Free gp120s typically present open-state conformations of the V1/V2 and V3 elements where the bridging sheet has formed, resulting in an RMSD greater than 6 Å relative to the closed state. The results indicated that a free gp120 in the closed state will not transition to this open-state conformation over the simulation timescales.

Therefore, these data demonstrated that the gp120_T_ V3-glycan epitope would be maintained in simulations involving binding to an antibody V_H_/V_L_.

Next, we prepared a system consisting of gp120_T_ and the variable heavy and light chains, V_H_/V_L_, of DH270.6. A molecular simulation-based analysis of protein-protein interactions suggested that encounter states with both partners oriented near the native bound conformation were most relevant for association (*27*). Thus, we placed the antibody near the epitope with the third heavy chain complementary determining region (HCDR3) positioned toward the N332 residue (Figure 1B). To diversify the V_H_/V_L_-gp120_T_ conformations, we launched four hundred independent 250 nanosecond (ns) simulations from this state. This yielded a wide variety of V_H_/V_L_ orientations relative to the Env epitope (Supplemental Figure 2A). Four antibody residues form the N332-glycan interaction interface: heavy chain residues D115 and light chain residues Y48, Y51, and S58 (Supplemental Figure 2B). Of the four hundred simulations, ∼100 did not form any of these interactions, while ∼50 simulations formed all four contacts. This assortment of interactions suggested that this set of conformations contained sufficient contact and orientation variability to initiate an MD-based search for relevant encounter states and the bound pose.

Here, we used a simulation technique referred to as adaptive sampling (*28*) to explore encounter states and the pathways that lead to the bound state. The adaptive sampling technique used the final states from the 400, 250 ns simulation set to launch the first of an iterative series of short, 250 ns simulation sets, each iteration involving 400 independent simulations, to search for important encounter states and the transitions that make up the association pathway. New iterations are initiated from potential transition states identified in the previous simulations to increase the rate of finding new encounter states, the states that connect them, and the bound state. Here, we conducted seventeen adaptive iterations, including 1,000 simulations launched from fully uncoupled states in multiple antibody V_H_/V_L_ orientations relative to the gp120_T_ (Supplemental Figure 3), yielding nearly two milliseconds of simulation time.

Forty-six of these simulations transitioned from encounter states to the bound state. This occurred through rotation of the antibody around the bound N332-glycan, allowing the antibody to form interactions with the GDIK motif (Figure 1E and F). Electrostatic interactions between antibody heavy chain R57 and gp120_T_ D325 often preceded bound state formation, suggesting this interaction aided in directing the rotation (Figure 1E). Of note, D325 was identified as a significant neutralization sensitivity signature, and the mutations N325 or K325 are significant resistance signatures for DH270 lineage bnAbs 10-1074, PGT121, and PGT128(*29*), consistent with D325 mutations conferring *in vivo* escape to PGT121 (*30*) or 10-1074 (*31*). These neutralization signatures support our MD findings and suggest that DH270 interactions with D325, particularly with V_H_ R57 in the binding transition, may be relevant to other V3 glycan bnAbs. These data demonstrated that MD simulation can locate encounter states and the correct bound conformations for these intricate interactions, starting from an unbound bnAb state.

### DH270.6 bnAb association pathways identify vaccine design targets

The completed simulation set provided detailed structural information for different encounter states and their exchange with nearby states, leading to the known bnAb bound state conformation. An MSM of the association process prepared using the simulated set described this process as time-dependent, memoryless probabilistic jumps between states. States were defined kinetically based on time-dependent changes in distances between known binding site residue contacts using a time-lagged independent component analysis (TICA) transformation method. This transformation identified slowly relaxing degrees of freedom in the system, which could then be clustered for encounter state assignments. These encounter state assignments and their connectivity to other states could then be used to construct a Markov model. Thus, the antibody association pathways could be defined by the relative exchange between states and each state’s probability of moving toward the bound state, defined by the so-called committor distribution, which determines forward transition probabilities based on the MSM (*32*) (Figure 2A, Supplemental Figures 4 to 6). The association pathway provided a map of transitions that can be used to identify target sites for modification of the association process and, therefore, the association rate, k_a_.

**Figure 2.**
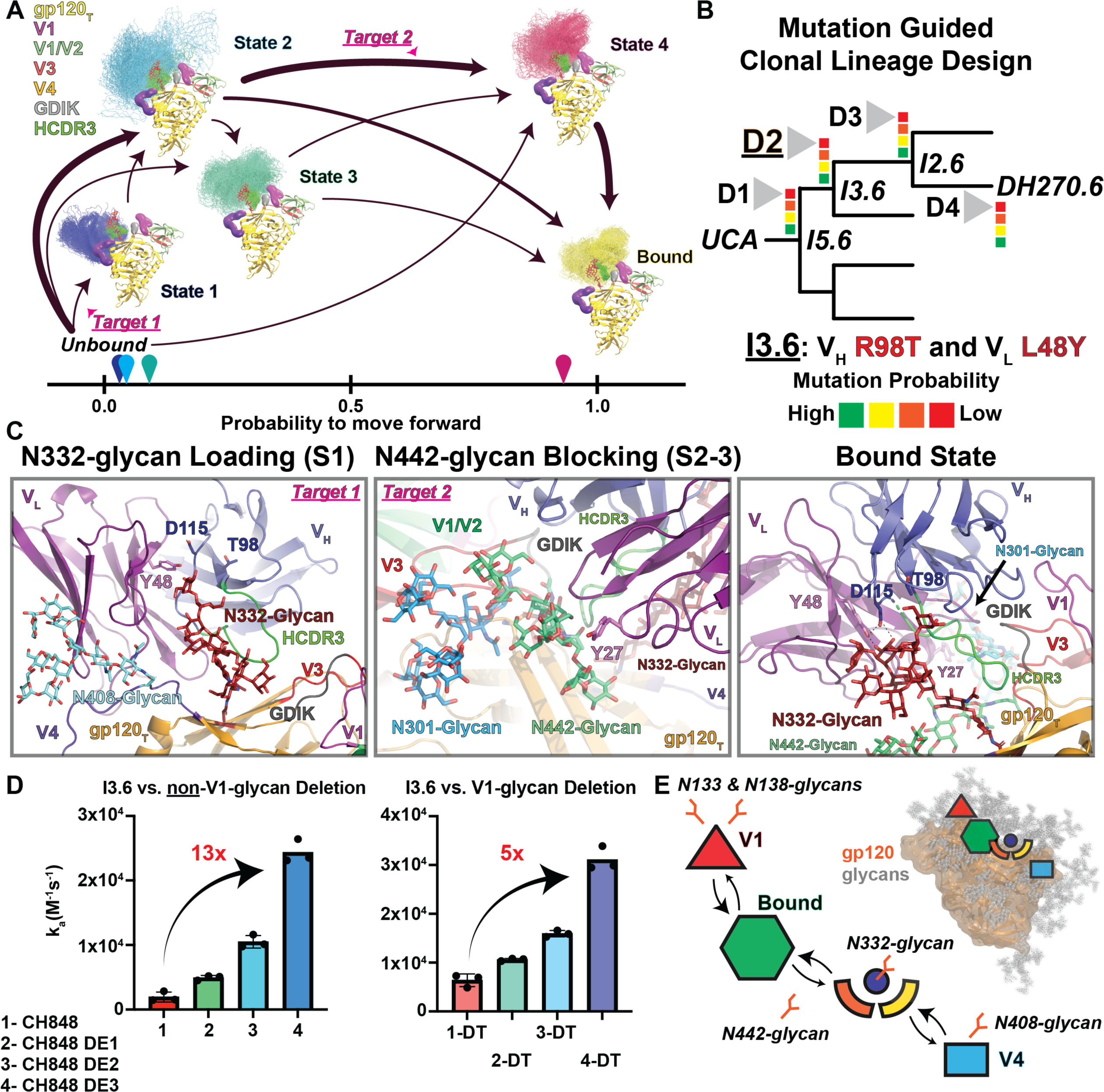
The association pathway for the DH270.6 Fv identifies design targets. **A)** Association pathway for DH270.6 determined from an MSM prepared from the simulated set. Important regions of the gp120T (blue) include the V1 loop (purple), the V4 loop (light orange), the GDIK motif (grey), and the antibody HCDR3 (bright green). A single gp120 state is used throughout, with 50 representative samples of the antibody shown for each state (ensemble of line structures). Arrows show transition paths with sizes adjusted to indicate each transition’s relative flux. Structures are positioned based on the probability that each state will move forward to the bound state. Transitions targeted for design are highlighted (pink). **B)** Graphical representation of lineage-based design depicting a simplified phylogenetic tree leading from the inferred UCA and intermediates to the mature DH270.6 bnAb. Design targets D1-D4, each targeting low probability mutations (red), with the I3.6 targeting D2 identified as the primary focus for design. The I3.6 intermediate contains two key improbable mutations, the V_H_ R98T and V_L_ L48Y. **C)** (left) Representative state from State 1 highlighting the position of the N408-glycan and the contacts between the design target D2 mutation sites V_H_ T98 and V_L_ Y48. (middle) Representative state from State 3 highlighting the position of the N442-glycan relative to the N301-glycan and its bound state interactive Y27 residue sidechain. (right) The bound state conformation shows interaction with the N332-glycan and N301-glycan and the displaced position of the N442-glycan. **D)** (left) Association rates determined by SPR for the parent CH848 Env SOSIP ectodomain and each design (right) Association kinetics for a CH848 parent and design in which two glycans in the V1 loop have been eliminated via mutation (referred to as DT). Arrows and values indicate the fold change in k_a_ between the parent and combined mutations. Three replicate measures are shown with error bars indicating the standard deviation. **E)** (lower left) Simplified graphical representation of encounter transition in the DH270.6 interaction with gp120_T_. (upper right) Shapes placed in their relative positions on the gp120_T_ (orange) map encounter states to the structure. Encounter state 1 maps to the cyan V4 block while Encounter state 2 and 3 map to the orange and yellow.

Our immunogen design targets were defined by functional improbable mutations in each inferred intermediate antibody along the DH270.6 maturation pathway that are critical for achieving neutralization breadth (Figure 2B) (*3*). Our previous efforts identified a design capable of selecting key mutations in the first intermediate, I5 (*8*). Specifically, we found that eliminating N-linked glycan sites at positions N133D and N138T in the CH848-d949 V1 loop (DT) enhanced DH270.UCA affinity and enhanced selection of a key heavy chain mutation, G57R, in DH270.UCA knockin mice (*8*). We, therefore, focused here on two key mutations in the DH270 B cell lineage intermediate antibody, I3: heavy chain R98T and light chain L48Y (Figure 2B).

The association path for the V3-glycan bnAb, DH270.6, proceeded through three encounter states, Encounter States 1 through 3, that reached the bound state predominantly through a pre-bound state, state 4 (Figure 2A). Encounter State 1 involved coupling of the light chain to the gp120_T_ outer domain, adjacent to the V4 loop (Figure 2C left, Supplemental Figure 7A). With the HCDR3 loop oriented toward the N332-glycan, this state acted to present the V_H_ D115/V_L_ Y48 N332-glycan binding groove to the glycan (Figure 2C, Supplemental Figure 7A). Next, encounter states 2 and 3 resulted from coupling the N332-glycan to the antibody with various degrees of rotation about the glycan base (Supplemental Figure 7). These states accessed the bound conformation through a flexible pre-bound state, Encounter State 4 (Figure 2A, Supplemental Figure 7). In the MSM, access to the bound state depended on the coupling of the N301-glycan with the light chain and movement of the V1 loop that we previously defined using cryo-EM structure determination (Figure 2A, Supplemental Figure 7B and C) (*8*). These results showed that a series of encounter states that range from being distant to the target epitope to those that are closer to it enables access to the previously observed DH270.6 bound conformation. (*8, 33*).

We next designed Envs with faster DH270 lineage antibody association rates based on this pathway. A dominant pathway feature was the importance of early coupling of the N332-glycan D1 arm to the V_H_ D115/V_L_ Y48 groove (Figure 2C left, Encounter State 1). The heavy chain R98T mutation eliminated a salt bridge with D115, allowing D115 to form closer interactions with the N332-glycan, and L48Y added additional hydrogen bonding contacts (*25*). We hypothesized that enhancing access to this state would increase the antibody association rate specifically through interactions with these two residues and would thus yield a strong positive association rate gradient. This would be accomplished by improving access of the V_H_ D115/V_L_ Y48 groove to the N332-glycan and by increasing the probability that the encounter state will reach the pre-bound state.

Encounter State 1 stood out as a useful target due to its presentation of this groove to the N332-glycan. The N332-glycan acted as a tether for the antibody in the MSM, allowing the antibody to rotate into the bound state. Increasing the likelihood of N332-glycan capture early in the encounter process should, therefore, enhance the association rate.

Access to Encounter State 1 was hindered by an Env glycan at position N408 (Figure 2C). We thus eliminated the N-linked glycosylation site via an N408K mutation in addition to improving Encounter State 1 heavy chain contacts via Env mutations T290K and E293T, here referred to as the DH270 Encounter design 1 (DE1). We measured binding of DE1 to the V_H_ R98T/V_L_ L48Y containing I3.6 intermediate and other clonal lineage antibodies using surface plasmon resonance (SPR; Supplemental Figures 8 to 10, Supplemental Table 1). We found that the N408K mutation resulted in a doubling of the antibody-Env association rate, consistent with the predicted association pathway (Figure 2D, Supplemental Figure 8, Supplemental Table 1). Our MD simulation results showed that Encounter State 1 transitioned with the highest probability to the pre-bound state, Encounter State 4, through rotation about the N332-glycan, which occurred by visiting Encounter State 2. This transition was hindered by the close coupling of an Env glycan at position N442 to the antibody light chain (Figure 2C middle). The N442-glycan interaction with the light chain mirrored that of the N301-glycan in the bound conformation but limited interactions of the antibody with the conserved GDIK motif in the V3 loop of the Env (Supplemental Figure 7D). Thus, eliminating the N442-glycan was predicted to increase the probability that the N332-glycan rotation states, Encounter States 2 and 3, would convert to the pre-bound state. Mutation of N442 to alanine, referred to here as DE2, indeed resulted in a 5-fold improvement in the association rate of I3.6 (Figure 2D). The effect of N442-glycan removal was consistent with data showing that the presence of the N442-glycan correlated with early intermediate neutralization resistance (Supplemental Figure 11). In addition, we performed mammalian display-based selection for higher affinity Envs that also identified an N442D mutation (Supplemental Figure 12). Thus, these results showed that the MSM correctly identified encounter states and that the encounter-based design approach could be confirmed by mammalian display Env selection.

We next asked whether the MSM faithfully represented the transitions between encounter states. Two mutations are coupled in their effect on binding when the sum of the individual, single-mutant effects differs from the double-mutant effects relative to the unmutated construct. These double-mutant cycles (*34*) provide a powerful tool for probing transition intermediate states such as encounter states. The association pathway between the V4 proximal Encounter State 1 must transition through the N442-glycan hindered Encounter State 2 pathway to reach the bound state. Thus, the MSM indicated the N408K and N442A mutation effects on the association rate should be coupled. The sum of association rate improvements for the DE1 and DE2 designs was ∼7-fold. However, when introduced together, the association rate improved by ∼13-fold, nearly double that of the individual mutations. A double mutant cycle calculation based on transition state theory (*35*) for these two mutations yielded a ΔΔG of -2.7 kcal/mol, indicating the mutations were indeed coupled, thus confirming the MSM association transition pathway (Figure 2D).

We next asked what effect the V1-glycan deleted DT design mutations had on the association pathway. The CH848-d949 DT design showed a ∼3-fold enhanced association rate compared to the unmutated CH848-d949 SOSIP Env with a ∼2.5-fold improved dissociation rate for the I3.6 intermediate (Supplemental Figure 9, Supplemental Table 1). The enhanced association rate suggested that the elimination of the V1 glycans opened new encounter state transitions. We hypothesized that, if the elimination of the V1-glycans in the DT design opened a new, alternative association path, the effect of the DE1 and DE2 mutations would be reduced and these mutations would not show coupling. We, therefore, introduced the DE1 and DE2 mutations, independently and together in the DT design, referred to as DT-DE1-3, respectively, and measured their association rates. The association rate improvements for the I3.6 intermediate were reduced compared to the non-DT design. The DT-DE1 and DT-DE2 constructs showed ∼1.5 and 2.5-fold greater association rate, respectively, to I3.6 than the association rate for the DT construct. As predicted, these mutations showed little coupling when combined in the DT-DE3 design, with a transition state ΔΔG of -0.4 kcal/mol (Figure 2D, Supplemental Table 1). This loss in coupling confirmed that the elimination of the V1 glycans in CH848-d949 opened a new association pathway and showed that the DH270 lineage acquired the ability to recognize the HIV-1 Env through multiple encounter sites beginning at the I3.6 intermediate (Figure 2E).

### Key DH270 clonal lineage mutations selected in vivo by association pathway Env designs

BCR triggering strength is linearly correlated with association rates (*36*). Enhancing the association rate in immunogen design, therefore, might improve the selection of target bnAb residues. We next asked whether the CH848-d949 N408K/N442A design would yield such an association gradient by showing stronger binding to the I3.6 intermediate where the V_H_ R98T/V_L_ L48Y mutations occur, relative to the earlier I5.6 intermediate.

We measured the affinity and kinetics of binding of the designed immunogen to the DH270 UCA, as well as to DH270 intermediates I5.6 through I2 and the mature DH270.6 antibody. We compared the binding kinetics of the CH848-d949 N408K/N442A containing DE3 design to those of the unmutated CH848-d949 (Supplemental Figures 8 to 10, Supplemental Table 1). The DE3 design showed a four-fold greater association rate for the I3.6 intermediate compared to the earlier I5.6 intermediate (Figure 3A). The unmutated CH848-d949 trimer showed a twelve-fold association rate gradient between I5.6 to I3.6 intermediate antibodies, but substantially lower association rates than the DE3 design, at forty-five and thirteen-fold greater rates for the DE3 design, respectively (Figure 3A and B, Supplemental Figure 8). Thus, the DE3 design showed a more favorable affinity gradient for the selection of I3.6 mutations than the unmutated CH848-d949 trimer.

**Figure 3.**
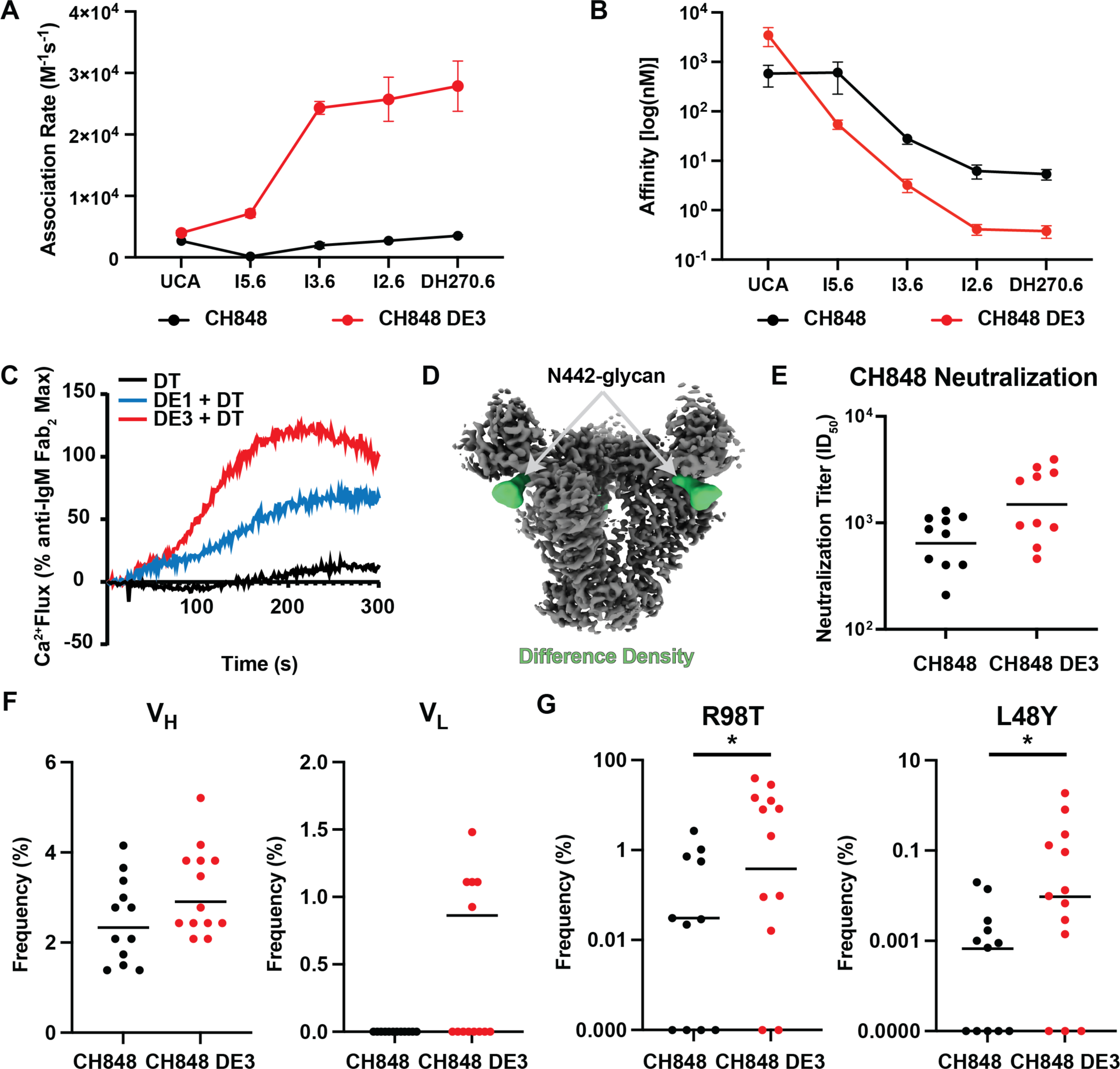
DH270.6 encounter state design improves the selection of target mutations. **A)** Association rates for the parent CH8483 Env SOSIP (black) and CH848 DE3 design (red) for the DH270 lineage UCA, intermediates, and DH270.6. **B)** Affinities for the parent CH8483 Env SOSIP (black) and CH848 DE3 design (red) for the DH270 lineage UCA, intermediates, and DH270.6. **C)** DH270.UCA Ramos B cell BCR triggering measured as a percentage of calcium flux relative to the maximum flux observed for an anti-IgM Fab_2_. **D)** Map of the DE3 design bound to DH270.6 (grey) and a difference map between the parent CH848 bound to DH270.6 and the DE3 design bound to DH270.6 highlighting the N442-glycan missing density (green). **E)** Serum neutralization of the parent CH848 pseudo-virus. Bars indicate the mean neutralization titer. Significance was calculated using a two-tailed, non-parametric Mann-Whitney test (* p < 0.5, N=10 each). **F)** Mutation frequencies for the human heavy (left) and light (right) chain genes in the immunized IA4 knock-in mice. Bars indicate the mean frequencies. **G)** Mutation frequencies for the IA4 key improbable mutations heavy chain R98T and light chain L48Y. Bars indicate the mean frequencies. Significance was calculated using a two-tailed, non-parametric Mann-Whitney test (* p < 0.5, N=11 for CH848-d949 and N=13 for DE3).

We then measured antigen-induced B cell calcium flux for each Env construct in a Ramos B cell line transfected with an IgM DH270 bnAb lineage unmutated common ancestor (UCA) antibody compared to the calcium flux induced by binding of an anti-IgM Fab_2_ antibody. Though the affinity for the DE1+DT design was slightly lower for DH270.UCA compared to the DT design, at 607 and 474 nanomolar, respectively, DE1+DT showed a ∼5-fold enhanced maximum calcium flux over the DT (Figure 3C, Supplemental Figure 8A, Supplemental Table 1). These data were consistent with the doubling of the association rate for DE1+DT compared to DT despite the faster DE1+DT dissociation rate. Similarly, DE3+DT had an association rate ∼3-fold faster than DT but with a ∼3-fold faster dissociation rate, giving rise to a similar affinity to DT (Supplemental Figure 8A, Supplemental Table 1). Thus, the DE3+DT design nevertheless showed over an order of magnitude higher calcium flux, indicating the enhanced DE1 and DE3 Env design association rates improved BCR triggering (Figure 3C).

We next tested whether differences in association and dissociation rates could be related to unintended Env structural changes caused by the designed mutations. We obtained a cryo-EM structure of the CH848-d949 and DE3 SOSIP Env bound to DH270.6 at 3.6 Angstroms and compared it to a previously determined CH848-d949 Env bound to DH270.6 (*33*), and found that the only discernable difference between the structures was the absence of density for the N442-glycan in the DE3 structure (Figure 3D, Supplemental Figure 13). We then compared the cryoEM structures of these Envs bound to the CD4 binding site VRC01 bnAb to examine the unliganded V3-glycan bnAb epitope structure. Differences in the structures were only visible at the N442-glycan position, where the DE3 design lacked density, and the Fab constant region, which showed a shifted elbow angle (Supplemental Figure 14). Thus, the design-enhanced kinetics are attributable to the encounter states observed in the MSM.

We next asked whether the improved association and affinity gradients would enhance selection of key V_H_ R98T/V_L_ L48Y I3.6 mutations <u>in vivo</u> in DH270 bnAb intermediate antibody knock-in mice. We immunized fourteen IA4 intermediate knock-in mice with either the CH848-d949 10.17 construct or the DE3 design. IA4 is a mutated version of I5.6 containing V_H_ S60Y and V_L_ V100I, neither of which is essential for neutralization breadth or potency (*15, 26*). Mice immunized with the DE3 design displayed an elevated mean titer of CH848-d949 pseudovirus neutralization compared to mice immunized with unmutated CH848-d949 Env and showed greater neutralization breadth and potency for several heterologous HIV-1 viruses (Figure 3E Supplemental Figure 15). Trends in elevated mean values for the DE3 design for total V_H_ and V_L_ mutations after vaccination were seen (Figure 3F). Our primary immunogen design goal was to specifically select for the improbable V_H_ R98T and V_L_ L48Y mutations that occur in the I3.6 intermediate. Next-generation sequencing of DH270 B cells in the mice immunized with DE3 induced significantly enhanced selection of the target V_H_ R98T and V_L_ L48Y mutations (Figure 3G). To assess whether the DE3 design selected for other known improbable mutations, we examined the frequency of the I2 intermediate mutation G110Y. This mutation was not involved in the relevant DE3 encounter states and did not show significant differences between the groups. (Supplemental Figure 15B). These results indicated that the D3 immunogen precisely selected the targeted neutralization-critical R98T and L48Y mutations, as targeted by our association path-based design strategy. It was important to determine if both mutations could be selected together in the same antibody. From a mouse immunized with DE3 that showed considerable neutralization breadth (Supplemental Figure 15A), we isolated a total of 46 DH270 bnAb B cell lineage memory B cell receptors. Of these, 10 had both V_H_ R98T/V_L_ L48Y mutations, and 9 contained R98T only (Supplemental Figure 15C). These results showed that the association pathway-based design selected for specific mutations targeted in the DH270 bnAb intermediate B cells by vaccination.

### Association pathway-designed immunogens for bnAb immunogens for the CD4 binding site

To determine the ability of association pathway immunogen design to be effective for targeting Env bnAb sites other than the V3-glycan epitope, we designed boosting immunogens for a CD4 binding site (CD4bs) bnAb B cell lineage. The most potent CD4bs bnAbs attain their neutralization breadth by mimicking CD4 interactions with Env. In contrast to V3-glycan bnAbs, CD4 mimicking bnAbs, are genetically restricted to two VH genes, VH1-2*02 or VH1-46 (*37–39*). Both show structural mimicry of the CD4 receptor interaction with the Env trimer (*20, 40*). Antibodies of the VH1-2*02-class use a rare five aa Light Chain Complementarity Determining Region 3 (LCDR3) and contain either deletions or multiple glycine mutations in the LCDR1 to accommodate the proximal gp120 loop D and N276 glycan (*40*). The VH1-46-class of bnAbs is an attractive target for HIV-1 vaccine design because neither deletions in the LCDR1 nor atypically short LCDR3s are required for their neutralization breadth, and a high-affinity germline targeting Env has been designed for the unmutated common ancestor (UCA)_of the VH1-46 prototype B cell lineage, CH235 (*41*). This immunogen selected for a key improbable mutation, K19T, in a CH235 UCA knock-in mouse model (*8*), and elicited CD4 mimicking serum-neutralizing CD4bs antibody precursors derived from rhesus VH genes orthologous to human VH1-46 (*42*). Here, we focused on the first two intermediates, the early I60 and the next intermediate antibody, I59, and two later intermediate antibodies that displayed substantial gains in neutralization breadth, I39 and I35 (Figure 4A) (*20, 42*).

**Figure 4.**
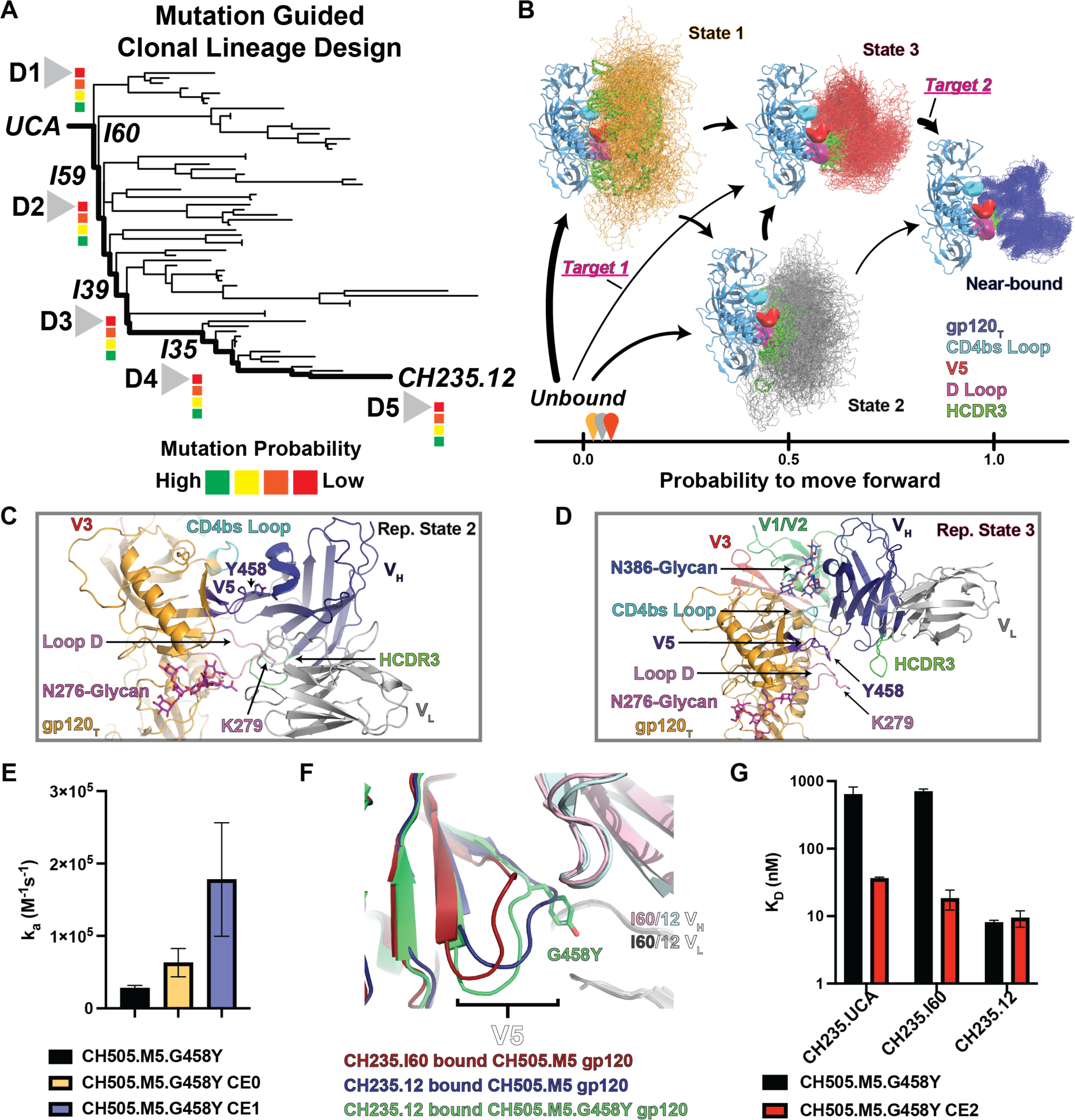
The association pathway for the CH235.12 Fv identifies design targets. **A)** Graphical representation of lineage-based design depicting a simplified phylogenetic tree leading from the inferred UCA and intermediates to the mature CH235.12 bnAb. Design targets D1-D4, each targeting low probability mutations (red), that are key for neutralization breadth. ***B*)** Association pathway for CH235.12 determined from an MSM prepared from the simulated set. Important regions of the gp120_T_ (blue) include the CD4bs loop (cyan), the V5 loop (red), the loop D (magenta), and the antibody HCDR3 (bright green). A single gp120 state is used throughout, with 50 representative samples of the antibody shown for each state (ensemble of line structures). Arrows show transition paths with sizes adjusted to indicate the relative flux through each transition. Structures are positioned based on the probability that each state will move forward to the bound state along the x-axis. Transitions targeted for design are highlighted (pink). **C)** Representative state from encounter State 2 highlighting the position of the N276-glycan and contact of the HCDR3 with loop D. **D)** Representative state from encounter State 3 highlighting the position of the N386-glycan and contact of the HCDR2 proximal sheet residues with the CD4bs loop. **E)** Mature CH235.12 association rates for the CH505.M5.G458Y parent and the CH505.M5.G458Y CE0 and CE1 designs. Error bars are standard errors from three replicate measures. **F)** Aligned structures of the CH505.M5 trimer bound to the early intermediate CH235.I60 Fab or mature bnAb CH235.12 Fab bound to either CH505.M5 or CH505.M5.G458Y highlighting the difference in the V5 loop. **G)** CH235 UCA, intermediate I60, and mature CH235.12 Fab affinities for the CH505.M5.G458Y parent and the CH505.M5.G458Y CE2 design. Error bars are standard errors from three replicate measures.

The MD system was prepared similarly to the CH848-DH270.6 system with a truncated CH505TF gp120 and a CH235.12 Fv positioned unbound near the CD4bs and with the antibody HCDR3 oriented toward a gp120_T_ (Supplemental Figure 16A). A total of 27 adaptive iterations were conducted, yielding 6,597 simulations for a total accumulated simulation time of 1.6 ms, including 34 transitions to a near-bound state. A common transition path involved coupling of the Env CD4-binding site loop with a β-sheet region adjacent to the antibody Heavy Chain Complementarity Determining Region 2 (HCDR2), followed by a rotation of the antibody that brought the light chain close to its final point of contact with the Env loop D (Supplemental Figures 16C to E). Unlike DH270.6, the fully matured CH235.12 antibody did not transition beyond a near-bound state, here defined as a bound state RMSD below 4 Å but without maintaining specific V_L_ and HCDR3 contacts (Supplemental Figure 16D). This appeared to be due to steric hindrance caused by flexible epitope loops that prevented formation of specific bound state contacts. This near-bound state nevertheless likely represents the final stage of the association process where these contacts are formed. We therefore used this state as the model endpoint to interrogate more distant encounter states. The association pathway determined from a MSM of these simulations revealed three encounter intermediate states, States 1-3, that all led to the near-bound state (Figure 4B, Supplemental Figures 17 to 19). Like the DH270.6 pre-bound state, the overall near-bound conformation was similar to the bound conformation but displayed considerable flexibility (Figure 4B, Supplemental Figure 17C). The dominant pathway led from diverse structural states surrounding the CD4bs, with the HCDR3 orientated toward the g120_T_ (Figure 4B). This state was connected to two distinct encounter states: one in which the HCDR3 interacted with V5 and loop D (Figure 4C) and the other state was with an HCDR2-adjacent V_H_ contact with the CD4bs loop (Figure 4D). Each encounter state had direct access to the near-bound state (Figure 4B).

In the context of the Env trimer, several conformations in Encounter States 1 and 2 would be partially occluded by the adjacent gp120 protomers, whereas Encounter State 3 is predicted to not be occluded by adjacent protomers (Supplemental Figure 20). We, therefore, selected Encounter State 3 for further design consideration since our vaccine Env immunogen platform is a trimer. Encounter State 3 is stabilized by polar contacts between CH235.12 V_H_ R56 and N54 sidechains and mainchain gp120 residues 364 to 366 of the CD4bs as well as residues 469 to 471. These contacts are similar to those found in a crystal structure of CH235.12 bound to the clade A/E 93TH057 Env gp120 (*20*) and act to anchor the antibody to the Env. These residues are at the periphery of the paratope, creating a hinge-like anchor about which the antibody can rotate into the bound configuration. These residues are G54 and S56 in the CH235.UCA and would provide limited stabilization of this encounter. Consistent with an important role for the changes observed in the mature CH235.12, we have shown mutations acquired at these residues enhance Env binding breadth when added to the CH235.UCA (*20*). Their appearance in an early encounter state here suggests they play important roles in both reaching and maintaining the bound state and are, therefore, important targets for design.

As in the DH270.6 pathway design, we focused on opening this state to collision and enhancing passage from this state to the near-bound state. Two glycans distant from the epitope limited access to the State 3 conformation - one was positioned on the V1/V2 domain at position N197, and the other on the outer domain at N386. Deletion of the N197-glycan was previously shown to enhance virus neutralization by CD4bs targeting bnAbs (*43*). We, therefore first examined an N-linked glycosylation site deletion at N197 (N197D) (*42*). The results showed a nearly 6-fold association rate enhancement for the CH235 UCA and a nearly ∼9-fold reduced association rate for the first intermediate, I60 (Supplemental Figures 23 to 25 and 26A, Supplemental Table 2). However, the following intermediate, I59, and later intermediates, I39 and I35, did not show association rate differences (Supplemental Figures 23 to 25 and 26A, Supplemental Table 2). The I39 and I35 intermediates showed reduced dissociation rates at 2- and 3-fold, respectively (Supplemental Figures 23 to 25 and 26A, Supplemental Table 2). The mature CH235.12 bnAb showed a ∼4-fold enhanced association rate and no change in the dissociation rate (Supplemental Figures 23 to 25 and 26A, Supplemental Table 2). The enhanced association rate for CH235.12 was consistent with the CH235.12 association pathway, which was hindered by the presence of the N197-glycan. The substantial differential in the N197D association and dissociation rates between the UCA, intermediates, and mature antibodies suggest a complex interplay between the effects of differing antibody mutations as the CH235 VH1-46 CD4bs B cell lineage develops.

Examination of a representative set of State 3 structures suggested that replacing the N386-glycan with an arginine could aid in attracting this state through interaction with CH235.12 F_v_ aspartate and glutamate residues (Supplemental Figure 21B). Consistent with Encounter State 3 structural observations, an N386A mutation, referred to here as CH235 Encounter design 0 (CE0), enhanced the CH235.12 Fab association rate by ∼2 fold and a stabilized form of an N386R mutation, referred to here as CE1, which contains the F14 and 2P stabilization mutations (*44, 45*), enhanced the CH235.12 Fab by ∼6 fold (Figure 4E, Supplemental Figures 23-25 and 26B, Supplemental Table 2). The non-stabilized CE1 design showed only ∼2-fold enhanced association rate, suggesting stabilization affected the rate (Supplemental Table 2). We therefore measured affinities for the F14+2P stabilized, unmutated CH505.M5.G458Y trimer. No changes in CH235.12 association rate were observed for this construct, indicating the stabilization alone did not yield the CE1 association rate.

These results show that the N386R and stabilization mutations were coupled in their effect on the CH235.12 association rate (Supplemental Table 2). Substantial impacts on the association rate for the N386A mutation were not observed for the CH235 UCA or intermediates I60 and I59. Modest enhancements began at I39 and I35, suggesting mutations acquired late in CH235 maturation permitted direct use of the N386-glycan occluded encounter site (Supplemental Figures 23 to 25 and 26B, Supplemental Table 2). Dissociation rates were largely unaffected by the N386 PNGS removal in the UCA, I60, I59, and I39 but showed a two-to four-fold dissociation rate increase, eliminating affinity gains (Supplemental Figures 23 to 25 and 26B, Supplemental Table 2). The N386-glycan site is adjacent to the CH235 antibody binding CD4bs loop interaction interface. As glycans are known to stabilize the protein fold (*46, 47*), eliminating this glycan may increase CD4bs loop dynamics and thereby impact bound state stability. The increase in the association rates for CH235.12 with N197 and N386 PNGS substitutions is consistent with the simulation-based association pathway that shows these glycans block encounter formation. However, the variability in PNGS removal impacts on the antibody lineage member kinetic rates suggests that these pathways may change substantially as the lineage develops.

We next examined gp120 dynamics in Encounter State 3 and the near-bound state. The CH235.12 antibody contacts the gp120 CD4bs loop, the V5 loop, and loop D in the bound state. Of these, the most conformationally variable loop was V5 (Supplemental Figure 21C). To better understand the impacts of this flexibility on antibody binding, we determined cryo-EM structures of our previous CH505 design that contained an N279K mutation (M5) in loop D bound to the early CH235 intermediate I60 and the mature CH235.12 bnAb in addition to a CH505.M5 design containing a G458Y (CH505.M5.G458Y) mutation in V5 bound to CH235.12 (Supplemental Figure 22). Examination of the V5 loop in these structures revealed marked differences in its conformation (Figure 4F). In the I60 bound CH505.M5, the V5 loop between residues 458 to 461 is positioned distant from the I60 light chain, showing limited interaction with the heavy chain. In the CH235.12 bound CH505.M5 structure, residues G458 and G459 rest in a groove between the V_H_ and V_L_, packing against the V_L_ W94 sidechain. This is permitted in the CH235.12 bound structure due to the I39 V_H_ W47L mutation, which allows the W94 rotamer to shift by ∼90°, creating the groove for V5 interaction. The introduction of I39 W47L mutation along with I59 S56R in the CH235 UCA and post I39 CH235 G54W was previously shown to increase recognition from three to five of fifteen heterologous HIV-1 Envs from multiple clades (*20*), suggesting this shift in V5 interaction plays an important role in neutralization breadth. The CH235.12 bound CH505.M5.G458Y V5 loop is positioned intermediate to these two V5 positions, inserting the aromatic tyrosine sidechain into the V_H_/V_L_ groove. These results suggest later intermediates can effectively manage V5 conformational variability.

We then asked whether limiting V5 loop flexibility in a CH505.M5.G458Y construct via a disulfide staple at the base of the V5 loop between residues 457 and 465, referred to here as CE2, could enhance early CH235 clonal lineage antibody affinities. A cryo-EM-derived structure of the CE2 design confirmed disulfide bond formation (Supplemental Figure 27). The V5-loop between residues 458 and 464 was poorly resolved in the cryo-EM map, suggesting these residues are conformationally dynamic.

We next compared CH235 clonal lineage kinetics between the parent CH505.M5.G458Y construct and the CE2 design. The UCA showed a ∼14-fold improvement in the dissociation rate with the CE2 design, with no change in the association rate. (Figure 4G, Supplemental Figures 23 to 25 and 26C, Supplemental Table 2). The early I60 intermediate showed a ∼2-fold enhanced association rate with a ∼20-fold improvement in the dissociation rate (Figure 4G, Supplemental Figures 23 to 25 and 26C, Supplemental Table 2). While these early-stage CH235 antibodies showed substantial dissociation rate improvements in the CE2 construct, the I59, I39, I35, and mature CH235.12 antibodies did not show dissociation rate improvements, with ∼2-fold association rate improvements for each (Figure 4G, Supplemental Figures 23 to 25 and 26C, Supplemental Table 2). The elimination of these dissociation rate gains beginning at the I59 intermediate suggests mutations occurring at I59 allow the antibody to manage V5 loop flexibility effectively. These include V_H_ S56R and D95N and V_L_ S30R. The V5-loop varies significantly between differing variants and clades, acquiring mutations, insertions, and deletions. While the I59 mutations outweighed the effect of the V5-staple before the acquisition of W47L this later mutation may nevertheless be essential for Env recognition breadth. Alternatively, the D457C mutation may affect the extensive, water-mediated interactions that couple the antibody R56 residue with the Env D457 (*20*), thus limiting affinity gains introduced by the V5-staple. These results show that antibody mutation effects can couple with Env residues positively and negatively and that the acquisition of certain antibody mutations can influence affinity gains or losses for additional mutations.

### An improved immunogen for selection of key functional improbable mutations in the CH235 bnAb B cell lineage

The CH235.12 N54 and R56 sidechains played an essential role in Encounter State 3, formed important contacts in the bound state (*20*), and enhanced heterologous Env binding (*37*), and therefore represent important immunogen design targets. Our previous designs included a CH235 UCA affinity enhancing loop D N279K mutation in the Env loop D. In the HIV-1 CH505 person living with HIV-1, the K279 residue was observed in Env sequences between days 22 and 93, suggesting that early CH235 antibody maturation occurred in the presence of N279 (*20*). Between days 93 and 205 following HIV-1 transmission, the N279 residue was dominant. By day 365, only D279 was present (*20*). Considering the marked affinity enhancement observed for CH235 binding to the K279 variant and the short time window over which it was present during infection, we asked what effect K279 had on the CH235 interaction. A previously determined CH235 bnAb UCA structure (*41*) and our structures of the I60 and mature CH235.12 bnAbs bound CH505 trimers here show that the K279 side chain does not interact with the antibody (Figure 5A, Supplemental Figure 28). It was, therefore, not clear how this residue facilitated enhanced affinity. Examination of previously published x-ray crystal structures of HIV-1 isolate 93TH057 gp120 cores, either unliganded or in complex with CH235.12, revealed that the loop D residue L277, which was buried in the unliganded Env, was bound to CH235.12 in the antibody-bound complex, demonstrating a conformational switch of the loop between its unliganded and antibody-bound states (Figure 5B) (*20, 48*). We hypothesized that the N279K mutation enhanced loop D rearrangements by increasing the likelihood that L277 moves to its extracted state, thereby increasing the association rate by eliminating a bottleneck to reaching the bound state. We, therefore, measured the binding of antibodies CH235 UCA, I60, I59, and CH235.12 with an engineered immunogen, CE3, that incorporated the V5 disulfide and the N386R substitution with or without the N279K substitution (Supplemental Table 2). Both designs with or without N279K showed substantially higher affinity for CH235 clonal lineage antibody binding than the previous design (Supplementary Table 2) (*8*). The association rate of the CH235 UCA was over an order of magnitude greater for the K279 design than the N279 design, and the dissociation rate for the K279 design was ∼3 fold lower than that for the N279 CE3 design (Figure 5C, Supplemental Table 2). The larger magnitude effect on the association rate between the N279 and K279 constructs is consistent with the K279 residue predominantly influencing association as predicted by the MSM.

**Figure 5.**
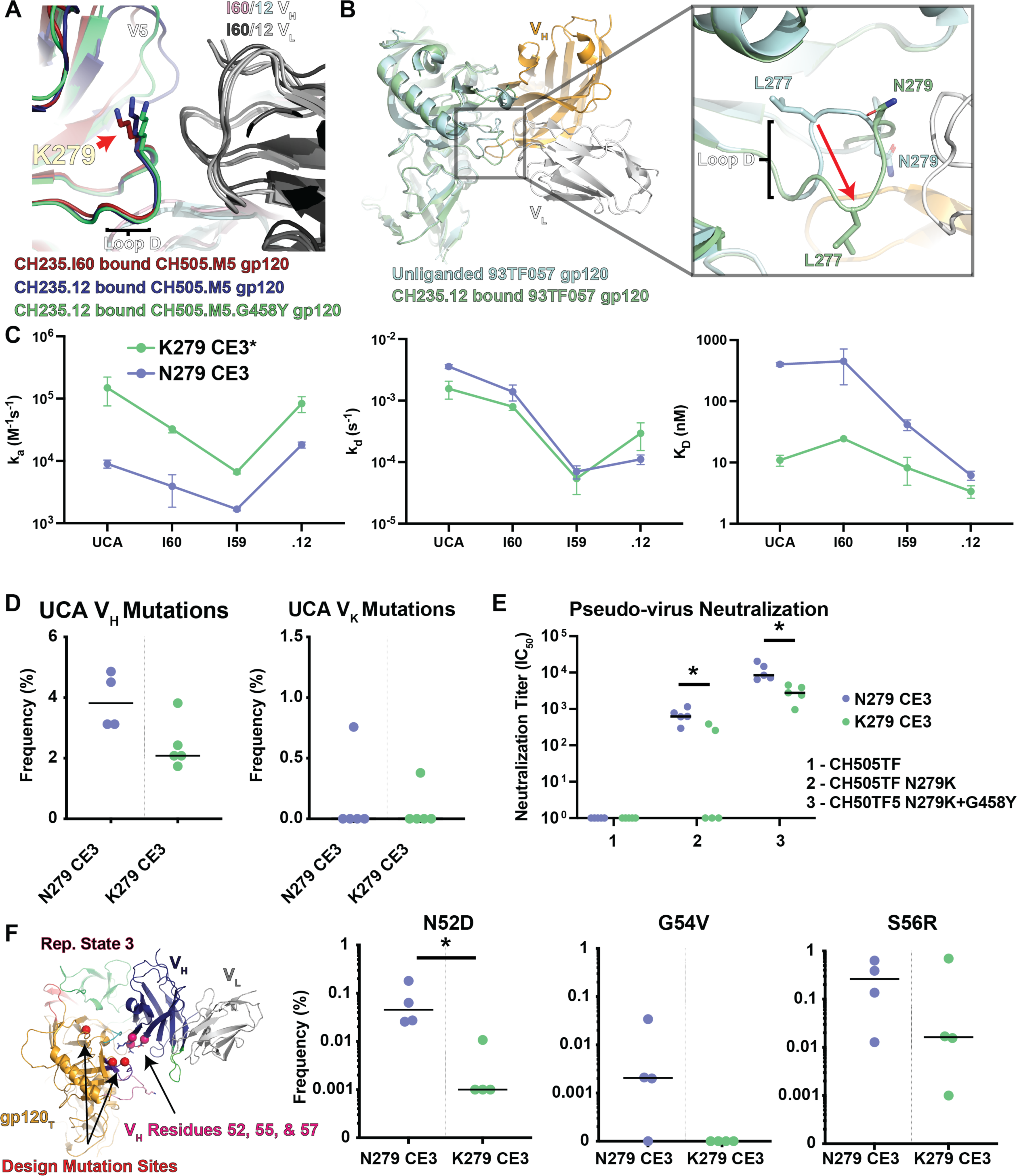
CH235.12 encounter state design improves the selection of target mutations. **A)** Aligned structures of the CH505.M5.G458Y trimer bound to either early intermediate CH235.I60 Fab (dark blue gp120, purple V_H_, dark grey V_L_) or mature bnAb CH235.12 Fab (dark red gp120, green V_H_, light grey V_L_) highlighting the position of loop D residue K279. **B)** (left) Aligned gp120 cores unliganded (light blue; PDB ID 3TGT) and bound (light green; PDB ID 5F96) to CH235.12 (V_H_ orange, V_L_ light grey). (right) View of loop D in the unliganded and CH235.12 bound state showing displacement of residue L277. **C)** Kinetics (left, ka, and middle, kd) and affinity (right) for the CH505.M5.G458Y CE3 design (N386R+V5 disulfide+N197D) with either K279 or N279. **D)** Mutations frequencies for human heavy (left) and light (right) chain genes in CH235 UCA knock-in mice. **E)** CH235 UCA knock-in mouse serum neutralization titers for CH505 isolate-derived pseudo-viruses. Significance was calculated using a two-tailed, non-parametric Mann-Whitney test (* p < 0.5, N=4 for each group). **F)** (left) Representative structure of encounter State 3 highlighting the position of the design mutations (red spheres) and the encounter state contacts (pink spheres) in the CH235.12 HCDR2 proximal sheet. (right) mutation frequencies for key I60 mutations from immunized CH235 UCA knock-in mice. Significance was calculated using a two-tailed, non-parametric Mann-Whitney test (* p < 0.5, N=4 for each group).

We next asked how these differences in rate affect the kinetic rate and affinity gradients. Faster association rates (i.e., a positive association rate gradient) and slower dissociation rate (i.e., a negative dissociation rate gradient) lead to a decrease in the dissociation constant (KD), signifying improved binding at each stage of antibody development. The N279 and the K279 CE3 designs displayed negative association and dissociation gradients from the UCA to the I59 intermediate with positive gradients between I59 and the mature CH235.12 (Figure 5C, Supplementary Table 2). This resulted in little difference in affinity between the UCA and first intermediate, I60, for the N279 CE3 design and a positive UCA to I60 affinity gradient for the K279 design (Figure 5C). The affinity gradients were negative from I60 to I59 and I59 to CH235.12, showing that the binding improved. The K279 CE3 design antibody interactions were nevertheless substantially stronger than the N279 design.

Despite the markedly greater affinity of the K279 CE3 design for the CH235 lineage antibodies, we hypothesized that the N279 CE3 design would more effectively select for mutations at Encounter State 3 antibody anchor positions 54 and 56 in a CH235 UCA knock-in mouse model. Specifically, anchor residues at these sites would increase the encounter state lifetime, which would allow the antibody more time for loop D rearrangements. With the N279K mutation, this delay time would be unnecessary and would, therefore, not cause the selection of these encounter state residues.

Thus, we immunized CH235 UCA knock-in mice with either the N279 CE3 or the K279 CE3 Env designs. The median mutation frequency in the human heavy chain was higher for the N279 CE3 design immunized mice compared to the K279 CE3 immunized mice (Figure 5D). We first compared serum pseudovirus neutralization for CH505TF, CH505TF with the N279K mutation, and CH505TF with both the N279K and G458Y mutations. We did not observe either group’s neutralization of the CH505TF pseudovirus (Figure 5E). However, both mutant pseudoviruses were neutralized by sera from all mice immunized with the N279 CE3 design (Figure 5E). All sera from the K279 CE3 design immunized mice neutralized the CH505TF N279K and G458Y-mutated pseudovirus, while only two of five sera neutralized the N279K-mutated CH505 pseudo-virus (Figure 5E). Mean neutralization titers for both mutated pseudoviruses were greater for the N279-immunized mouse sera. We, therefore, asked whether specific residue selection frequencies in the mice could explain this difference in neutralization. Consistent with predictions based on the CH235.12 association pathway, I60 intermediate V_H_ mutation G54V and I59 intermediate V_H_ mutation S56R showed elevated mean frequencies in the N279 CE3 immunized mice (Figure 5F). Additionally, an adjacent mutation, N52D, that stabilizes the LCDR2 loop showed a significant increase in frequency. The encounter State 3 proximal I60 intermediate V_H_ mutation K19T also showed an increased frequency in addition to a loop D proximal HCDR3 I60 intermediate V_H_ mutation A97V and a pre-I39 intermediate V_L_ framework mutation T71S (Supplemental Figures 29 and 30). These results showed that interactive antibody-antigen residue level mechanistic information and association-pathway-based design can be used to directly engineer immunogens targeted to disparate Env bnAb epitopes to select for antibody mutations at specific Env sites.

## Discussion

Here, we examined encounter complex formation based on molecular dynamics and Markov modeling to design boosting immunogens to select functional mutations of HIV-1 bnAbs. In doing so, we uncovered the 3-dimensional pathway these antibodies take to reach the bound or near-bound states on the Env, finding that epitope distant sites can be essential for antibody binding. This is consistent with findings from other methods such as the neutralization Env signature analysis, which identified non-contact Env signatures (both epitope proximal and distal) that show statistically robust associations with neutralization sensitivity to each class of bNAbs (*29*). While it remains to be systematically investigated, we speculate that such non-contact Env residues could impact bNAb binding or neutralization because they lie in the transient antibody footprints of one or more of the encounter states, and ultimately modulate the probability of reaching the bound state.

Combined with structural analysis of bnAb precursor-Env interactions, this pathway-based approach proved effective in 1) enhancing kinetic rates for target antibody mutations and 2) inducing, by vaccination, bnAb intermediate antibodies bearing the targeted mutations. For the V3-glycan lineage DH270, targeting the R98T and L48Y mutations showed mechanistically how these mutations aided in antibody binding and directly identified how the antigen could be modified to amplify their effects. This amplification specifically enhanced the selection of these mutations by vaccination, but not other known, critical mutations. For the CD4bs lineage CH235, we enhanced selection for mutations at the CD4bs proximal HCDR2 contacts, including S56R and G54V, by engineering a role for a critical encounter state. Thus, from these studies, a clear mechanism for the effects of previously identified mutations emerged that we leveraged to demonstrate that retaining certain bottlenecks while removing others can positively influence the selection of bnAb maturation mutations at specific antigenic sites via immunization. The association pathway immunogen design approach provided a robust path to designing immunogens that guide bnAb B cell lineage maturation.

The difference in affinity between the two CH235-targeted Env designs highlights a fundamental challenge in vaccine design. Simply removing hurdles to enhanced binding affinity does not guarantee that breadth-enhancing mutations will be selected in vaccination. For the DH270 lineage, we enhanced affinity directly through the targeted R98T and L48Y mutations while ensuring the gradient between the unmutated and mutated forms of the antibody was strong. For the CH235 lineage design, rather than focusing on high affinity binding, we optimized the immunogen to fully use the encounter state anchoring residue mutations we targeted. Each mutation’s role in the antibody reaching and retaining the bound state defined whether a particular bnAb mutation will be positively or negatively selected based on the properties of the antigen surface. This was exemplified in the CH235 G55V mutation, which is highly probable but was negatively selected in the K279 design. The association pathway indicated this occurred because it was rendered unimportant by bypassing its role in the association process. These results emphasize the necessity of understanding the role in Env binding for all antibody residues at the atomic scale.

The importance of encounter states in association pathways for interactive partners with non-diffusion-limited association rates is well documented (*49*). Typical barriers include conformational selection, induced fit, and steric factors that narrow the orientation space over which productive collisions can occur (*49*). In the case of antibody binding to HIV-1, each of these proved critical in both the DH270.6 and CH235.12 bnAb B cell lineage pathways. Consistent with the Env glycans acting as a shield, productive antibody-Env collisions occupied specific, limited orientations. While the encounter states displayed greater motion than bound states, movement occurred about specific hinge points with well-defined contact surfaces. Transitions from these collision-induced states then faced a conformational selection hurdle presented by glycan or loop motions that effectively acted as gates to the movement toward the bound state. In both DH270.6 and CH235.12, the final barrier to reaching the fully bound state involved induced fit, where a bound conformation required a conformational change in a proximal loop. For the V3-glycan bnAb, DH270.6, this involved displacement of the Env V1 loop by the DH270.6 V_H_ R57 residue. For CH235.12 CD4bs bnAb, the loop D must rearrange. This transition did not occur in our simulations, suggesting that this process is relatively slow, necessitating a long-lived encounter state to ensure that the antibody is not dislodged prior to loop D rearrangement. Defining this hierarchy allowed us to pinpoint Env mutations for association rate enhancement at two different levels of the association pathway. We first identified early encounter states involving target contacts and removed barriers to collision, namely, the removal of encounter proximal glycans. Removal of bound state proximal glycans has been tested previously, and, though holes in the glycan shield can encourage off-target responses(*50*), these immunogens have shown enhanced bnAb antibody affinities (*12, 51–55*). The previous approaches were, however, empirical, whereas the present approach is mechanistic and can guide complimentary antigen-antibody surface design at the deletion site for specific bnAb mutation selection. The second step in our approach involved removing barriers to transitions from collision-induced states to nearer-bound states. We showed that this approach can effectively guide the design of over an order of magnitude enhancement in antibody-antigen affinity for two distinct epitopes.

This high-precision approach has several limitations. Major hurdles exist in acquiring the large amount of data and information needed to facilitate this approach. Previous efforts provided a robust backdrop of clonal lineage ontogeny, virus neutralization, Env affinity, and high-resolution structure information, all key requisites to initiating a study like that presented here. It is also clear that specific bnAb mutation targets are needed.

HIV-1 bnAbs are extensively somatically mutated, often acquiring upwards of fifty mutations. Many of these mutations are unnecessary for neutralization breadth, as exemplified by minimally mutated bnAbs containing as few as twelve mutations while retaining breadth and potency (*13, 26, 56*). Few minimally mutated bnAbs have been identified, and the process for identification remains a time and resource-intensive task. Finally, the computational burden of the molecular simulation approach here is significant, requiring dedicated access to many machines running continuously for months at a time to predict a single antibody association path, creating a significant bottleneck to designing an effective HIV-1 vaccine. Each of these hurdles is substantial and the present study, shows the proof-of-concept for this undertaking.

With each year, new HIV-1 bnAbs are identified. Each of these represents a target for immunogen design, and they suggest that the antibody repertoire space amenable to broad neutralization is large. While no one precursor is guaranteed to be present in any particular individual, a vaccine designed to mature multiple bnAbs would greatly increase the probability of inducing one or more known bnAbs. Expanding this approach to known bnAbs, accelerating the throughput of bnAb identification, identifying the functional bnAb mutations, and defining mutation-targeted affinity gradients will be essential to design vaccines that can select for a polyclonal HIV-1 bnAb response. Machine learning models are being applied to solve complex biological problems such as mapping the mammalian brain connectome (*57*). It is evident from the studies presented here that it is a feasible goal to directly design Envs that target and select specific bnAb B cell lineage functional, improbable mutations. For a successful HIV-1 vaccine, it will be necessary to map bnAb-Env encounters at each step of B lineage development for multiple bnAb epitopes, and, as well, to accomplish this for multiple B cell lineages at each epitope. As with mapping the brain connectome, as more MSM data are acquired, developing machine learning models will be important to accelerate HIV-1 vaccine development.

## Methods

### Recombinant HIV-1 envelope SOSIPs, antibodies, and Fab production

The DS stabilized (*58*) CH848.10.17.d949 DT and DE1-3 envelope SOSIPs and the 4.1 (*59*), 2P (*45*), and F14 stabilized (*44*) CH505.M5.G458Y, CE0-3 SOSIPs were expressed in Freestyle^TM^ 293-F cells (ThermoFisher Cat No. R79007). The modified CH505 transmitted founder Env SOSIPs were chimeras containing the CH505 gp120 and BG505 gp41 domains. To express a sufficient amount of protein for immunization, the K279 construct required additional stabilization mutations, F14 and 2P. Before transfection, cells were diluted in Freestyle^TM^ 293 Expression Medium (Cat No. 12338018) to 1.25x10^6^ cells/mL at a volume of 950 mL. Plasmid DNA expressing the envelope SOSIP and furin were co-transfected at a 4:1 ratio (650 μg and 150 μg per transfection liter, respectively) and incubated with 293fectin^TM^ transfection reagent (ThermoFisher Cat No. 12347019) in Opti-MEM I Reduced Serum Medium (ThermoFisher Cat No. 31985062) to allow for complex formation. The diluted mixture was added to the cell culture which was incubated at 37°C, 9% CO_2_ on a shaker at 120 rpm for 6 days. On day 6 the cell supernatant was harvested by centrifuging the cell culture at 4000 xg for 45 minutes. The supernatant was filtered with a 0.45 μm PES filter and concentrated to approximately 100 mL using a Vivaflow^®^ 200 cross-flow cassette (Sartorius Cat No. VF20P2).

Envelope SOSIPs were purified using a PGT145 or PGT151 affinity chromatography equilibrated in 15 mM HEPES, and 150 mM NaCl (pH = 7.1). A PGT145 and PGT151 Ab affinity column was made by coupling mAbs to CNBr-activated Sepharose 4B (Cat No. 170430-01, GE Bio-Sciences) and packed into a Tricorn column (GE Healthcare). The supernatant was applied over the column at 2 mL/min using an AKTA go chromatography system (Cytiva) followed by three-column volume wash steps with 15 mM HEPES, and 150 mM NaCl. Protein was eluted off the column using 3 M MgCl_2_ and diluted in 15 mM HEPES, and 150 mM NaCl buffer. The protein sample was buffer exchanged into 15 mM HEPES, and 150 mM NaCl by ultrafiltration using a 100 kDa MWCO Amicon^®^ Ultra-15 Centrifugal Filter Unit (Millipore Aldrich Cat No. UFC9010) and concentrated to <0.5 mL for size exclusion chromatography. Size exclusion chromatography was performed using a Superose 6 10/300GL Column (Cytiva) on an AKTA go system in 15 mM HEPES, and 150 mM NaCl. Fractions containing trimeric SOSIP were collected. Lots produced were subjected to quality control including analytical SEC, SDS-PAGE, thermal shift analysis, biolayer interferometry (BLI), and negative stain electron microscopy (NSEM) to assure the presence of well-folded Env trimers.

### Thermal shift assay

Thermal shift assay was performed using Tycho NT.6 (NanoTemper Technologies). Envelope ectodomain SOSIPs were diluted (0.15LmgLml^−1^) in 15 mM HEPES buffer with 150 mM NaCl at pH 7.1. Intrinsic fluorescence was recorded at 330 nm and 350 nm while heating the sample from 35 to 95°C at a rate of 3°CLmin^−1^. The ratio of fluorescence (350/330 nm) and the inflection temperatures (Ti) were calculated by Tycho NT.6.

### Molecular Dynamics

Encounter state lifetimes can range from several to hundreds of microseconds. MD simulations at these timescales are accessible only by specialized computing hardware that is not widely available. This was accomplished by building MSMs from the aggregated simulations to identify movements toward transition states. The states are used in the next iteration. Since simulations rarely access transition states and typically return to metastable states when they approach transition states, launching simulations from near-transition states increases the likelihood of moving to and beyond a true transition state.

The system size determines the calculation rate for atomistic MD simulations. As the number of atoms in the simulation increases, the computing time needed to complete a simulation increases geometrically.

Therefore, it is essential to prepare the smallest, physically realistic system possible to achieve sufficiently robust information that is of biological relevance. We prepared systems of N- and C-terminal truncated gp120 domains extracted from Env SOSIP closed-state structures (PDB IDs 6UM6 and 6UDA for CH848.10.17.d949 and CH505TF, respectively). Antibody Fv structures were prepared from crystal structures (PDB IDs 5TQA and 5F96 for DH270.6 and CH235.12, respectively). Missing loops were added using MODELER (*60*), Unbound antibody Fvs were positioned near the relevant epitope in PyMol(*61*). The Fv-gp120_T_ system was glycosylated using the CHARMM-GUI Glycan Modeler (*62–67*) based Glycosite-predicted (*68*) glycosylation sites with either mannose-9 or mannose-5 glycans, for the CH848 and CH505 systems, respectively. The glycosylated system was immersed in a 150 Å octahedral water box, neutralized, and brought to a NaCl concentration of 150 mM. The Amber22 software package with pmemd CUDA (*69–71*) using the CHARMM36 force field (*72–75*) and TIP3P water model (*76*) was used for simulations. Electrostatic interactions were calculated with the Particle Mesh Ewald method (*77*) with a cutoff of 12 Å and a switching distance starting at 10 Å. The SHAKE algorithm (*78*) with hydrogen mass repartitioning (*79*) was used to constrain hydrogen atoms and allow for a 4 fs timestep. To equilibrate the systems, 10,000 steps of energy minimization were performed with 1000 steps of steepest descent minimization with 500 kcal/mol•Å restraints on protein followed by an unrestrained energy minimization consisting of 10,000 steps and 1000 steepest-descent steps. Minimization was followed by 20 ps of NVT heating with 10 kcal/mol•Å restraints applied to protein atoms and a subsequent 5 ns NPT equilibration with no restraints, with a constant temperature of 298.15 K maintained using a Langevin thermostat (*80*), and pressure of 1 atm was maintained with isotropic position scaling. A total of 200-400 250 ns unrestrained simulations in the NVT ensemble were performed to equilibrate the system and generate a wide ensemble of antibody orientations relative to the Env gp120. The cpptraj tool in AmberTools21 (*81*) was used for trajectory unwrapping.

### Adaptive Sampling

The last frame from each of the 200-400 250 ns was extracted for use in the first adaptive sampling routine epoch. Adaptive sampling was performed using the High-Throughput Molecular Dynamics (HTMD v. 1.24.2) package (*79*). Each subsequent epoch consisted of ∼200-400 independent simulations of 250 ns. At the completion of each epocj, simulations from each iteration were first projected using a distance-based metric between gp120 and Fv amino acid residues gp120 glycan atoms. This was followed by a TICA (*80*) projection using a lag time of 5 ns, retaining five dimensions. MSMs were then built using a lag time of 50 ns for the selection of new states for the next iteration. A total of 17 adaptive epochs were performed for the DH270.6-CH848-d949 and CH235.12-CH505 systems, yielding total simulation times of ∼2 ms and ∼1.1 ms for the CH848-d949 system and CH235.12-CH505 systems, respectively. Simulations were visualized in VMD and PyMol and selected using tools in the HTMD software.

### Markov state modeling

MSMs were prepared in HTMD using inverse exponential distances between amino acid residues and glycan units identified as contacts in relevant antibody-bound Env subunits (PDB IDs 6UM6 and 6UDA for DH270.6-CH848-d949 and CH235.12-CH505, respectively: HXB2 residues 135, 136, 137, 138, 139, 148, 149, 299, 300, 301, 321A, 322, 323, 324, 325, 326, 327, 328, 329, 330, 332, 415, and 417 and the N138, N156, N301, N332, and N442 glycans for CH848, DH270.6 V_H_ residues 33, 52, 55, 57, 59, 84, 101, 103, 104, 105, 106, 107, 108, 109, 110, 111, 113, and 115, and DH270.6 V_L_ residues 27, 32, 48, 51, 52, 57, 58, 93, 96, and 97 and HXB2 residues 280, 282, 365, 368, 429, 458, and 470 for CH505 TF, CH235.12 V_H_ residues 30, 55, 57, 62, 65, 72, 102, 104, 105, and 106 and CH235.12 V_L_ residues 92 and 94). MSMs were prepared in HTMD using a TICA lag time of 25 ns retaining five dimensions followed by K-means clustering using 500 cluster centers. The implied time scales (ITS) plots were used to select a lag time of 100 ns for MSM building. Models were coarse-grained via Perron cluster analysis (PCCA++) using 5 to 6 states and validated using the Chapman-Kolmogorov (CK) test. State statistics were collected for mean first passage times (MFPT), stationary distributions, and root-mean-square deviations (RMSDs) relative to known bound state structures for each respective system. The RMSD and contact metric means were model-weighted. Weighted state ensembles containing 50-250 structures were collected for visualization in VMD.

### Cryo-electron microscopy sample preparation, data collection, and processing

Purified Envelope SOSIP ectodomain preparations were prepared at concentrations of 4-5 mg/ml in 15 mM HEPES buffer with 150 mM NaCl at pH 7.1 and mixed with Fab at a 1:5 molar ratio. A total of 2.5 µl of the complex was deposited on a CF-1.2/1.3 grid that had been glow-discharged for 15 s in a PELCO easiGlow glow discharge cleaning system. After 30-s incubation in >95% humidity, the excess protein was blotted away for 2.5 s before being plunge-frozen in liquid ethane using a Leica EM GP2 plunge freezer (Leica Microsystems). Frozen grids were imaged in a Titan Krios (Thermo Fisher) equipped with a K3 detector (Gatan). Individual frames were aligned, dose-weighted, and CTF corrected, followed by particle picking, 2D classification, ab initio model generation, heterogeneous refinements, homogeneous 3D refinements, and local resolution calculations in cryoSPARC (Supp. Table 1) (113).

### Cryo-electron microscopy structure fitting and analysis

The antibody-bound SOSIP structures for DH270.6 and CH235 lineage antibodies (PDB IDs 6UM6 and 6UDA, respectively) structures were used to fit the cryo-EM maps in ChimeraX (114, 115). Mutations were made in PyMol (*82*). Coordinates were then fitted using Isolde (116) followed by iterative refinement using Phenix (*83*) real-space refinement. Structure and map analyses were performed using PyMol, Chimera (117), and ChimeraX (114, 115).

### Surface Plasmon Resonance

The SPR binding curves of the DH270 and CH235 lineage Fabs against CH848 SOSIPs and CH505M5 SOSIPs, respectively, were obtained using either a Biacore S200 or T200 instrument (Cytiva) in HBS-EP+ 1X running buffer (Cytiva). For the DH270 lineage Fabs (UCA3, I5.6, I3.6, I2.6 and DH270.6), PGT151 was immobilized onto a CM5 chip to approximately 10,000RU. The CH848 SOSIPs were injected at 5ul/min and captured to a level of 100-800RU with an average capture level of 450RU. Using the single cycle injection type, five sequential injections of the Fabs diluted from 25 to 1500nM were injected over the captured CH848 SOSIP Trimers at 50uL/min for 120s per concentration followed by a dissociation period of 600s. The Fabs were regenerated with a 12s pulse of Glycine pH2.0 at 50uL/min. Fab binding to a surface with negative control Flu antibody, Ab82, immobilized to a similar level as PGT151 as well as buffer binding were used for double reference subtraction to account for non-specific antibody binding and signal drift. Curve fitting analyses were performed using the 1:1 Langmuir model with a local Rmax. One interaction (DE1 with DH270UCA3) used the heterogeneous ligand model with the faster kinetics reported. The reported binding curves and kinetics for all DH270 Fabs are representative of at least one data set. For the CH235 lineage Fabs (UCA, I60, I59, I39, I35 and CH235.12), PGT151 was immobilized onto a CM5 chip to approximately 5,000-10,000RU. The CH505M5 SOSIPs were injected at 5ul/min and captured to a level of 100-800RU with an average capture level of 400RU. Using the single cycle injection type, five sequential injections of the Fabs diluted from 31.25-3000nM were injected over the captured CH505M5 SOSIP Trimers at 30uL/min for 120s per concentration followed by a dissociation period of 600s. The Fabs were regenerated with a 12s pulse of Glycine pH2.0 at 50uL/min. Fab binding to a surface with the negative control Flu antibody, Ab82, immobilized to a similar level as PGT151 as well as buffer binding were used for double reference subtraction to account for non-specific antibody binding and signal drift. Curve fitting analyses were performed using the 1:1 Langmuir model with a local Rmax. The reported binding curves and kinetics for all CH235 Fabs are representative of at least three data sets.

### B cell activation and Calcium Flux Measurements with Ramos cells

Calcium flux experiments were performed using the FlexStation 3 Microplate Reader (Molecular Devices) and Softmax Pro v7 software (Molecular Devices) in conjunction with the FLIPR Calcium 6 dye kit (Molecular Devices). On the day of the experiments, a cell count for the Ramos cells was performed with a Guava Muse Cell Analyzer (Luminex) to ensure cell viability was greater than 95% and to calculate the volume of cells needed for a concentration of 1x10^6^cells/mL. The appropriate volume of cells was then pelleted at 1500rpm for 5 minutes after which the supernatant was decanted and the cells were resuspended at a 2:1 ratio of RPMI media (Gibco) + FLIPR Calcium 6 dye (Molecular Devices). The cells were plated in a clear, U-bottom 96-well tissue culture plate (Costar) and incubated at 37°, 5% CO_2_ for 2h. Antigens were separately diluted down to a concentration of 500nM in 50uL of the 2:1 ratio of RPMI media (Gibco) + FLIPR Calcium 6 dye (Molecular Devices) and plated in a black, clear bottom 96-well plate. The final concentration of antigen would be 250nM based on the additional 50uL of cells added during the assay. A positive control stimulant, anti-human IgM F(ab’)_2_ (Jackson ImmunoResearch) was also included in the antigen plate. Using the FlexStation 3 multi-mode microplate reader (Molecular Devices), 50uL of the cells were added to 50uL of protein or Anti-human IgM F(ab’)_2_ diluted in RPMI/dye and continuously read for 5min. Calcium flux results were analyzed using Microsoft Excel and GraphPad Prism v9. The relative fluorescence of a blank well containing only the RPMI/dye mixture was used for background subtraction. Once subtracted, the antigen fluorescence was then normalized with respect to the maximum signal of the IgM control, and calcium flux values were presented as a percentage (% of Anti-hu IgM Fab2 Max). Calcium flux data are representative of at least 2 measurements for the DH270UCA3 cell line.

### Generation of the DH270 IA4 mice

The DH270 IA4 mice were generated by integrating pre-rearranged V(D)J exons of DH270 IA4 antibody heavy chain (HC) and light chain (LC) (Supplementary Table 3) into the mouse immunoglobulin heavy chain (IgH) J_H_ and kappa light chain (Igk) J_k_ loci respectively. The integration was performed in mouse embryonic stem (ES) cells. In the integration construct, the DH270 IA4 HC or LC V(D)J exons were preceded by mouse V_H_ promoter (for HC) or Vk promoter (for LC), which drive the expression of V(D)J exons in B cells. The expression cassette, which consists of the V promoter and V(D)J exon, were flanked by DNA fragments that were derived from the mouse J_H_ or Jk regions. Upon transfection of the construct into ES cells, homologous recombination mediates the integration of the DH270 IA4 V(D)J exons into the J_H_ or Jk loci. At the integration site, the DH270IA4 HC expression cassette replaces the whole mouse J_H_ region, from 877bp upstream of J_H_1 to 237bp downstream of J_H_4; the DH270 IA4 LC expression cassette replaces the entire mouse Jk region, from 114bp upstream of Jk1 to 286bp downstream of Jk4. ES clones with correct integration event were identified by Southern blotting. The neomycin resistance gene, which was used to select for stable integration of the construct, was subsequently removed by cre recombinases-mediated excision through flanking loxP sites; this step is necessary to prevent potential interference of local transcriptional regulation by the drug selection marker. The final ES clone was injected into Rag2 deficient blastocysts to generate chimeric mice. Because Rag2 is required for V(D)J recombination, all B and T cells in the chimeric mice are derived from injected ES cells, and the chimeric mice can therefore be used for analysis of lymphocyte function. The chimeric mice were subsequently bred with C57BL/6 mice for germline transmission. Since the original ES cell was derived from a F1 mouse that resulted from a cross between 129Sv and C57BL/6 mice, the mouse line is of mixed 129Sv and C57BL/6 genetic background.

### Knock-in Mouse Immunizations

The CH235 UCA KI mouse model has been described previously (*8*). 10-11 weeks old mice were used.

10 CH235 UCA KI mice were split into two groups (N = 5 each group), each group had at least one female mice. The synthetic Toll-like receptor 7/8 agonist 3M-052 absorbed to ALUM (3M-052-ALUM) was used as the adjuvant for the vaccine immunogens. Mouse vaccination studies were performed intramuscularly with 3M052-ALUM adjuvanted N279 CE3 design or the K279 CE3* design proteins. Vaccine immunogens were administered at 25mcg and formulated with 0.5mcg of adjuvant. Mice were immunized every two weeks for five times. Blood samples were collected 7 days prior to the first immunization (pre-bleed) and 7 days after each immunization. The DH270 IA4 KI mice are described above. 12-16 weeks old mice were used. 25 DH270 IA4 KI mice were split into two groups, each group had at least one female mice. The synthetic Toll-like receptor 7/8 agonist 3M-052 absorbed to ALUM (3M-052-ALUM) was used as the adjuvant for the vaccine immunogens. Mouse vaccination studies were performed intramuscularly with 3M052-ALUM adjuvanted CH848-d949 construct (N = 12) or the DE3 design (N = 13) proteins. Vaccine immunogens were administered at 17.5-25mcg and formulated with 0.5mcg of adjuvant. Mice were immunized every two weeks for six times.

Blood samples were collected 7 days prior to the first immunization (pre-bleed) and 7 days after each immunization. Necropsy was performed one week after the third immunization and blood, spleen, and lymph nodes were collected. All mice were cared for in a facility accredited by the Association for Assessment and Accreditation of Laboratory Animal Care International (AAALAC). All study protocol and all veterinarian procedures were approved by the Duke University Institutional Animal Care and Use Committee (IACUC).

### Antibody isolation and production

Single antigen-specific B cells were sorted as previously described (*8*). Briefly, inguinal and axillary lymph nodes were processed into single-cell suspensions and cryopreserved. Upon thawing, cells were counted then stained with optimal concentrations of the following fluorophore-antibody conjugations: BB700 anti-IgG1 (clone A85-1, BD Biosciences), BB700 anti-IgG2a (clone R2-40, BD Biosciences), BB700 anti-IgG3 (clone R40-82, BD Biosciences), PE anti-GL7 (clone GL7, Biolegend), PE-Cy7 anti-IgM (cloneR6-60.2, BD Biosciences), APC-R700 anti-CD19 (clone 1D3, BD Biosciences), BV510 anti-IgD (clone 11-26C.2a, BD Biosciences), BV605 anti-IgK (clone 187.1, BD Biosciences), and BV650 anti-B220 (clone RA3-6B2, BD Biosciences). Cells were also labeled with fluorophore-labeled CH848-d949 DT (10.17) in two colors (VB515 and Alexa Fluor 647). Dead cells were identified by labeling with LIVE/DEAD Fixable Near I/R stain (Thermo Fisher Scientific). Class-switched B cells were identified as single, viable lymphocytes that were B220+CD19+IgM-IgD-IgG+; antigen-specific B cells were identified as cells that co-labeled with CH848DT baits conjugated to VB515 and AlexaFluor647. Single cells were sorted on a BD FACS AriaIIu running DIVA version 8.0. Cells were sorted into cell lysis buffer and 5X first-strand synthesis buffer in individual wells of a 96-well PCR plate. Plates were frozen in a dry ice/ethanol bath immediately and stored at -80 °C until reverse transcription of RNA. Immunoglobulin genes were amplified from singly sorted antigen-specific B cells of DH270 IA4 knock-in mice by RT-PCR as previously described with some modifications (*8, 84, 85*). Briefly, immunoglobulin genes were reverse-transcribed with Superscript III (Thermo Fisher Scientific, <u>Waltham, MA</u>) using random hexamer oligonucleotides as primers. The complementary DNA was then used as template to perform nested PCR for amplification of antibody heavy and light chain genes using AmpliTaq Gold 360 (Thermo Fisher Scientific, <u>Waltham, MA</u>). In parallel, PCR reactions were done with mouse immunoglobulin-specific primers and DH270 variable region-specific primers. In one set of PCR reaction, mouse variable region and mouse constant region primers were used to amplify mouse heavy, kappa and lambda chain genes. In another set of PCR reaction, human variable region and mouse constant region primers were used to amplify heavy and lambda chain genes of the knock-in DH270 IA4 and its derivates. Positive PCR amplification of immunoglobulin genes was identified by gel electrophoresis. Positive PCR products were purified in Biomek FX Laboratory Automation Workstation (Beckman Coulter, Indianapolis, IN) and sequenced by Sanger sequencing. Contigs of the PCR amplicon sequence were made, and immunogenetics annotation was performed with software Cloanalyst (*86*) using both the human and mouse Ig gene libraries. Antibody genes were categorized as human or mouse based on the species of the V gene with the highest identity to the antibody sequence. The immunoglobulin gene amplicons were further used for protein expression by generating linear expression cassette of each gene. The linear expression cassettes were constructed by overlapping PCR to place the PCR-amplified VH and VK/L chain genes under the control of a CMV promoter along with heavy chain IgG1 constant region or light chain constant region and a BGH ploy A signal sequence. The linear expression cassettes of each paired heavy chain and light chain were then co-transfected into 293T cells. The supernatant containing recombinant antibodies were cleared of cells by centrifugation and used for binding assays. The genes of selected heavy chains were synthesized as IgG1 (GenScript). Kappa and lambda chains were synthesized similarly. Plasmids with the cloning of synthesized antibody genes were prepared using the Megaprep plasmid plus kit (Qiagen) for transient transfection. The paired heavy and light chain plasmid DNA were co-transfected into 293i cells using ExpiFectamine^™^ (Life Technologies, Carlsbad, CA) by following the manufacturer’s protocol. Antibody proteins were purified from the cell culture supernatant by protein A beads for binding, neutralization, and other immunological assays.

### Mammalian cell surface display of HIV-1 envelope

A site-saturation library was designed using CH848.3.D0949.10.17 gp140 with N133D and N138T substitutions (10.17DT) as a template (PMID: 31806786). N133D and N138T substitutions were added to enable HIV-1 bnAb DH270 inferred precursor binding (PMID: 31806786). The site-saturation envelope library was synthesized (GenScript) to contain envelopes with a single substitution. The substitutions introduced could be any of the other 19 possible amino acids at any one position within the gp120 subunit. The stabilized, cleaved HIV-1 envelope gp140 was fused to a C-terminal c-myc tag and HRV-3C cleavage site, followed by a platelet-derived growth factor receptor transmembrane region that ensured cell surface presentation of the envelope. The open reading frame encoding the envelope library was cloned into the mammalian expression vector VRC8400 CMV/R.

Sixteen micrograms of CMV/R vector carrying HIV-1 SOSIP library were co-transfected with 67 µg of carrier DNA (CMV/R vector) without the insert and 3.7 µg of plasmid coding Furin into Freestyle 293F cells using a 293fectin Transfection Kit (Thermo Scientific) according to manufacturer’s instruction. Cells were cultured for 48 hours with shaking at 120 rpm in an incubator filled with 8% CO2 at 37°C. Twenty million harvested transfected cells were stained with 500 µl of 100 µg/ml DH270 UCA and 10 µg/ml chicken anti-c-myc. DH270 UCA and anti-c-myc antibody binding were detected with 500 µl of 10 µg/ml goat-anti-human PE (Sigma) and 10 µg/ml goat anti-chicken Alexa 647 respectively. In separate tubes, two million cells were stained with 50 µl of the above antibodies for analytical flow cytometry. Gates for sorting were set such that cells that bound to DH270 with a higher fluorescence intensity than the unmutated envelope (prototype) were sorted. RNA was then isolated from sorted cells using PurelinkTM RNA micro kit (Invitrogen). RNA was subjected to reverse transcription primed by CMVR-SOSIP-rev (cacagc<u>agatct</u> tcatcagcggggctttttctg) using AccuScript Reverse Transcriptase (Agilent) to generate cDNA. Double-stranded DNA was amplified from cDNA with CMVR-SOSIP-fwd (gtcaccgtc<u>gtcgac</u>gccacc atggaaaccgatacactgct) and CMVR-SOSIP-rev using Herculase II Fusion DNA Polymerases (Agilent) and subcloned into CMVR vector by SalI/BglII digestion and ligation. The resulting constructs were transformed into electrocompetent DH10B using electroporation followed by culture in 500 mL of Luria broth. Plasmid was then extracted for another round of transfection and sorting. To get stronger binding envelopes, the selection stringency was increased by decreasing the concentration of DH270 UCA used to sort positive cells in each round. Sorted cells were sequenced by PacBio long-read sequencing.

PacBio Sequencing of library variants. RT-PCR products from the last cell sorting were purified using a gel-extraction kit (Qiaqen) and sequenced using PacBio long-read sequencing. The frequency of mutations at desired sites were calculated. DNA samples were quantified using Qubit 2.0 Fluorometer (Life Technologies, Carlsbad, CA, USA) and sample purity and integrity was checked using Nanodrop and Agilent TapeStation, respectively. DNA library preparations, sequencing reactions, and initial bioinformatics analysis were conducted at Azenta Life Sciences (Formerly Genewiz) (South Plainfield, NJ, USA). Following slow annealing, PacBio SMRTbell amplicon libraries for PacBio Sequel were constructed using SMRTbell Express Template Prep Kit 2.0 (PacBio, Menlo Park, CA, USA) using the manufacturer recommended protocol. The pooled library was bound to polymerase using the Sequel Binding Kit 3.0 (PacBio) and loaded onto PacBio Sequel using Sequel Sequencing Kit 3.0. Sequencing was performed on the required PacBio Sequel SMRT cells. The generated subreads were demultiplexed and clustered. Long Amplicon reads were obtained using PacBio laa pblaa v2.4.2 with default parameters. The generated subreads were demultiplexed and aligned to the reference sequence using the PacBio software pbmm2. Variant calling was performed using the PacBio software variantCaller arrow using default parameters. Variants that passed the default filters were parsed into a filtered vcf file for each sample. The substitutions observed in the most abundant variants were introduced into soluble HIV-1 envelope for characterization.

### HIV-1 neutralization assays

Neutralizing antibody assays were performed with HIV-1 Env-pseudotyped viruses and TZM-bl cells (NIH AIDS Research and Reference Reagent Program contributed by John Kappes and Xiaoyun Wu) as described previously(*87, 88*). Neutralization titers are the reciprocal sample dilution (for serum) or antibody concentration in µg/mL (for mAbs) at which relative luminescence units (RLU) were reduced by 80% or 50% (ID80/IC80 and ID50/IC50 respectively) compared to RLU in virus control wells after subtraction of background RLU in cell only control wells. Serum samples were heat-inactivated at 56 °C for 30 minutes prior to assay.

### Next Generation Sequencing of Mouse B Cell Receptor Repertoires

We performed next-generation sequencing (NGS) on mouse antibody heavy and light chain variable genes using an Illumina sequencing platform. RNA was purified from splenocytes using the RNeasy Mini Kit (Qiagen, Cat# 74104). Purified RNA was quantified via Nanodrop (Thermo Fisher Scientific) and used to generate Illumina-ready heavy and light chain sequencing libraries using the SMARTer Mouse BCR IgG H/K/L Profiling Kit (Takara, Cat# 634422). Briefly, 1 μg of total purified RNA from splenocytes was used for reverse transcription with Poly dT provided in the SMARTer Mouse BCR kit for cDNA synthesis. Heavy and light chain genes were then separately amplified using a 5’ RACE approach with reverse primers that anneal in the mouse IgG constant region for heavy chain genes and IgK for the light chain genes (SMARTer Mouse BCR IgG H/K/L Profiling Kit). The DH270 IA4 KI mouse model has the light chain gene knocked into the kappa locus, therefore kappa primers provided in the SMARTer Mouse BCR kit were used for light chain gene library preparation. 5 µl of cDNA was used for heavy and light chain gene amplification via two rounds of PCR (18 and 12 cycles per round). During the second round of PCR, Illumina adapters and indexes were added.

Illumina-ready sequencing libraries were then purified and size-selected by AMPure XP (Beckman Coulter, Cat# A63881) using kit recommendations. The heavy and light chain libraries per mouse were indexed separately, thus allowing us to deconvolute the mouse-specific sequences during analysis. Libraries were quantified using QuBit Fluorometer (Thermo Fisher). Mice were pooled by groups for sequencing on the Illumina MiSeq Reagent Kit v3 (600 cycle) (Illumina, Cat# MS-102-3003) using read lengths of 301/301. 20% PhiX was spiked in to increase sequence diversity of the libraries due to the predominance of the antibody knockin derived reads in the repertoires.

Annotation of antibody sequences with immunogenetic information, clonal clustering and clonal tree reconstruction were performed using the software Cloanalyst (https://www.bu.edu/computationalimmunology/research/software/). Due to variable recovery rates of heavy and light chain reads using the bulk single chain NGS sequencing of mouse BCR repertoires and the depth required for sufficient statistics for analysis of rare mutational events, mouse repertoires that had <1000 unique and functional bnAb UCA knock-in derived reads recovered were excluded from the data analysis. Functional heavy and light chains were defined by the presence of required immunogenetic characteristics as determined by Clonanalyst including: V gene regions of at least 200 base pairs in length, presence of invariant cysteines and tryptophan (heavy) or phenylalanine (light), CDR3s in reading frame 1, non-zero CDR3 length, and absence of stop codons. Reported mutation frequencies were calculated at the nucleotide level from the beginning of the antibody sequence through the invariant cysteine codon that precedes CDR3. Analysis of the probability of mutations was performed using the computational program ARMADiLLO as previously described (*3, 8*)

## Acknowledgments

Cryo-EM data were collected at the Duke Krios at the Duke University Shared Materials Instrumentation Facility (SMIF), a member of the North Carolina Research Triangle Nanotechnology Network (RTNN), which is supported by the National Science Foundation (award number ECCS-2025064) as part of the National Nanotechnology Coordinated Infrastructure (NNCI). This study used the computational resources offered by Duke Research Computing (http://rc.duke.edu; NIH 1S10OD018164-01) at Duke University. This project was supported by NIH, NIAID, Division of AIDS Consortia for HIV/AIDS Vaccine Development (CHAVD) Grant UM1AI144371 (B.F.H), R01AI145687 (P.A.), U54AI170752 (R.C.H. and P.A.), DP2-AI164323-03 (R.H.), and Translating Duke Health Initiative (R.C.H. and P.A.).

## Author Contributions

RCH and BFH designed the study. RCH prepared and edited the manuscript. RCH designed, conducted, and interpreted the molecular dynamics simulation results. RCH and SK analyzed the molecular dynamics simulations. CS, AW, YB, JB, MM and KOS prepared all proteins for this study. RCH, VS, KM, and PA collected and analyzed the cryo-EM data. RCH designed each immunogen. KA and MA collected and analyzed the SPR. data. AN conducted the mouse immunizations. CJ and KOS collected and analyzed pseudovirus neutralization data. DC and XL isolated B cells. KW, SV, and MB sequenced antibodies. KW and BK performed neutralization signature analyses. JL and KOS collected and analyzed mammalian display data. MT and FA developed the IA4 mouse model. KW and BK performed signature analysis. All authors read and edited the manuscript.

## Data and Code Availability

All data and code from this manuscript are available on reasonable request.

---

**Table 1.** Cryo-EM Data Collection and Refinement Statistics.

|  | DH270.6 – DE3 | VRC01 – DE3 |
| --- | --- | --- |
| **Data Collection** |  |  |
| Microscope | Titan Krios | Titan Krios |
| Voltage (kV) | 300 | 300 |
| Electron dose (e-/Å<sup>2</sup>) | ~60 | ~60 |
| Detector | Gatan K3 | Gatan K3 |
| Pixel Size (Å) | 1.08 | 1.08 |
| Defocus Range (µm) | 0.42-4.71 | 0.47-3.674 |
| Magnification | 105000 | 105000 |
| **Reconstruction** |  |  |
| Software | cryoSPARC | cryoSPARC |
| Particles | 236154 | 220881 |
| Symmetry | C3 | C3 |
| Box size (pix) | 300 | 300 |
| Resolution (Å) (FSC0.143)<sup>§</sup> | 3.6 | 3.7 |
| **Validation** |  |  |
| Protein residues | 2484 | 2457 |
| EMRinger Score | 2.79 | 2.06 |
| R.m.s. deviations |  |  |
| Bond lengths (Å) | 0.005 | 0.004 |
| Bond angles (°) | 0.905 | 0.814 |
| Molprobity score | 1.47 | 1.53 |
| Clash score | 2.03 | 2.98 |
| Rotamer outliers (%) | 0.00 | 0.00 |
| Ramachandran |  |  |
| Favored regions (%) | 92.79 | 93.21 |
| Disallowed regions (%) | 0.49 | 0.12 |

§ Resolutions are reported according to the FSC 0.143 gold-standard criterion

**Table 2.**
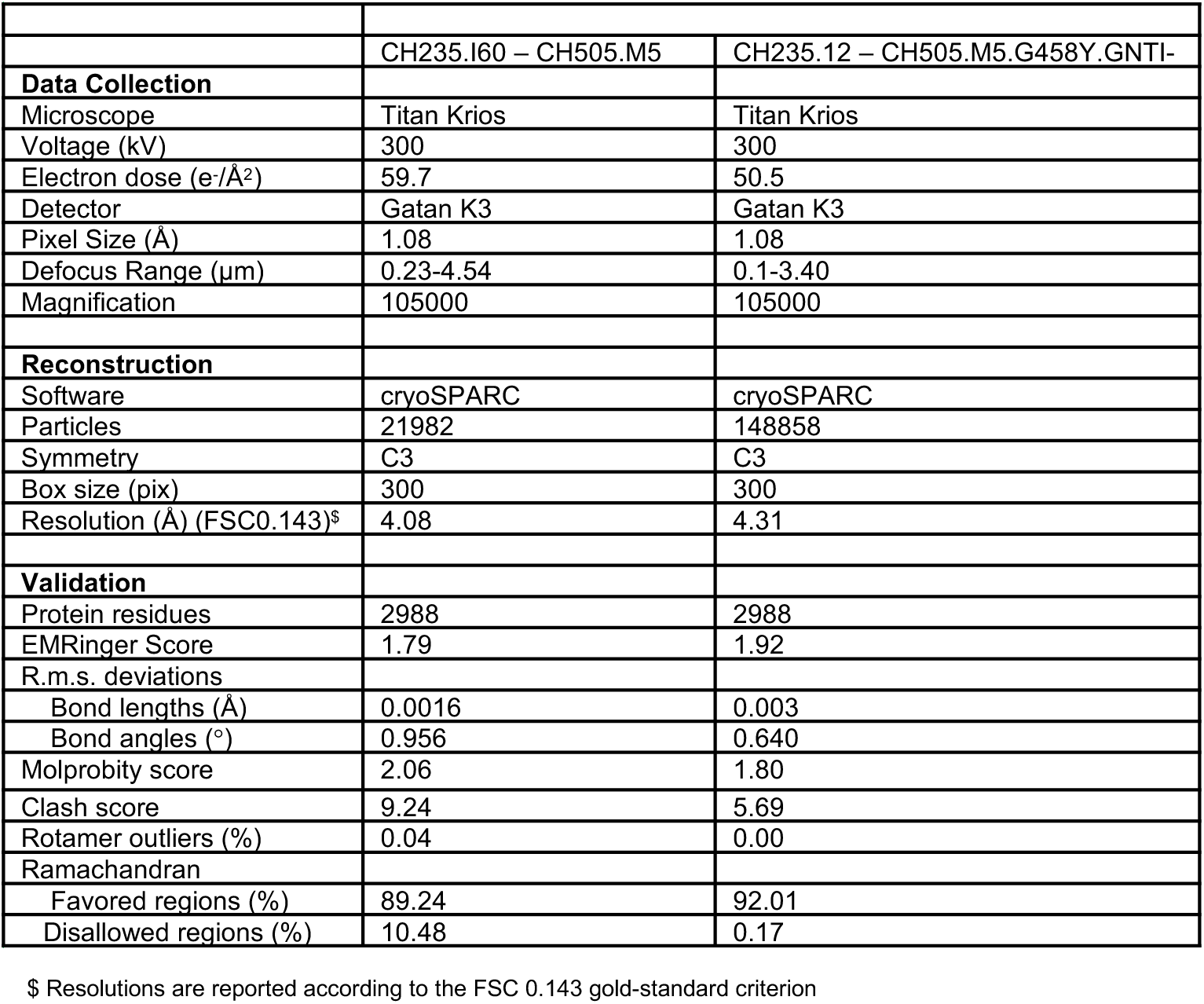
Cryo-EM Data Collection and Refinement Statistics.

**Table 3.** Cryo-EM Data Collection and Refinement Statistics.

|  |  |
| --- | --- |
|  | CE3 |
| **Data Collection** |  |
| Microscope | Titan Krios |
| Voltage (kV) | 300 |
| Electron dose (e-/Å<sup>2</sup>) | 60.5 |
| Detector | Gatan K3 |
| Pixel Size (Å) | 1.08 |
| Defocus Range (µm) | 0.46-4.62 |
| Magnification | 105000 |
| **Reconstruction** |  |
| Software | cryoSPARC |
| Particles | 106813 |
| Symmetry | C3 |
| Box size (pix) | 320 |
| Resolution (Å) (FSC0.143)<sup>\$</sup> | 3.9 |
| **Validation** |  |
| Protein residues | 1734 |
| EMRinger Score | 2.44 |
| R.m.s. deviations |  |
| Bond lengths (Å) | 0.004 |
| Bond angles (°) | 0.843 |
| Molprobity score | 1.38 |
| Clash score | 1.94 |
| Rotamer outliers (%) | 0.00 |
| Ramachandran |  |
| Favored regions (%) | 93.65 |
| Disallowed regions (%) | 0.52 |

<sup>\$</sup> Resolutions are reported according to the FSC 0.143 gold-standard criterion

