## Supplemental Material for "Engineering immunogens that select for specific mutations in HIV broadly neutralizing antibodies"

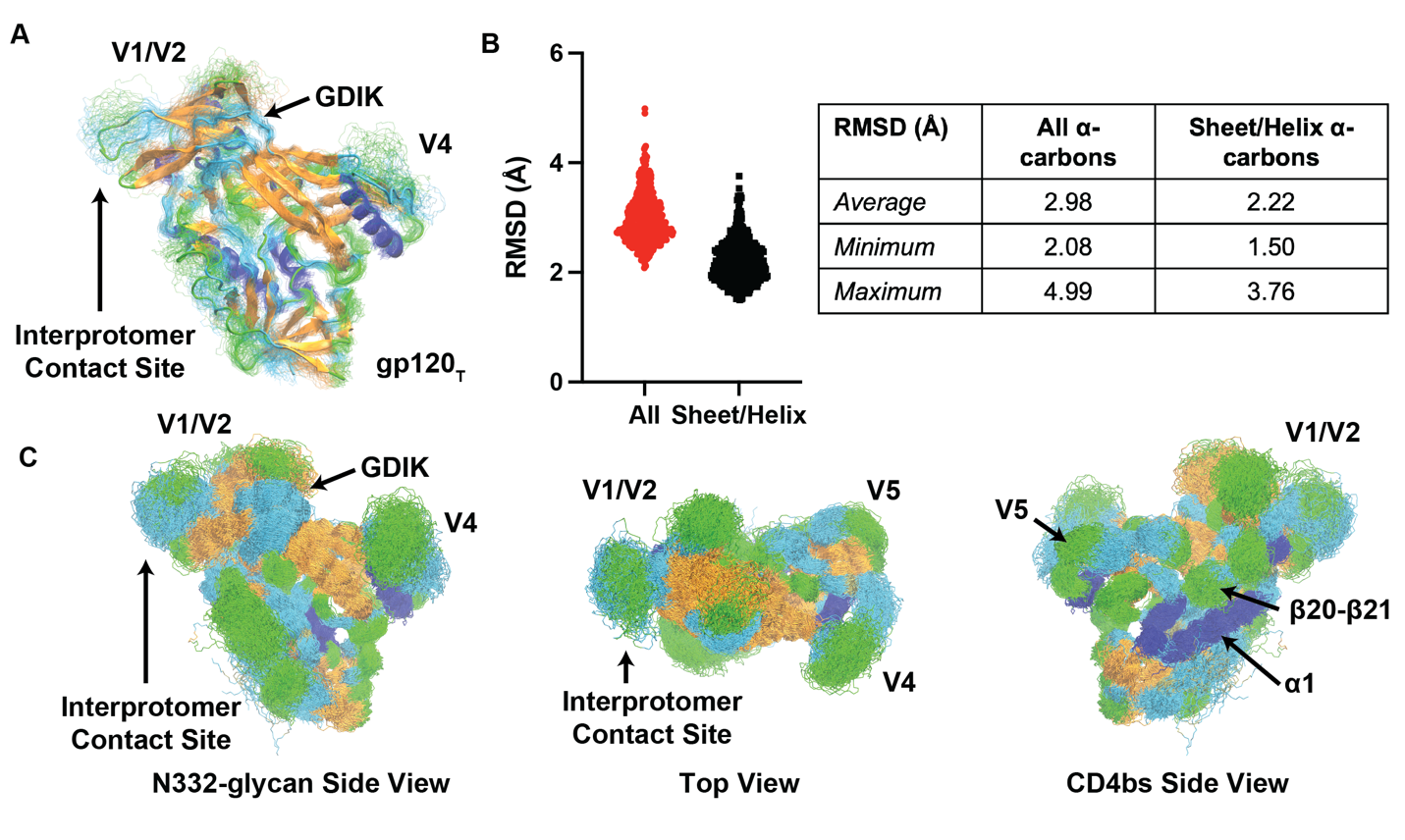

Supplemental Figure 1

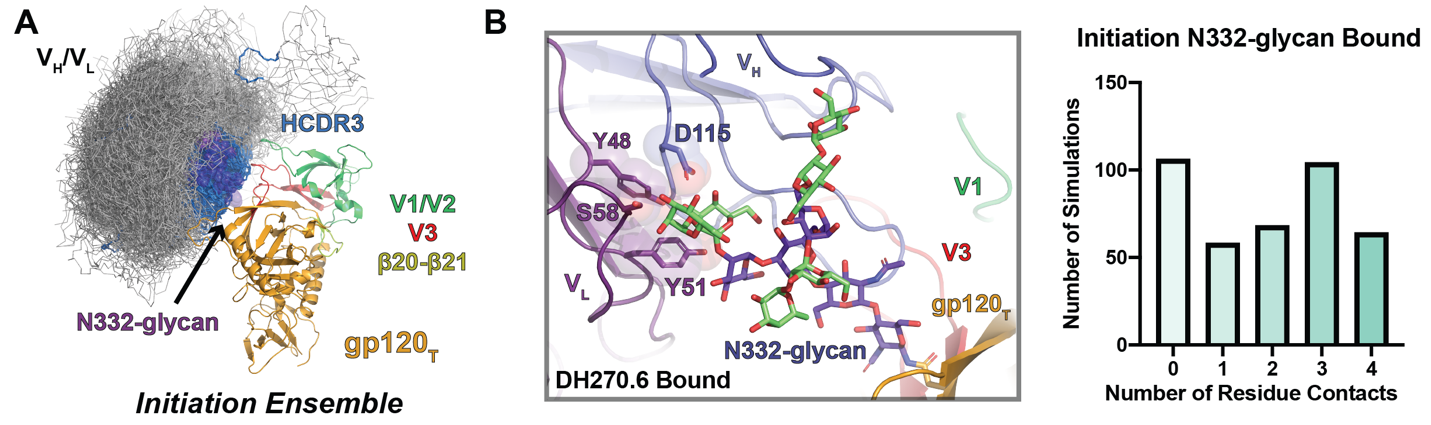

Supplemental Figure 2

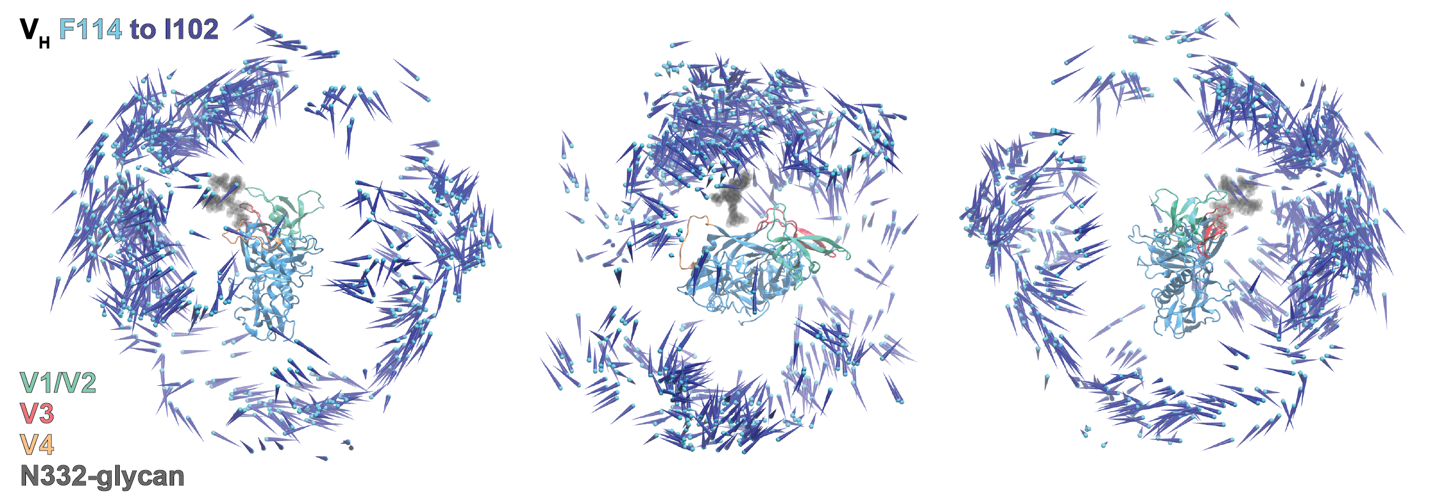

Supplemental Figure 3

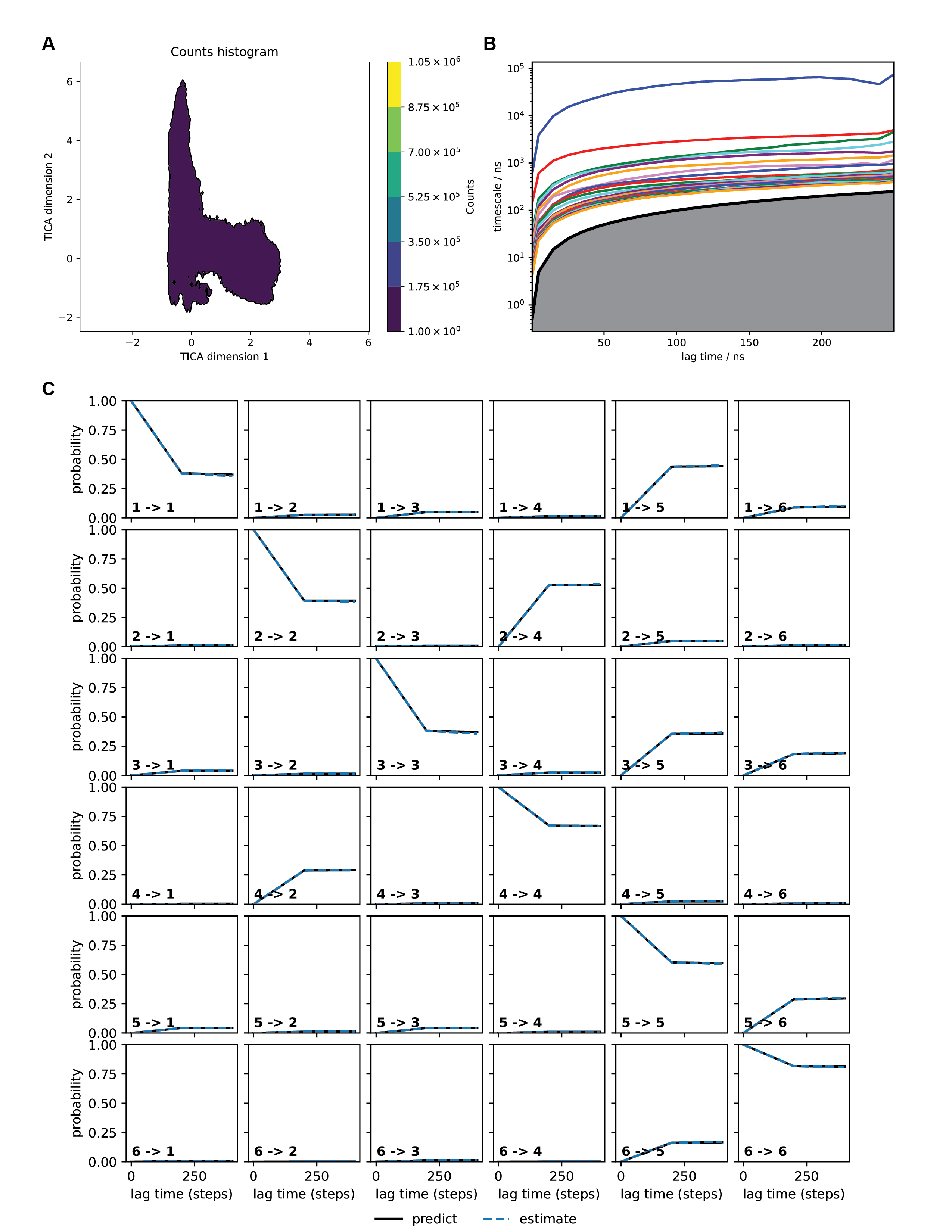

Supplemental Figure 4

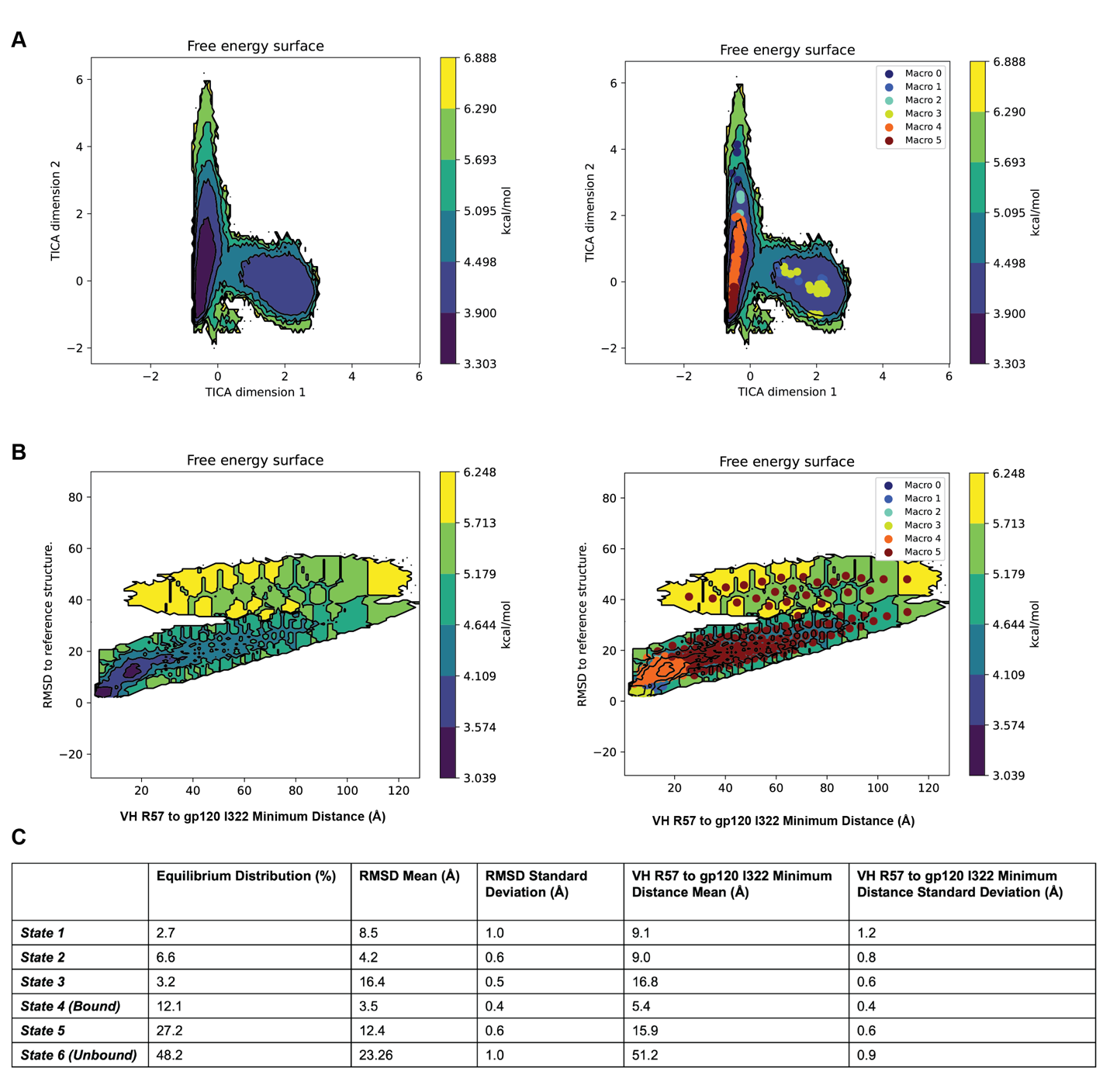

Supplemental Figure 5

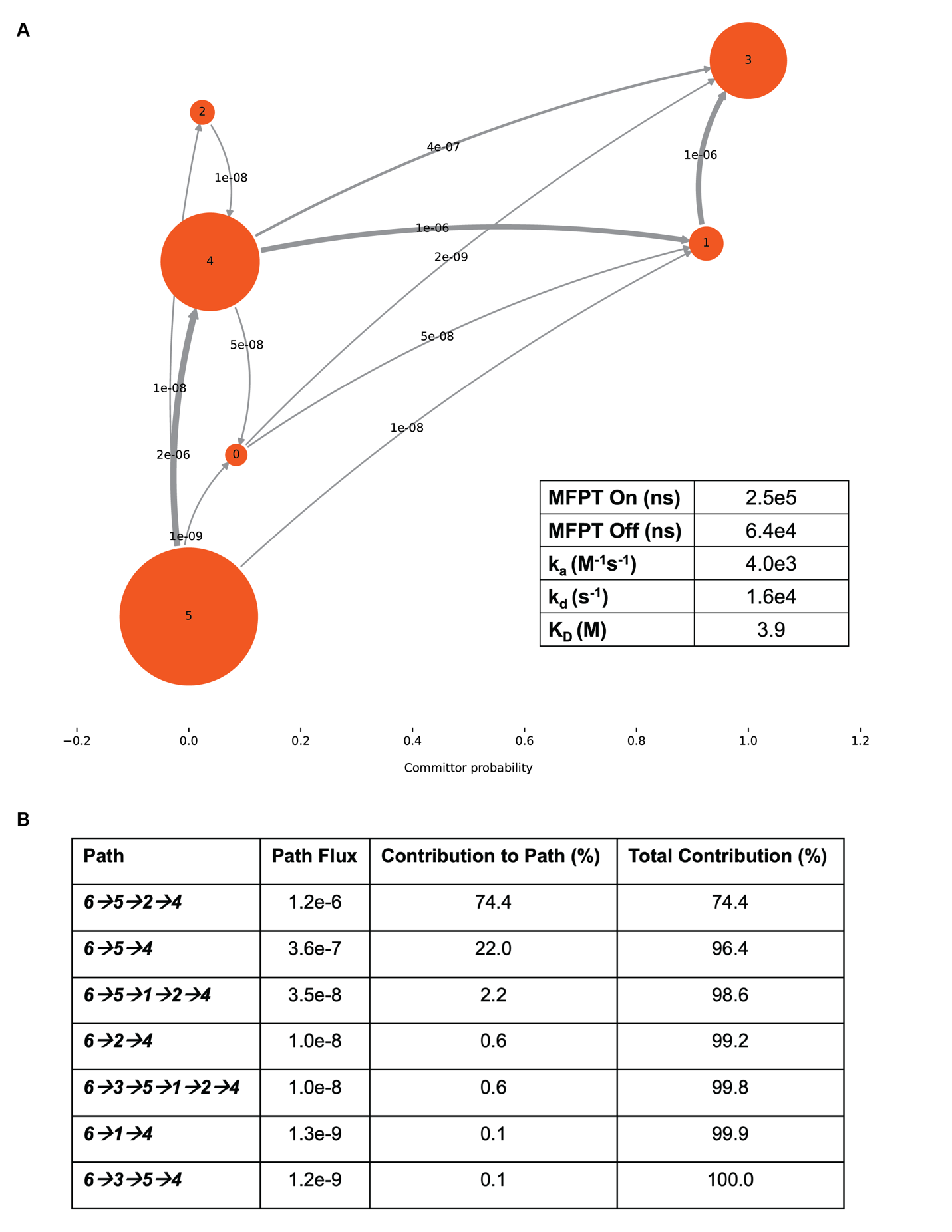

Supplemental Figure 6

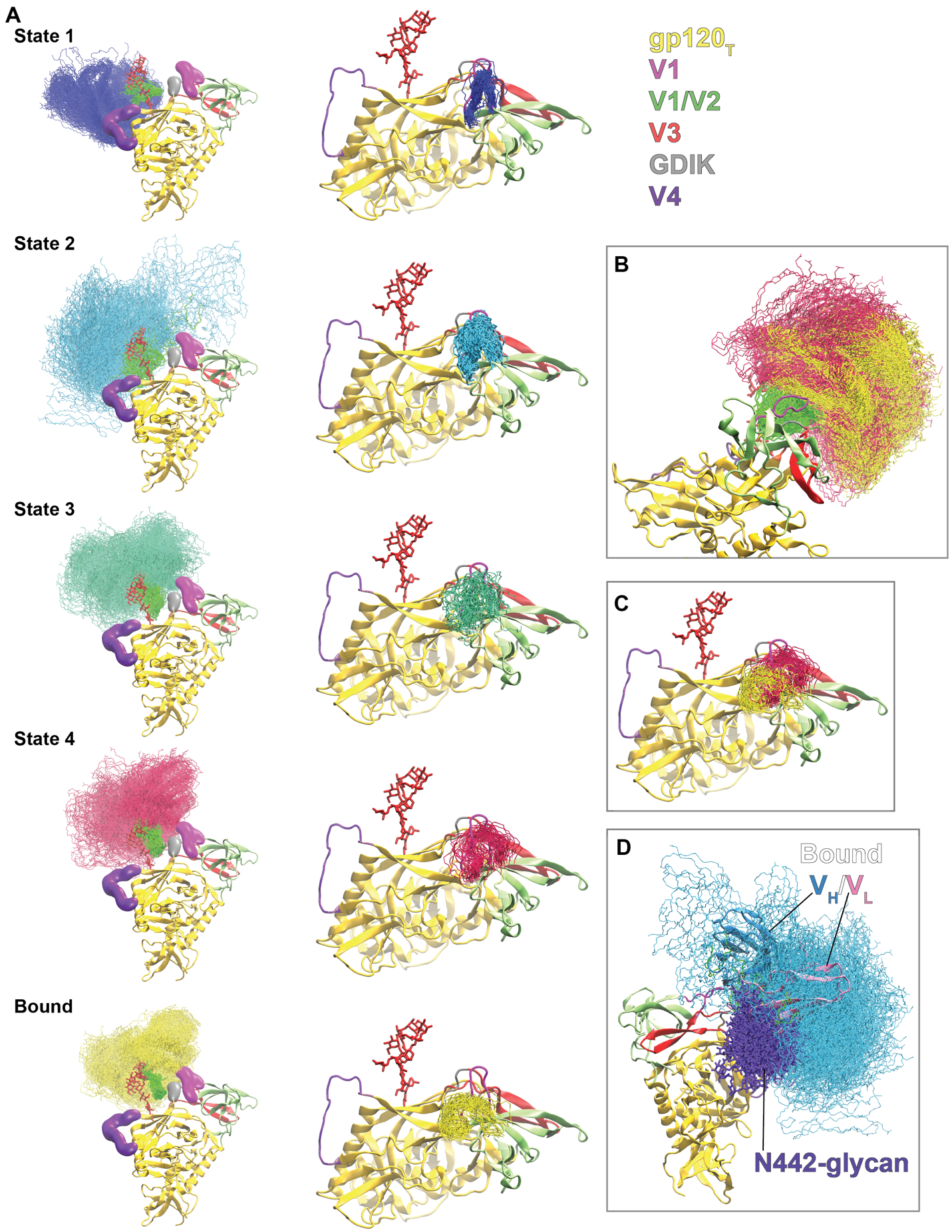
Supplemental Figure 7

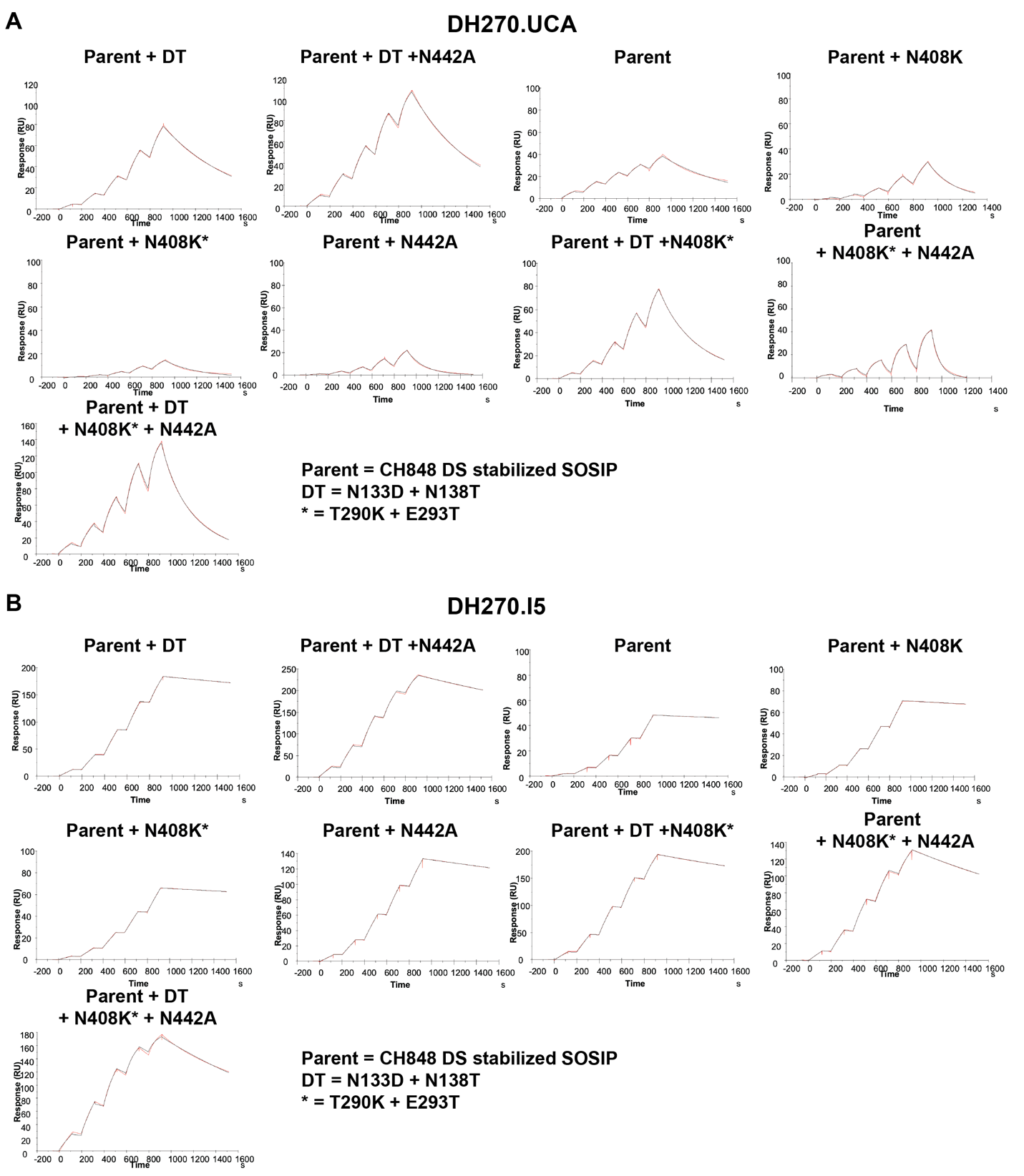
Supplemental Figure 8

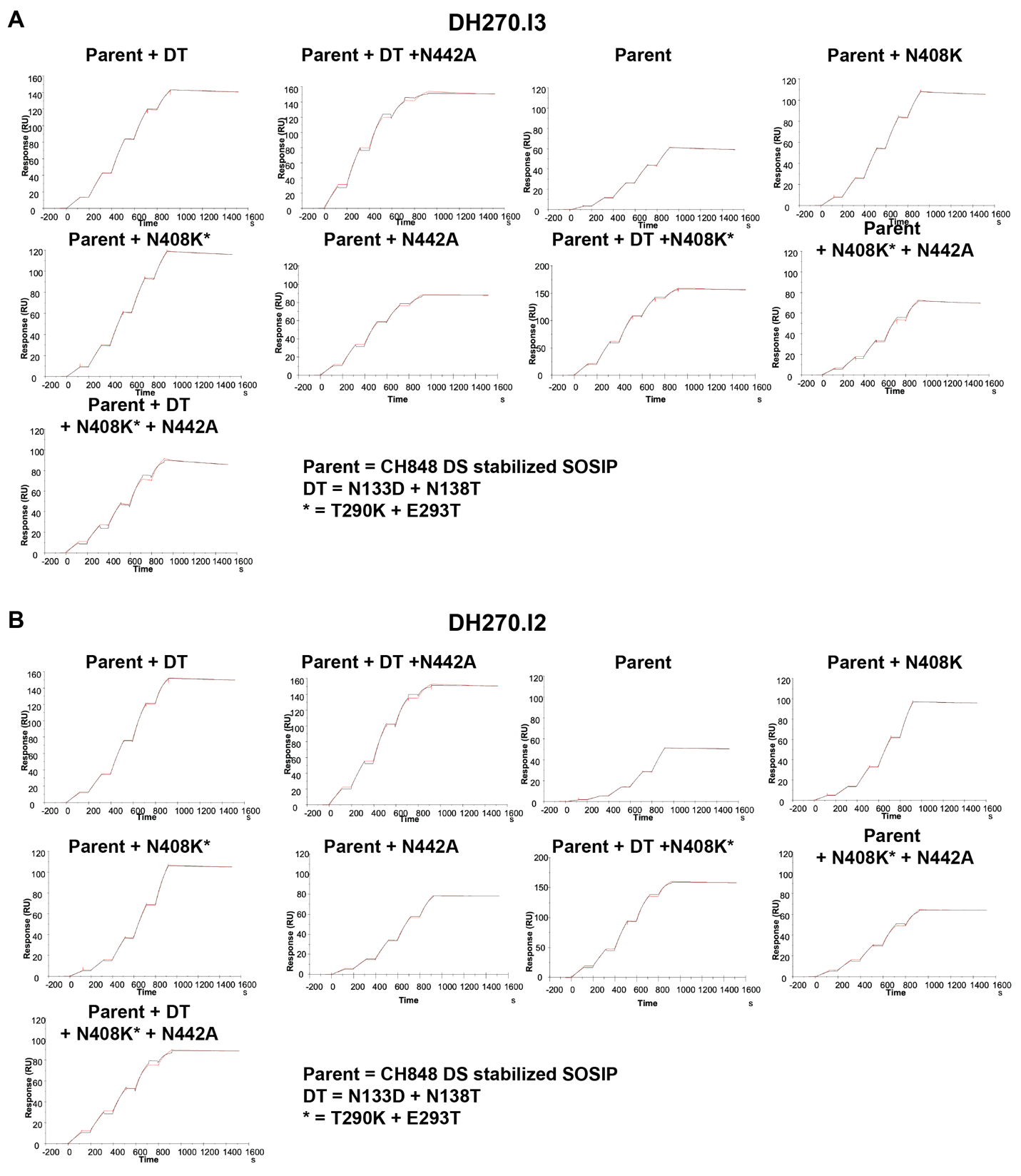

Supplemental Figure 9

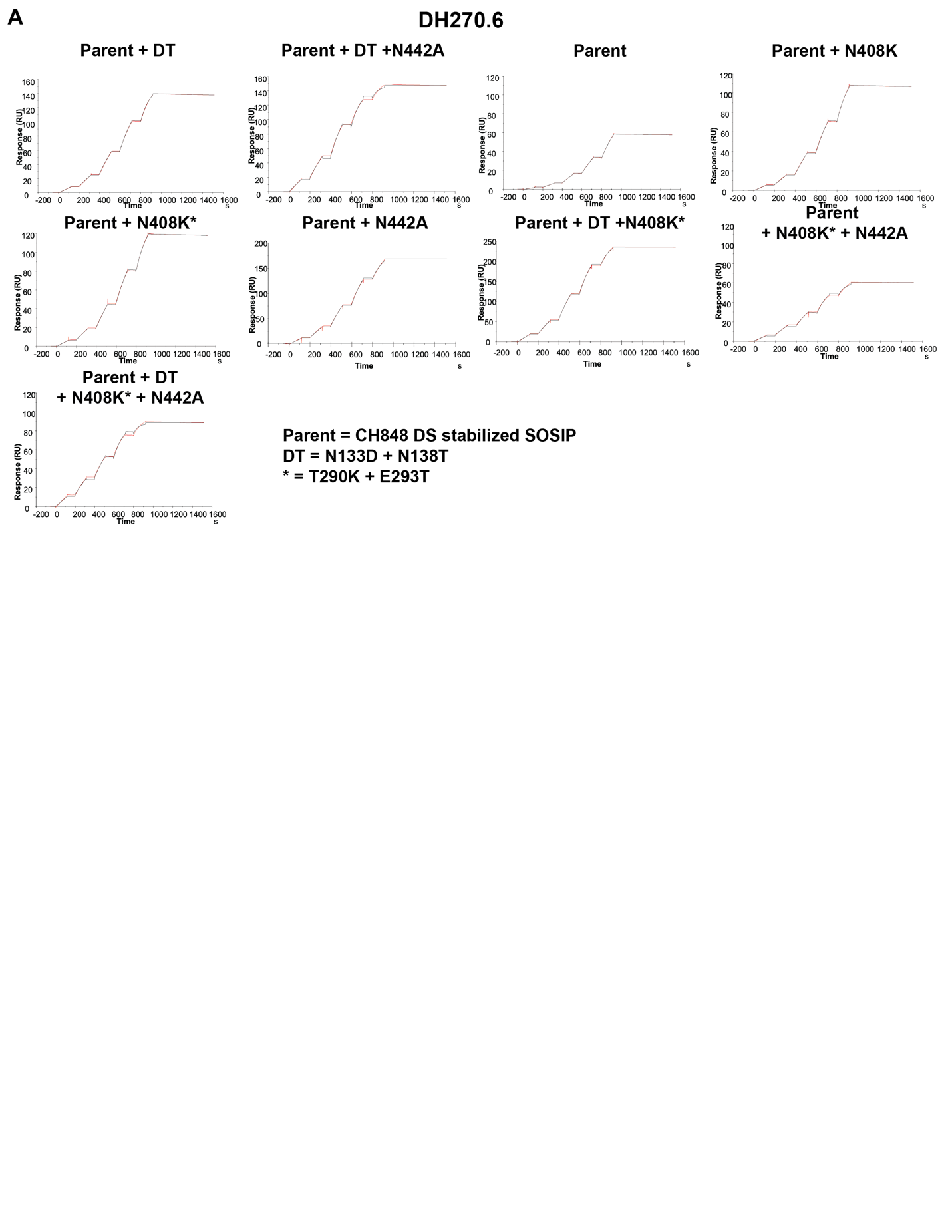

Supplemental Figure 10

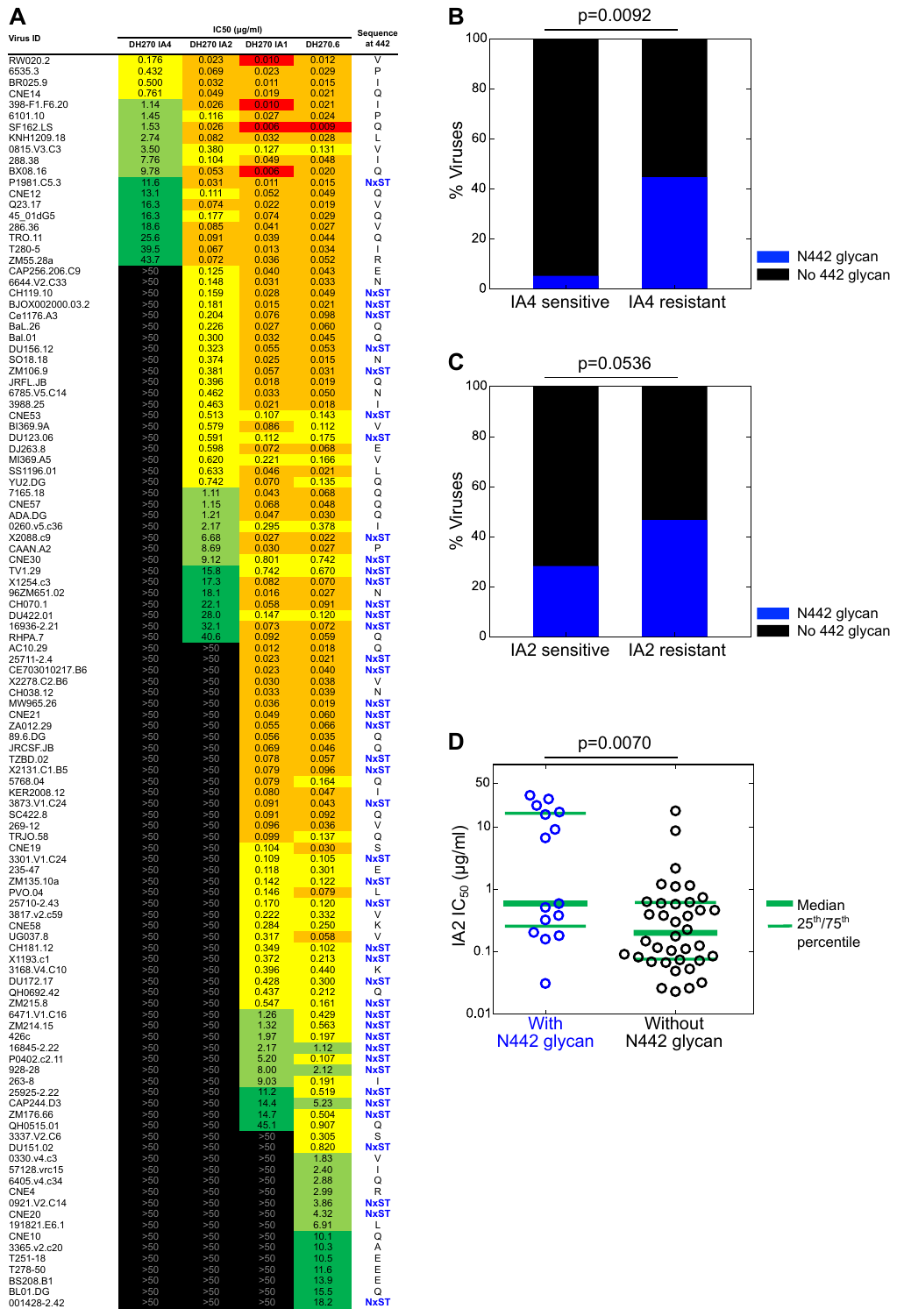

Supplemental Figure 11

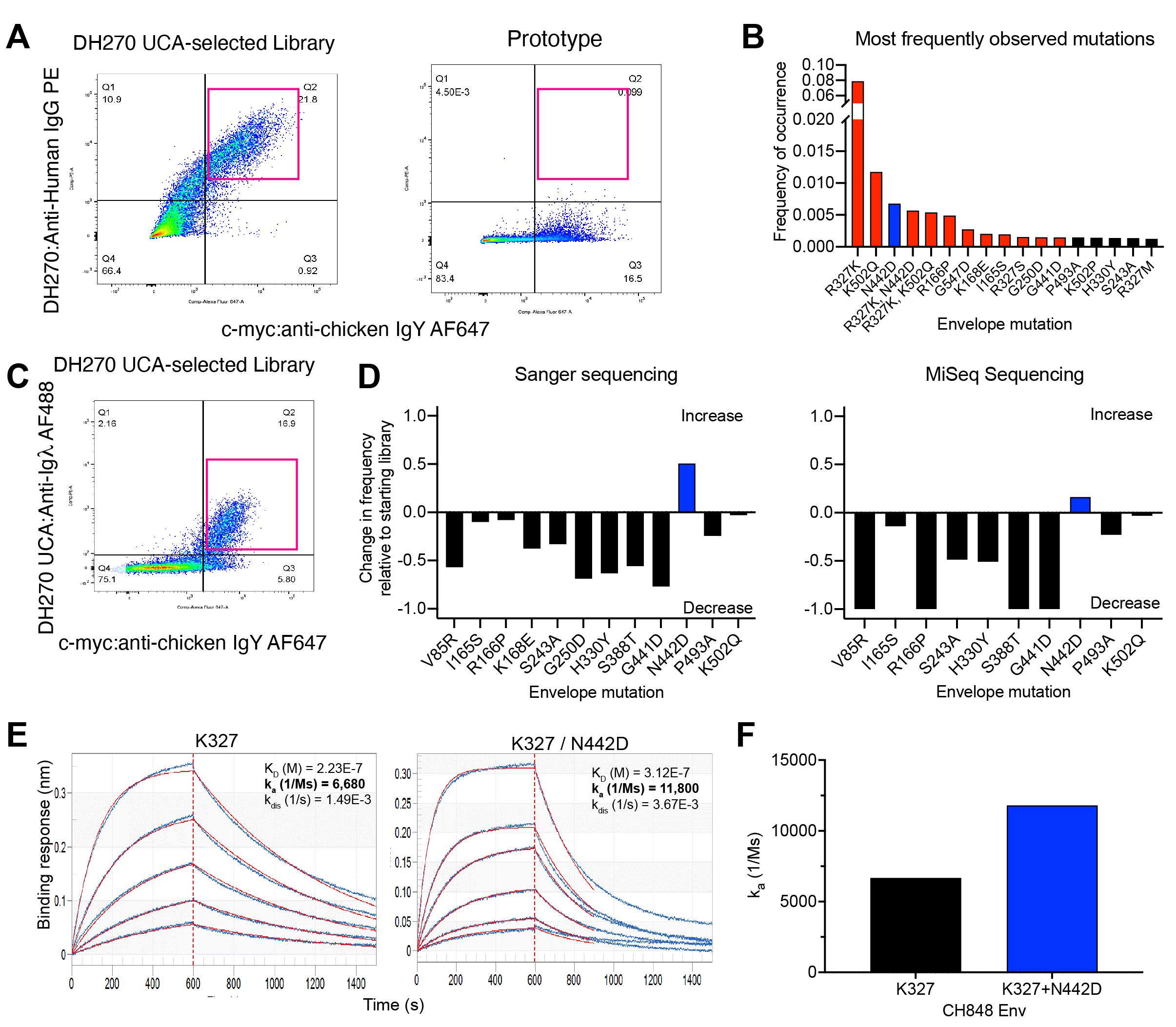

Supplemental Figure 12

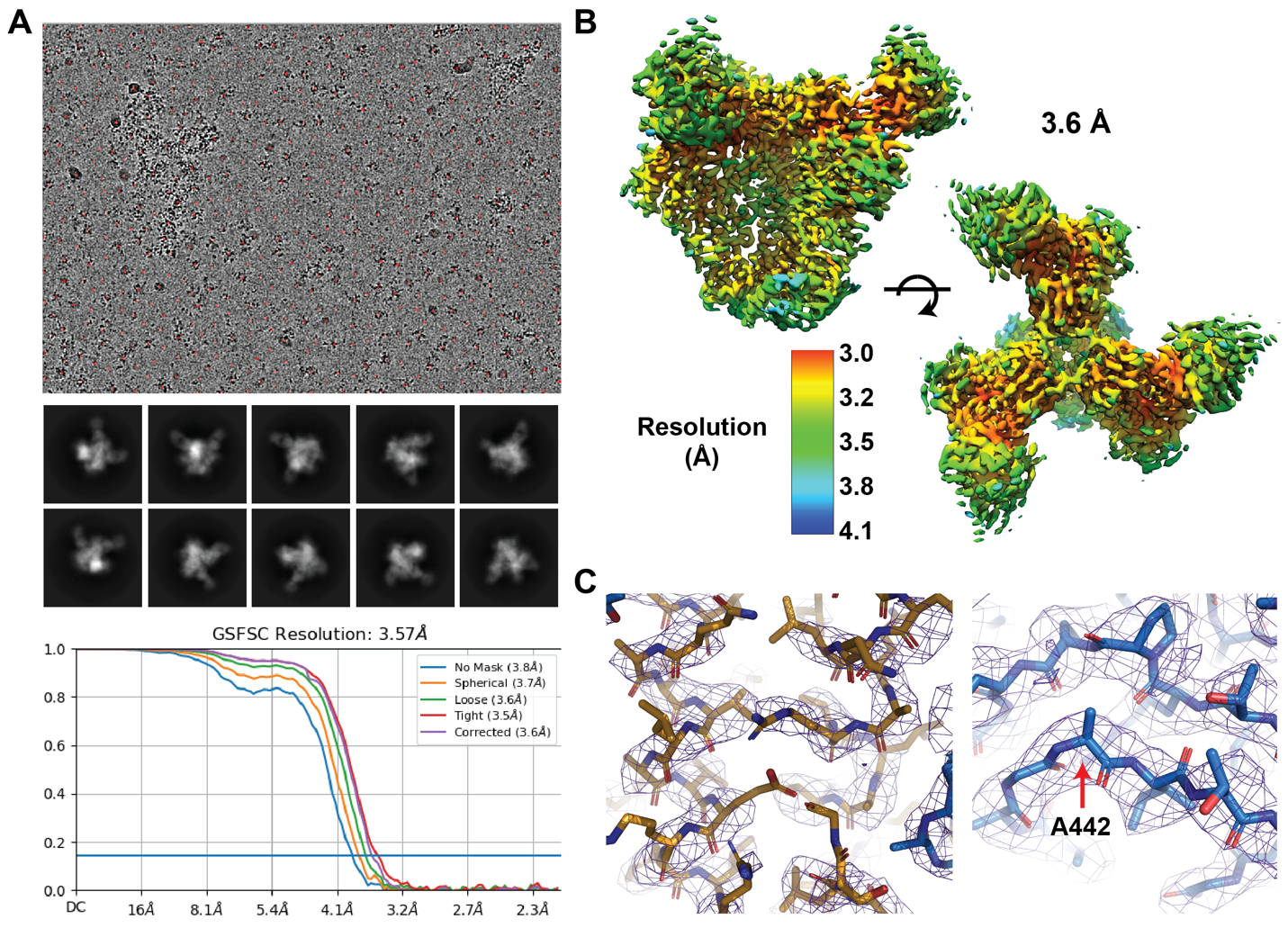

Supplemental Figure 13

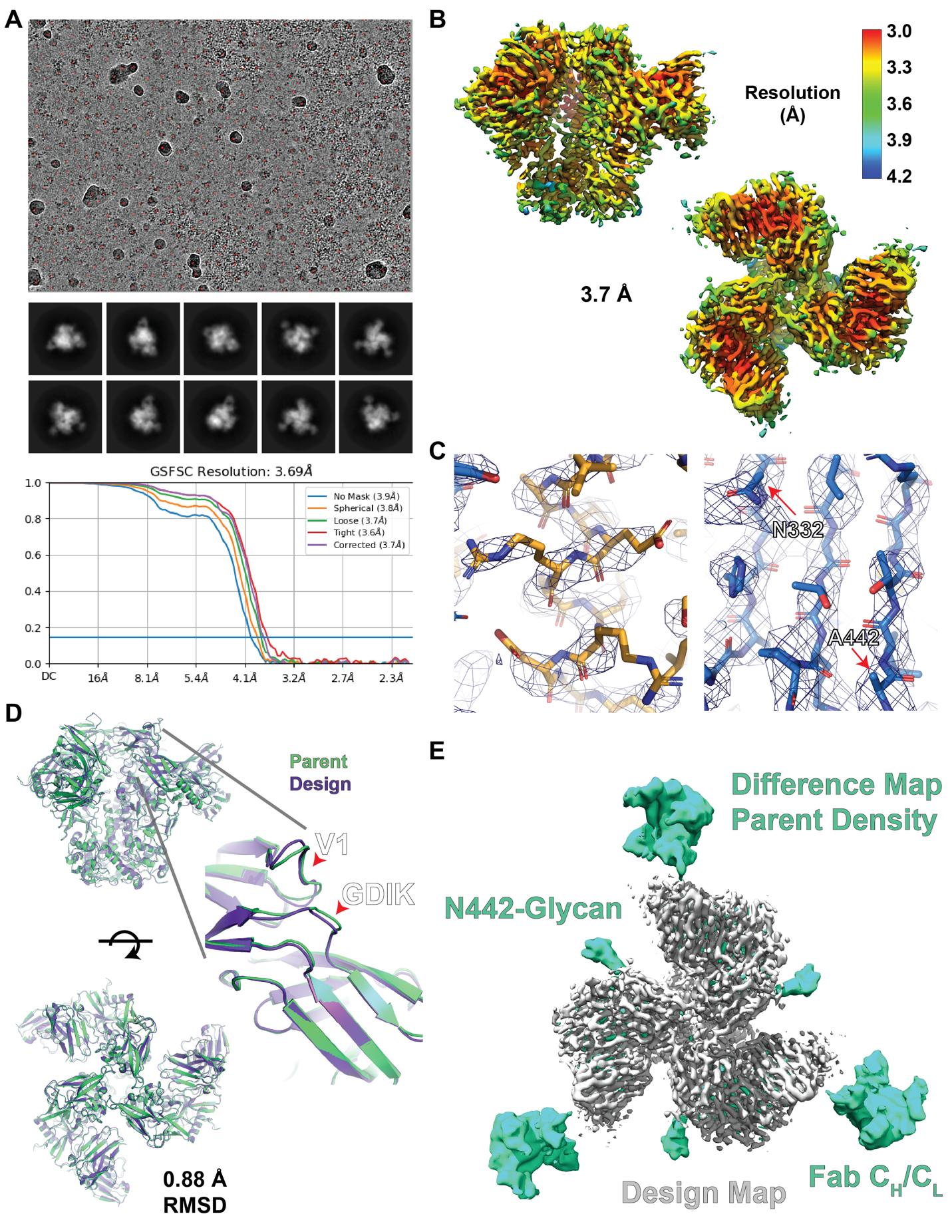

Supplemental Figure 14

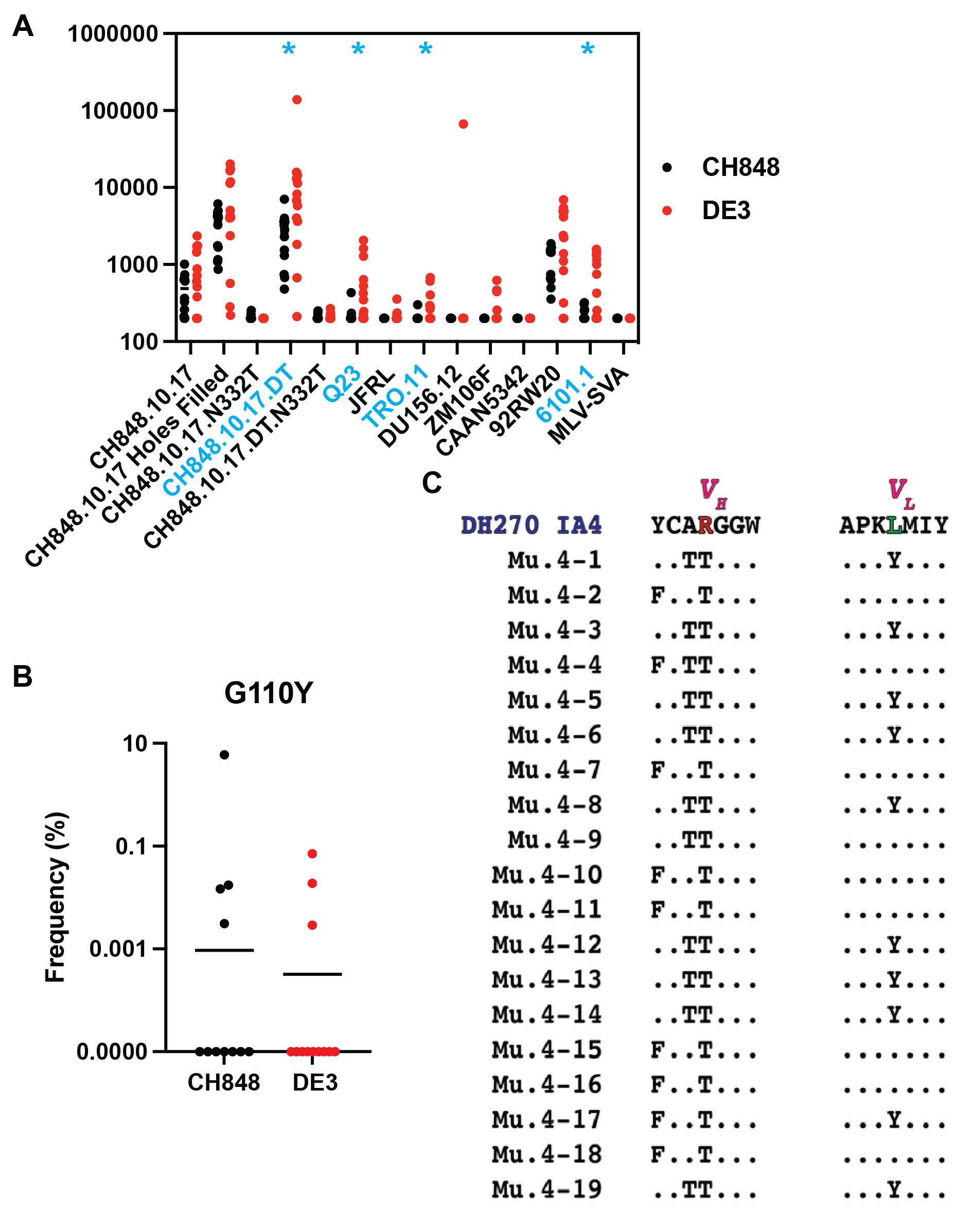
Supplemental Figure 15

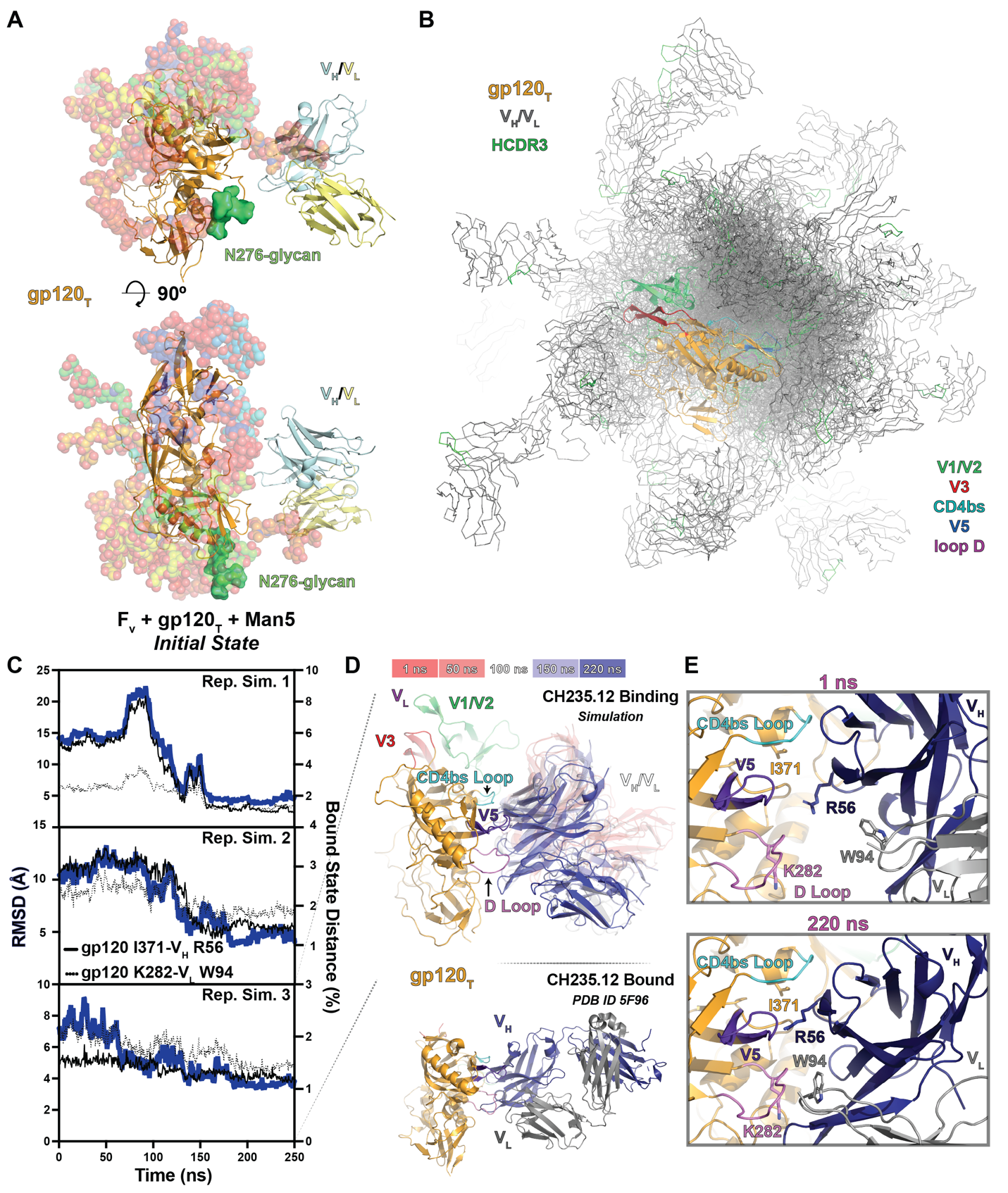

Supplemental Figure 16

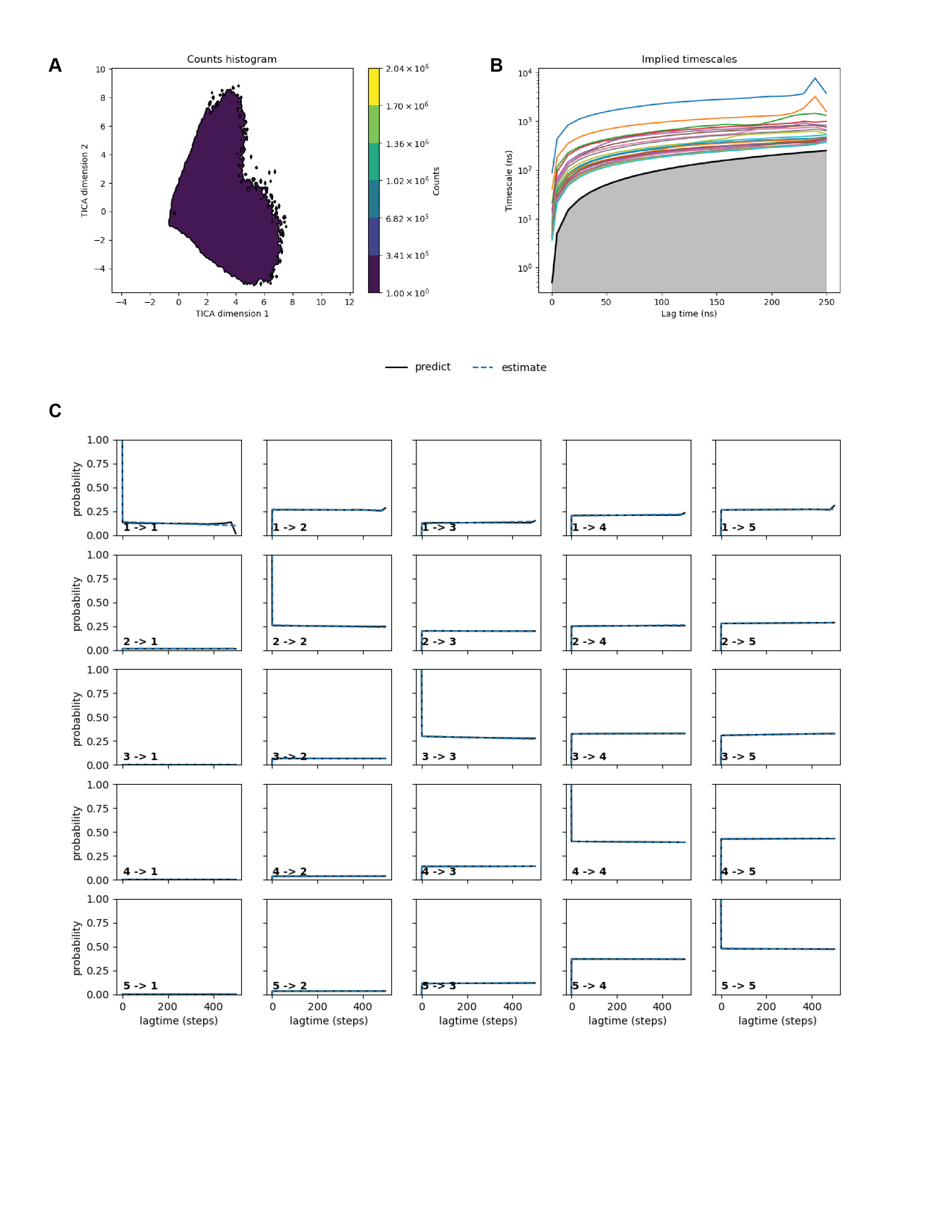

Supplemental Figure 17

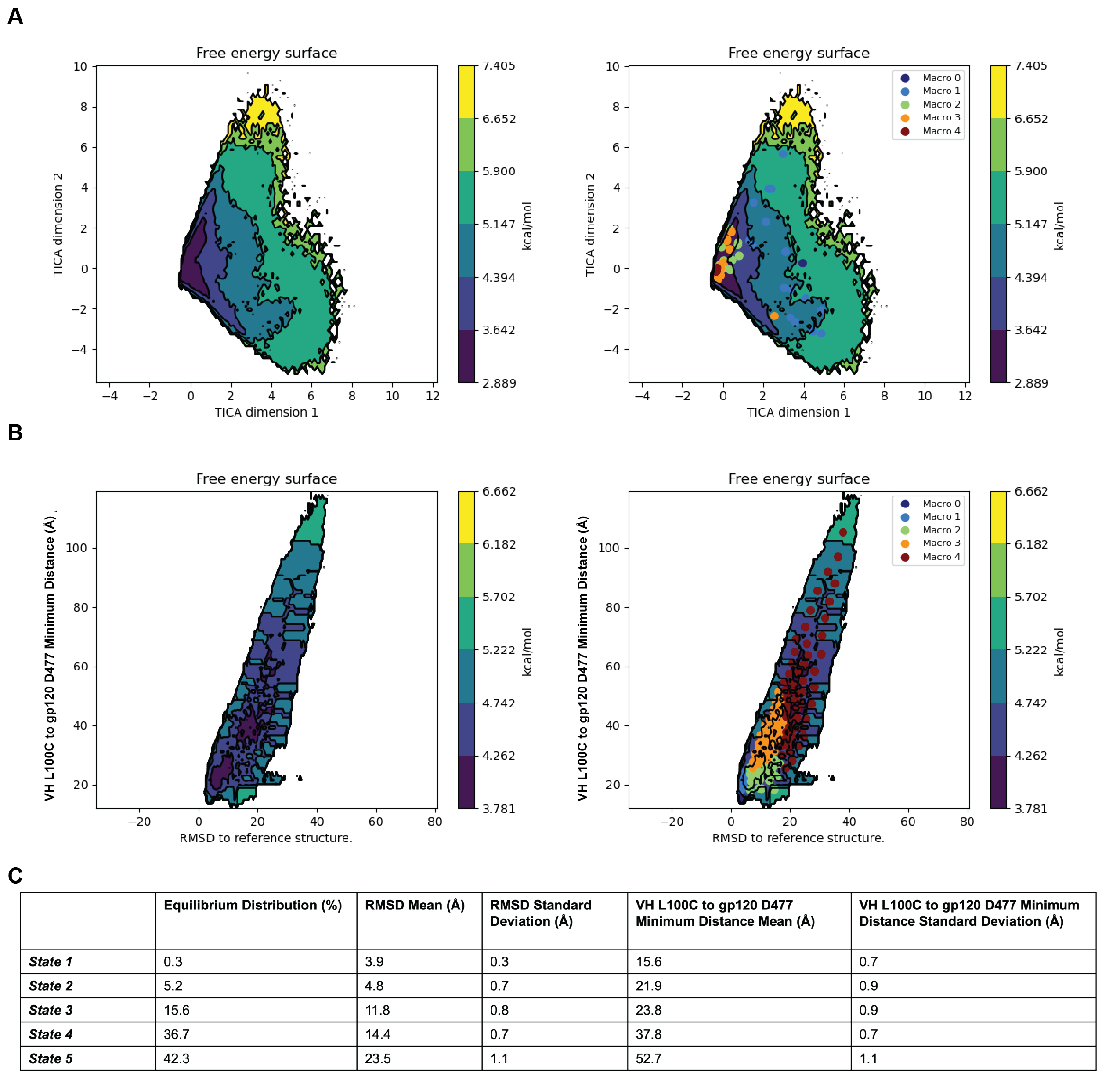
Supplemental Figure 18

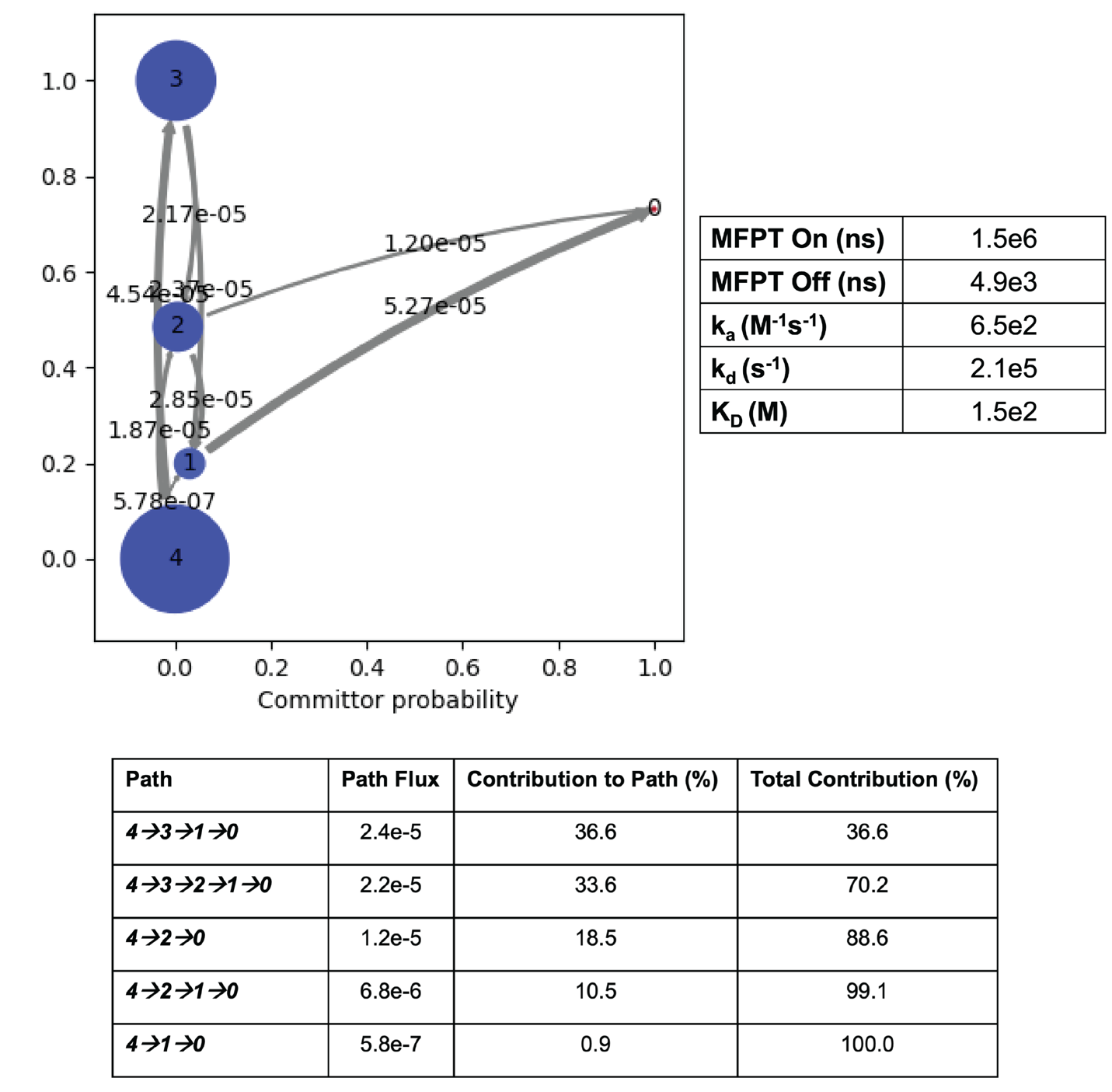

Supplemental Figure 19

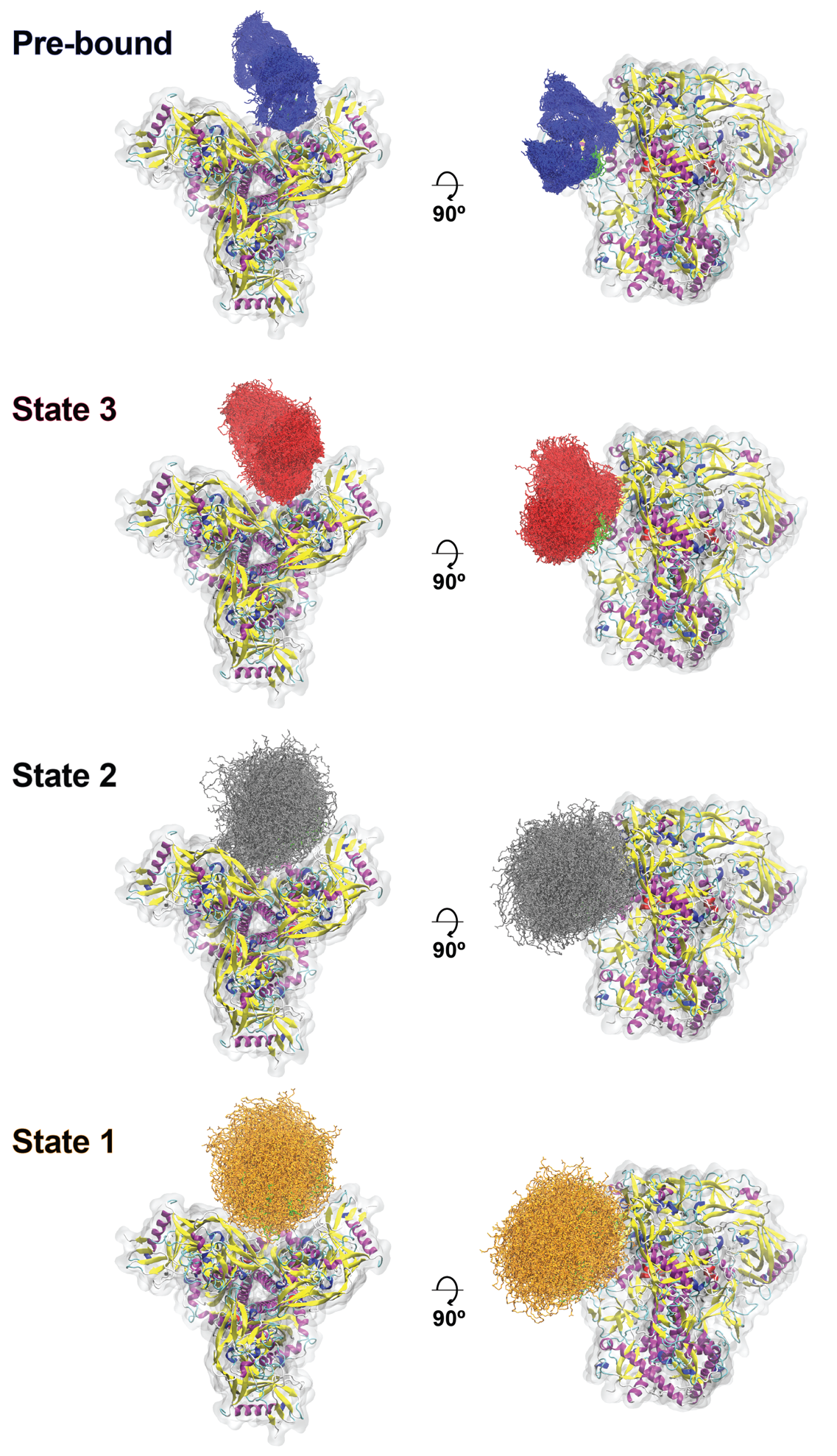

Supplemental Figure 20

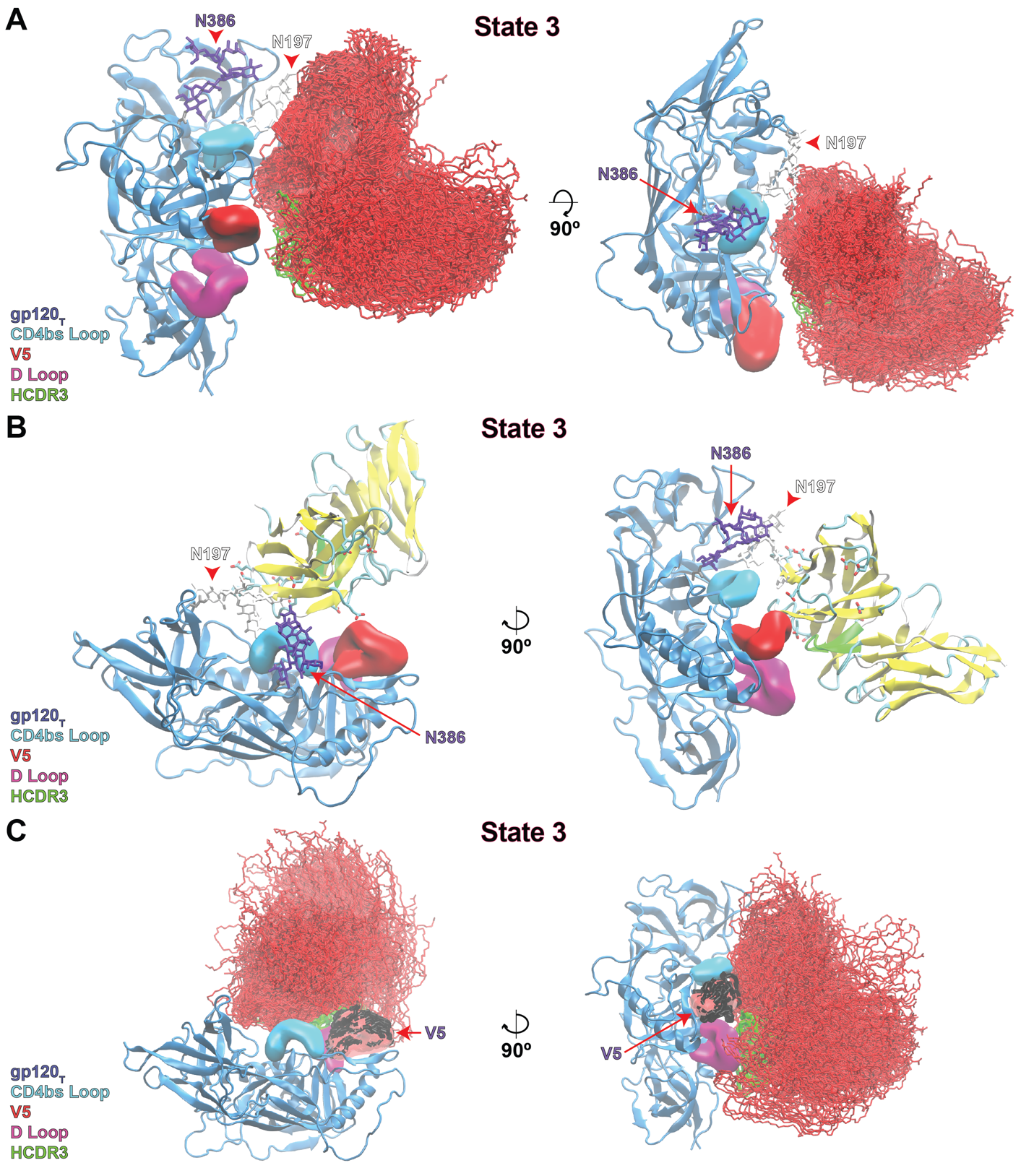

Supplemental Figure 21

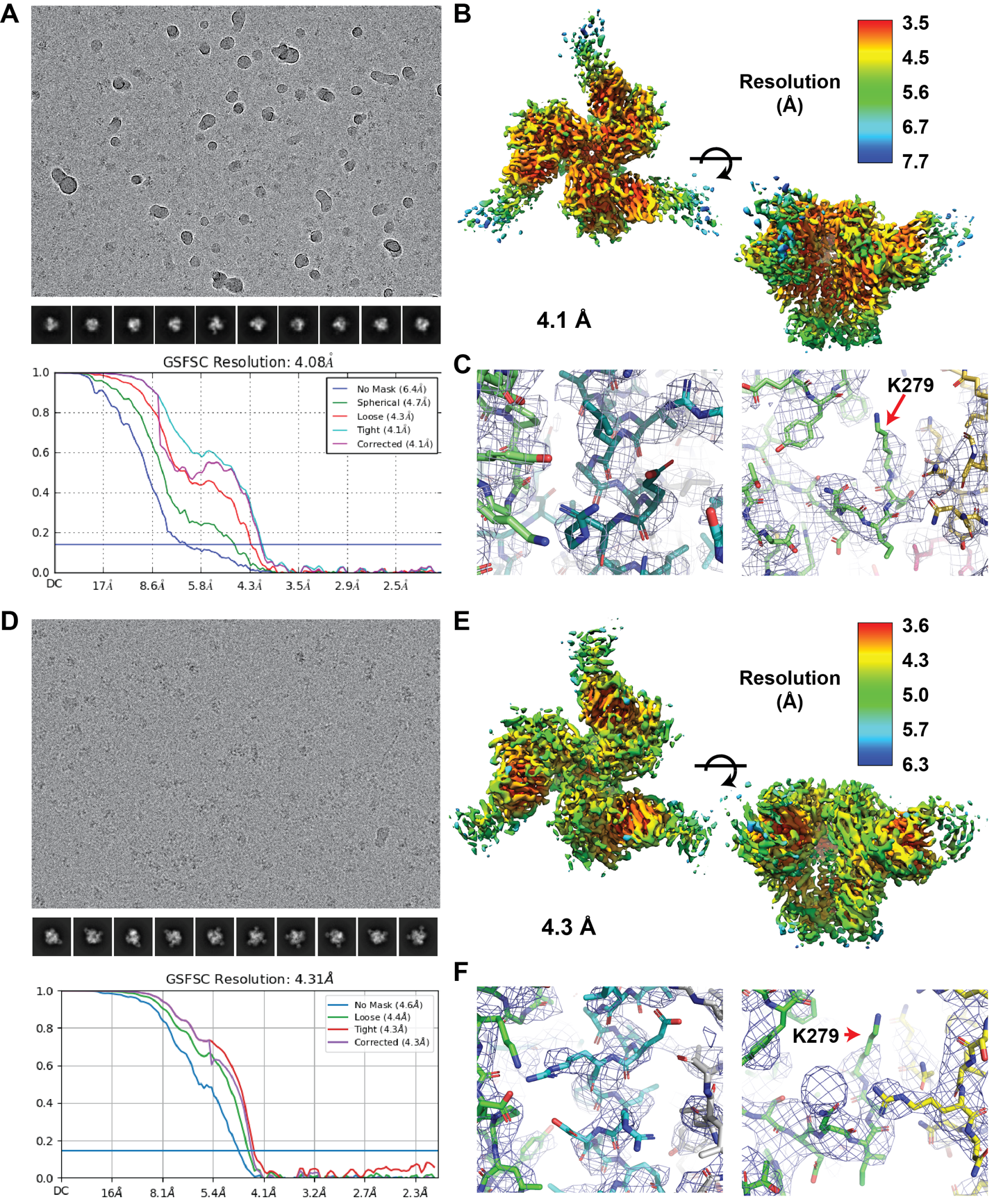

Supplemental Figure 22

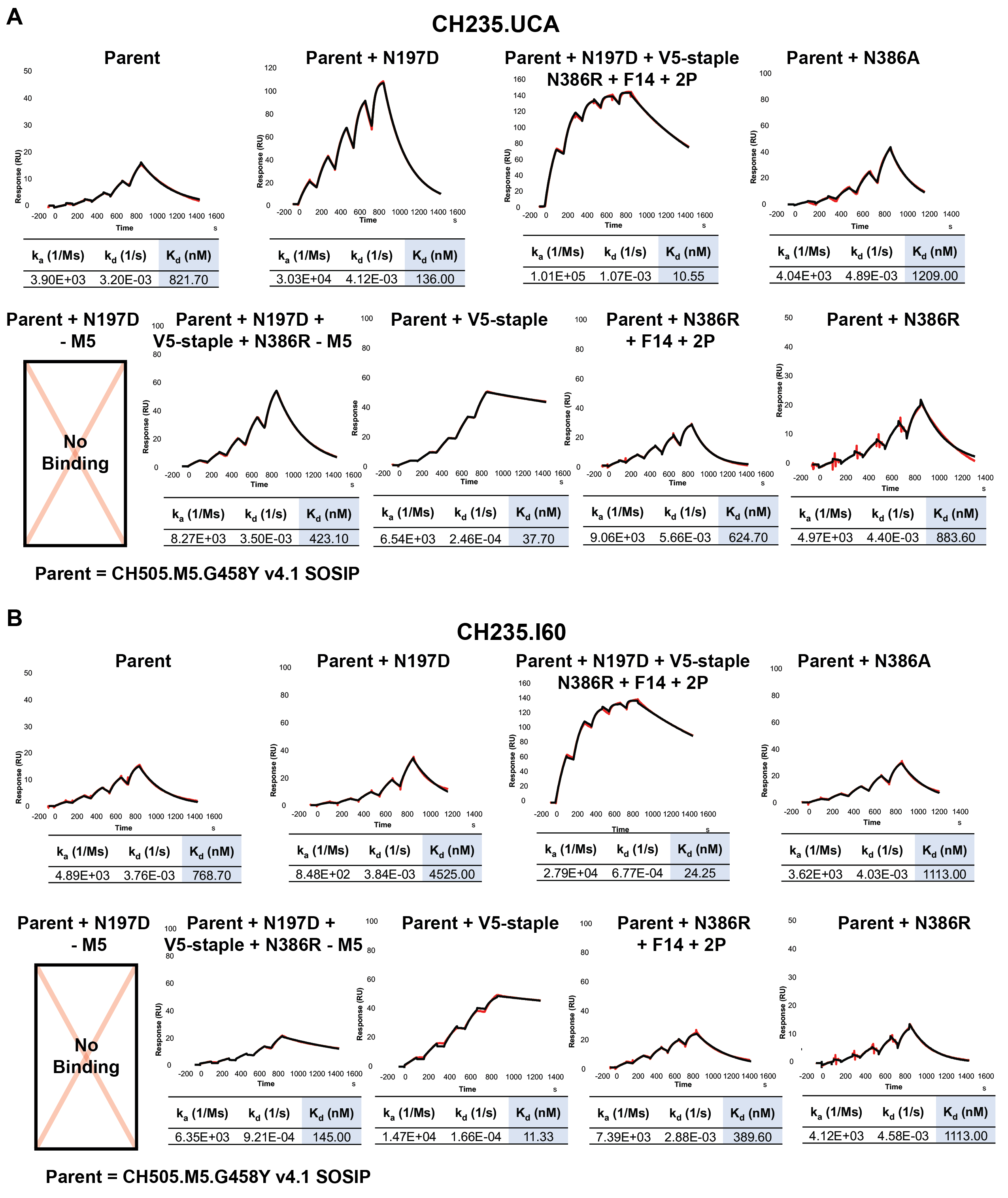

Supplemental Figure 23

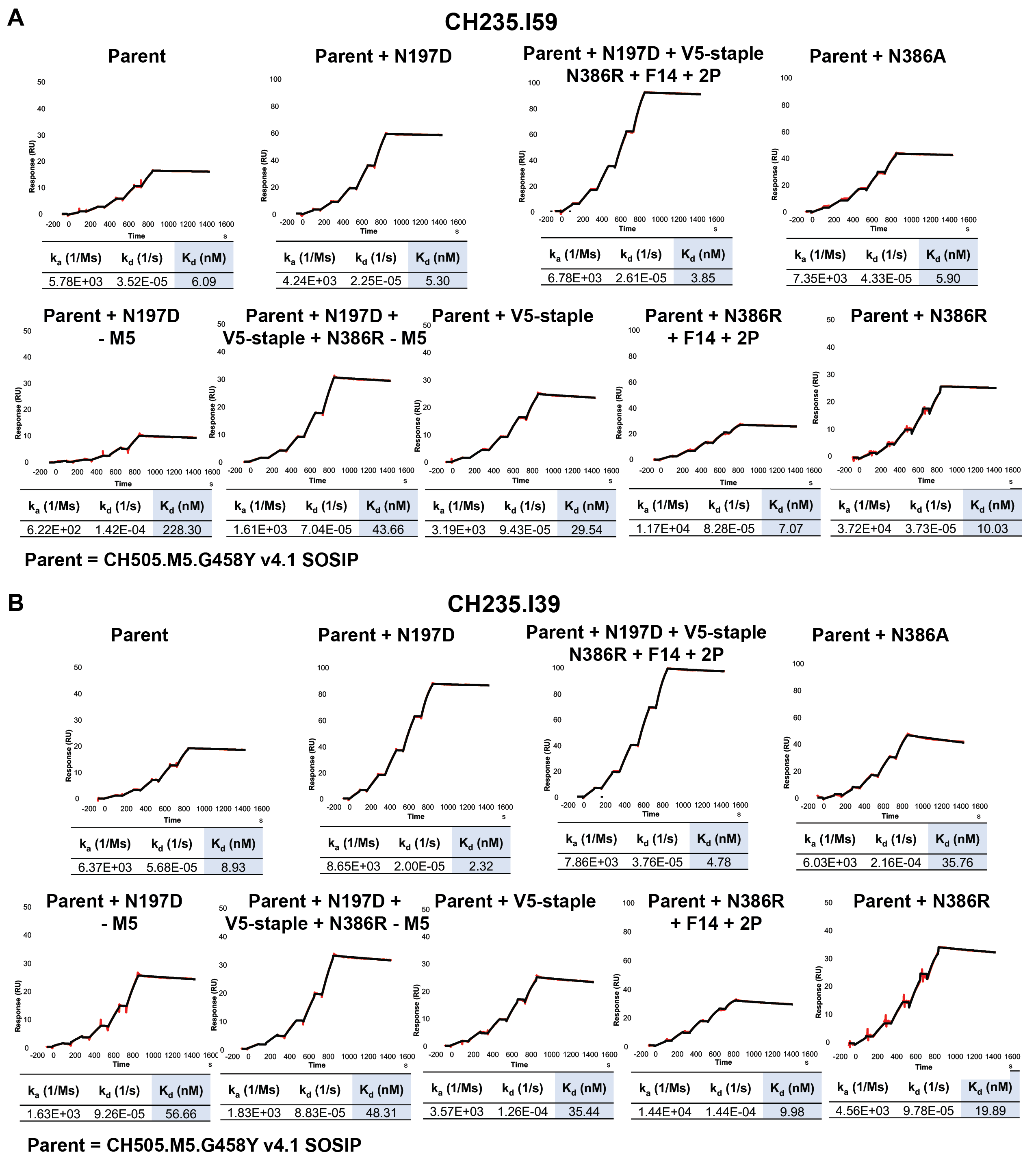

Supplemental Figure 24

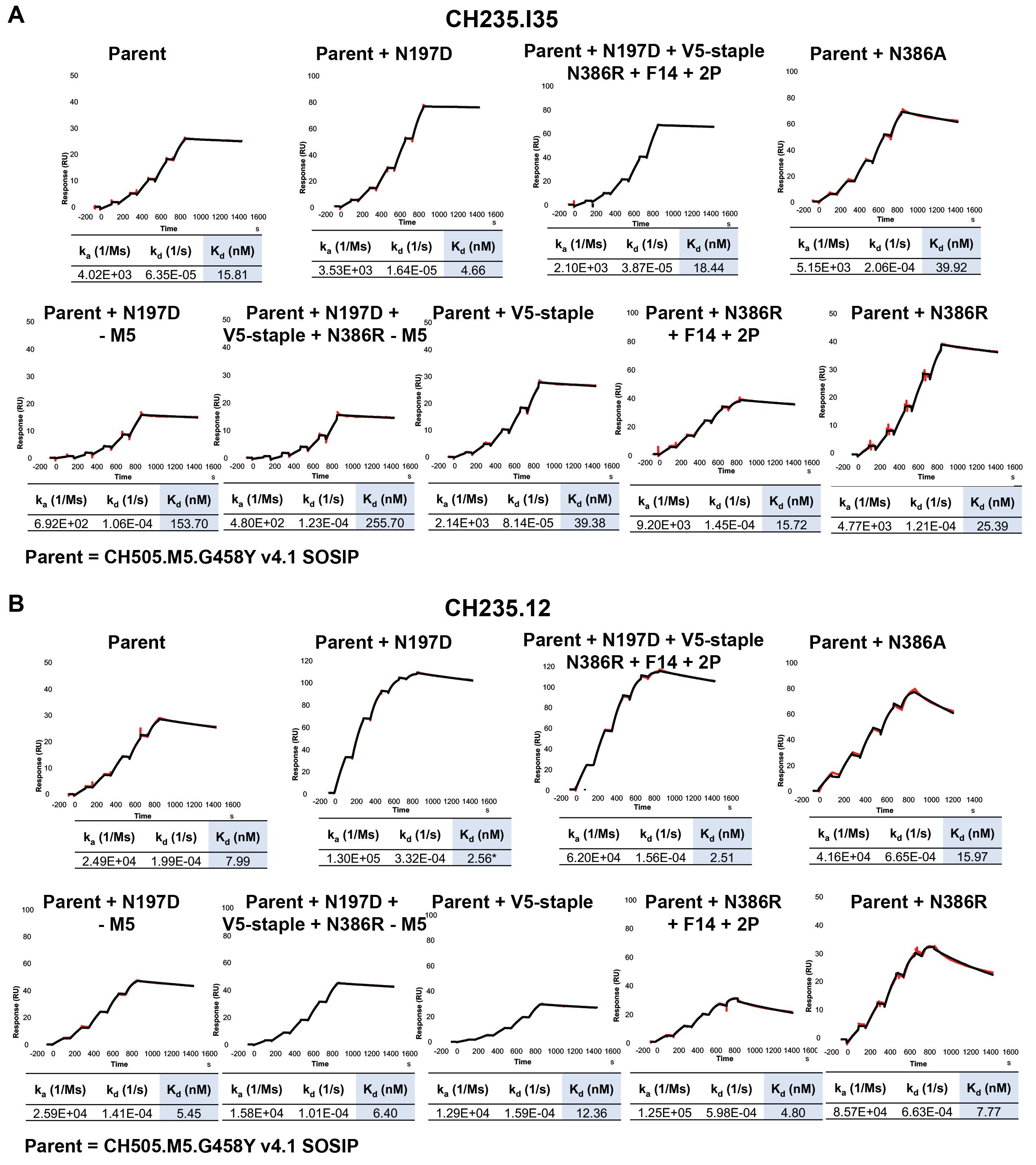

Supplemental Figure 25

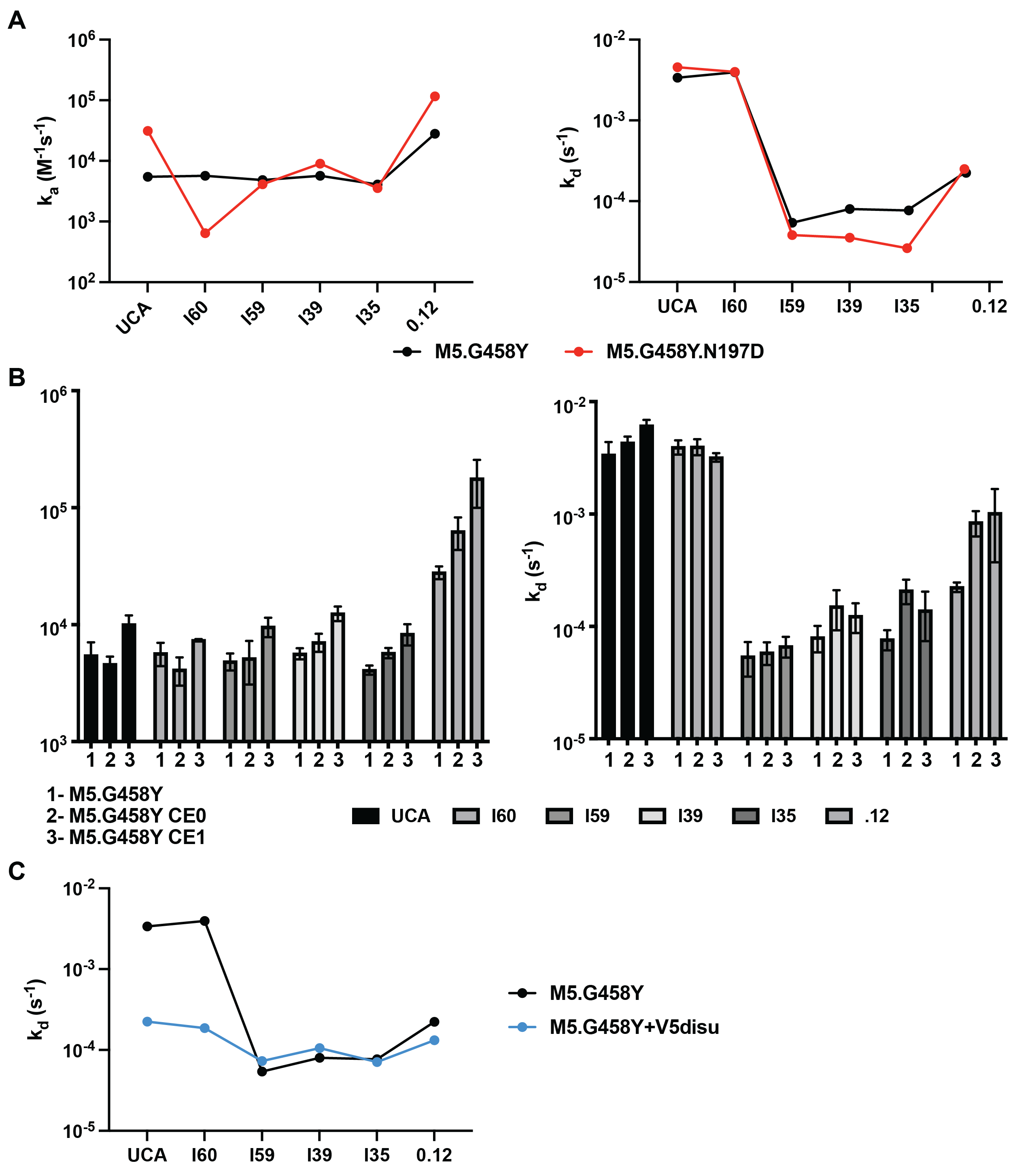

Supplemental Figure 26

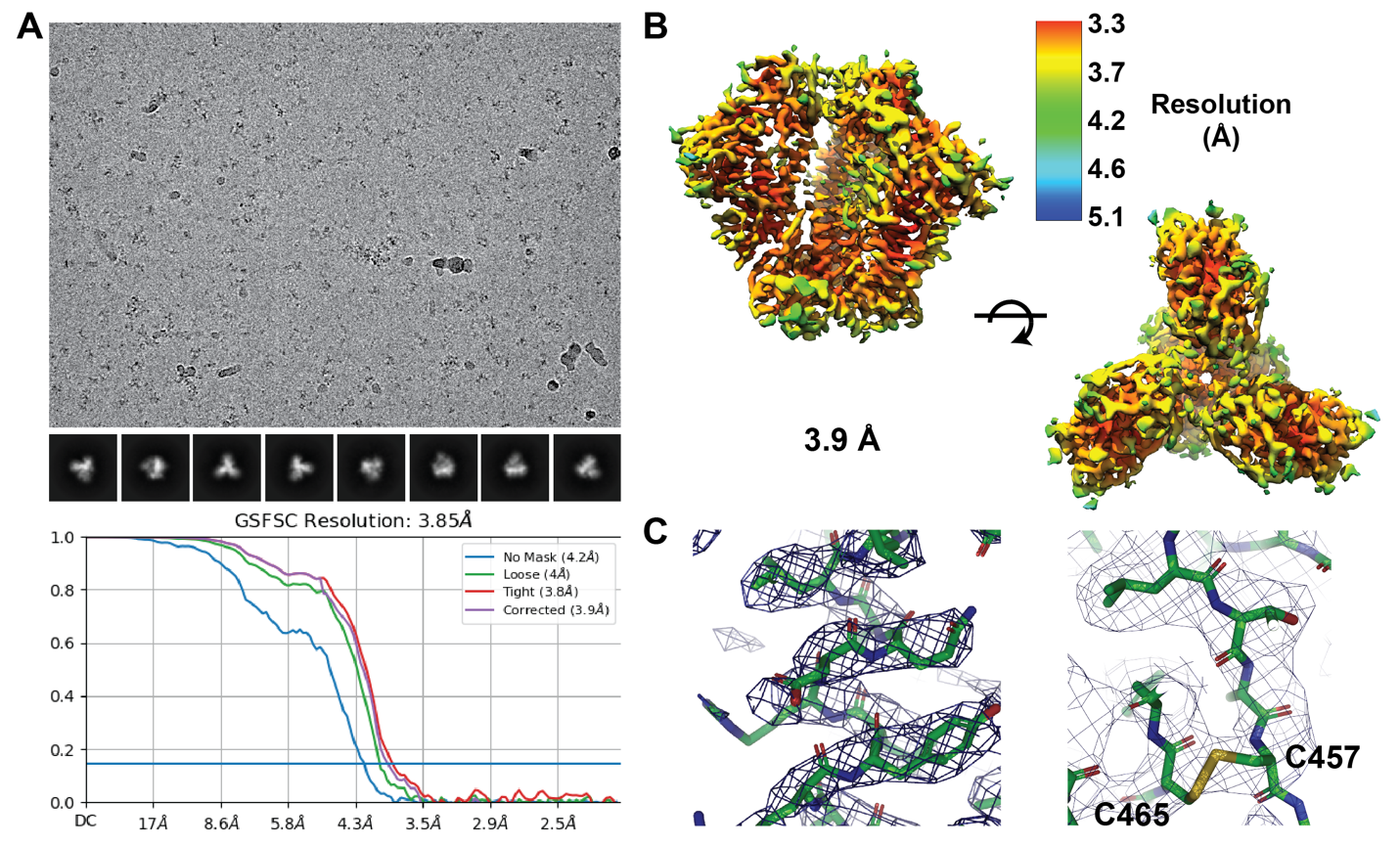

Supplemental Figure 27

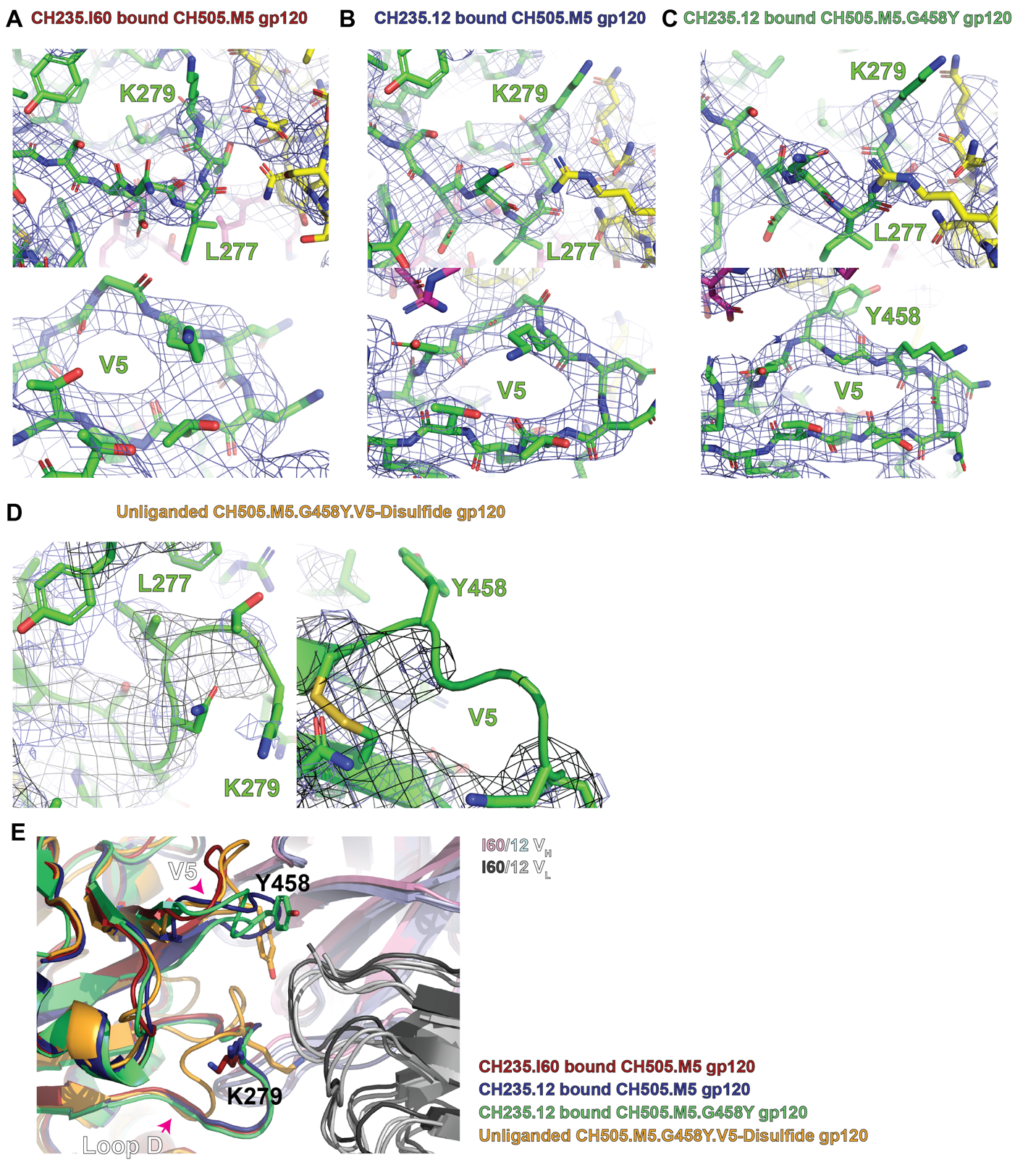

Supplemental Figure 28

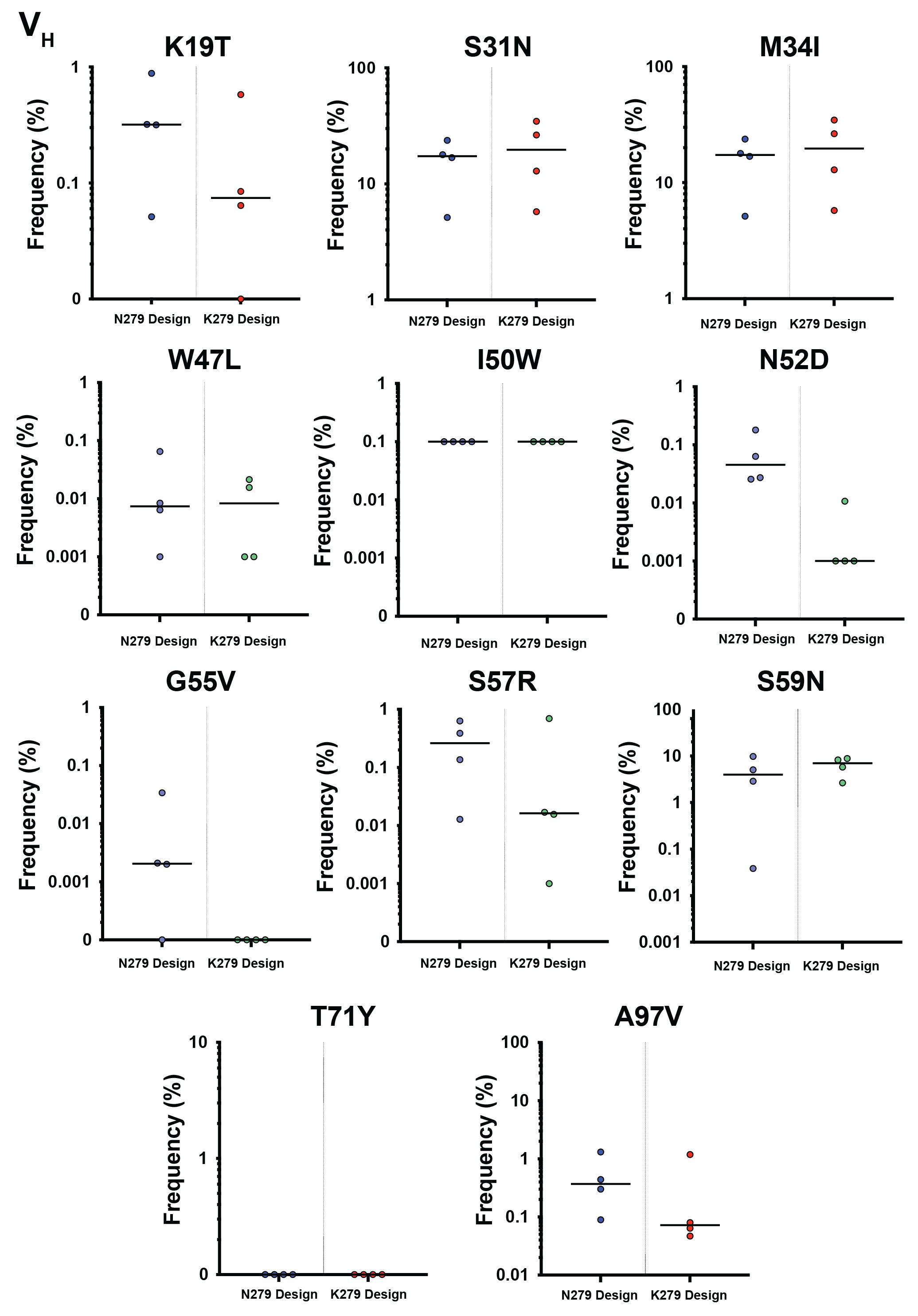

Supplemental Figure 29

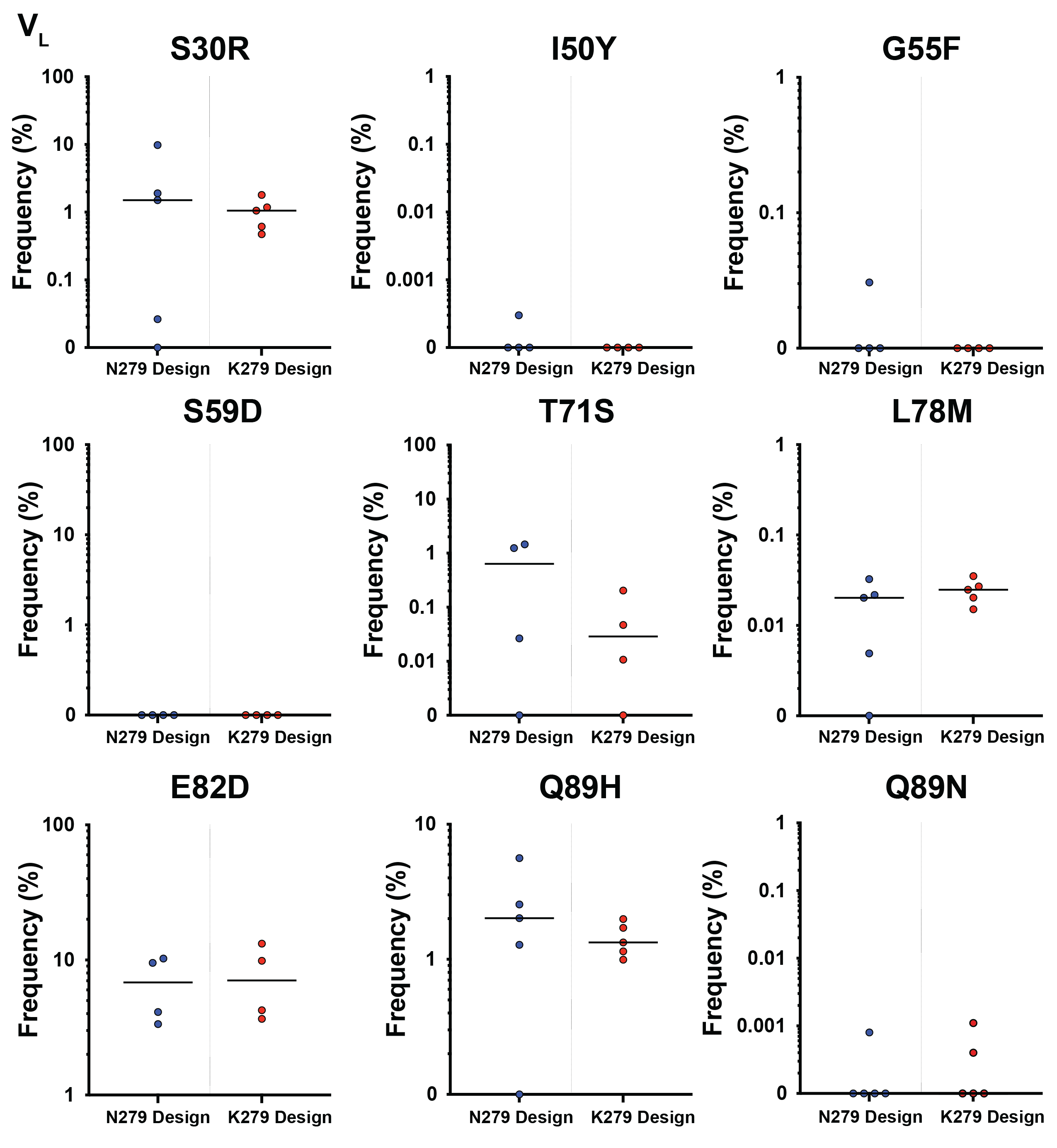

Supplemental Figure 30

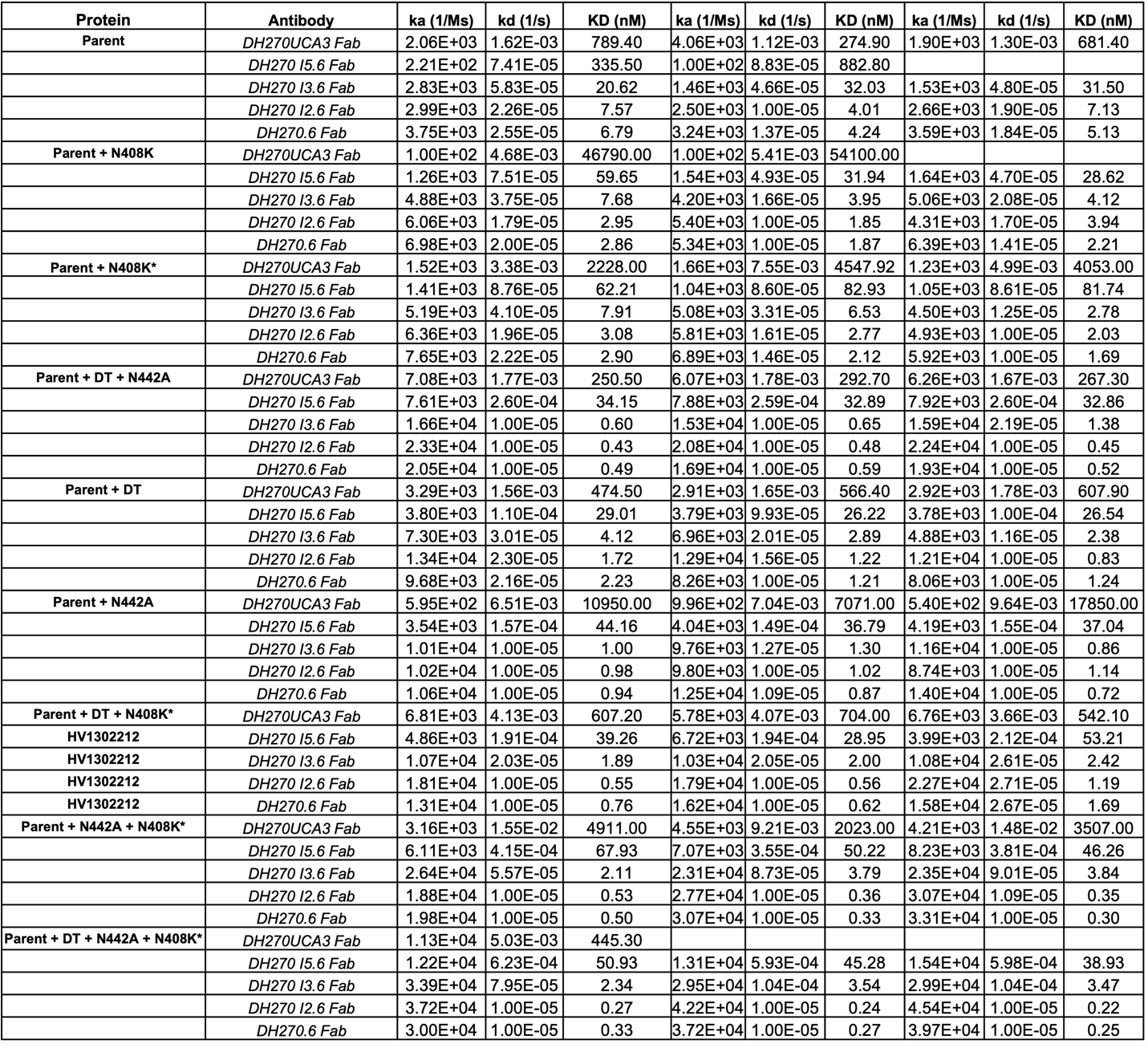

Supplemental Table 1

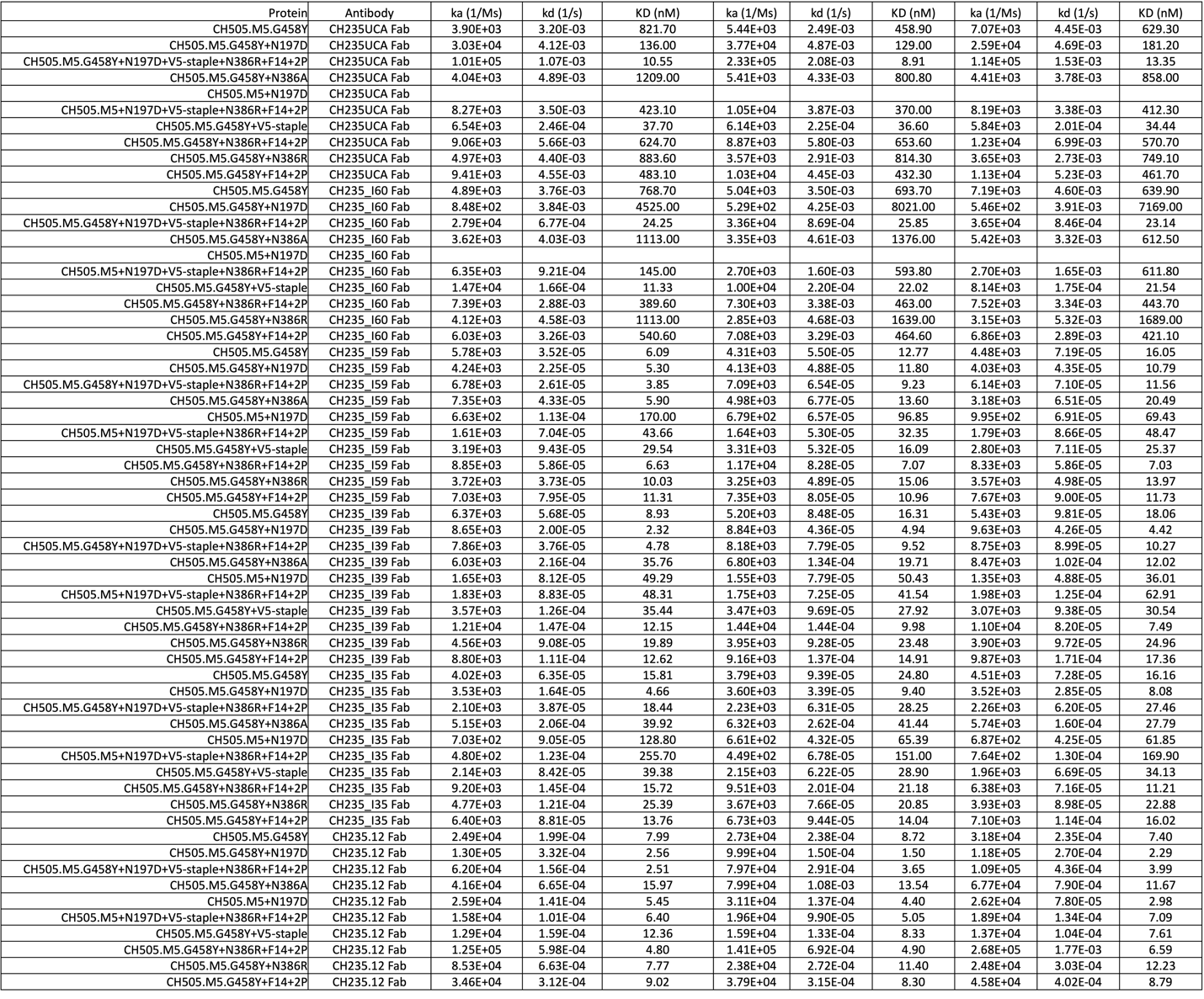

Supplemental Table 2

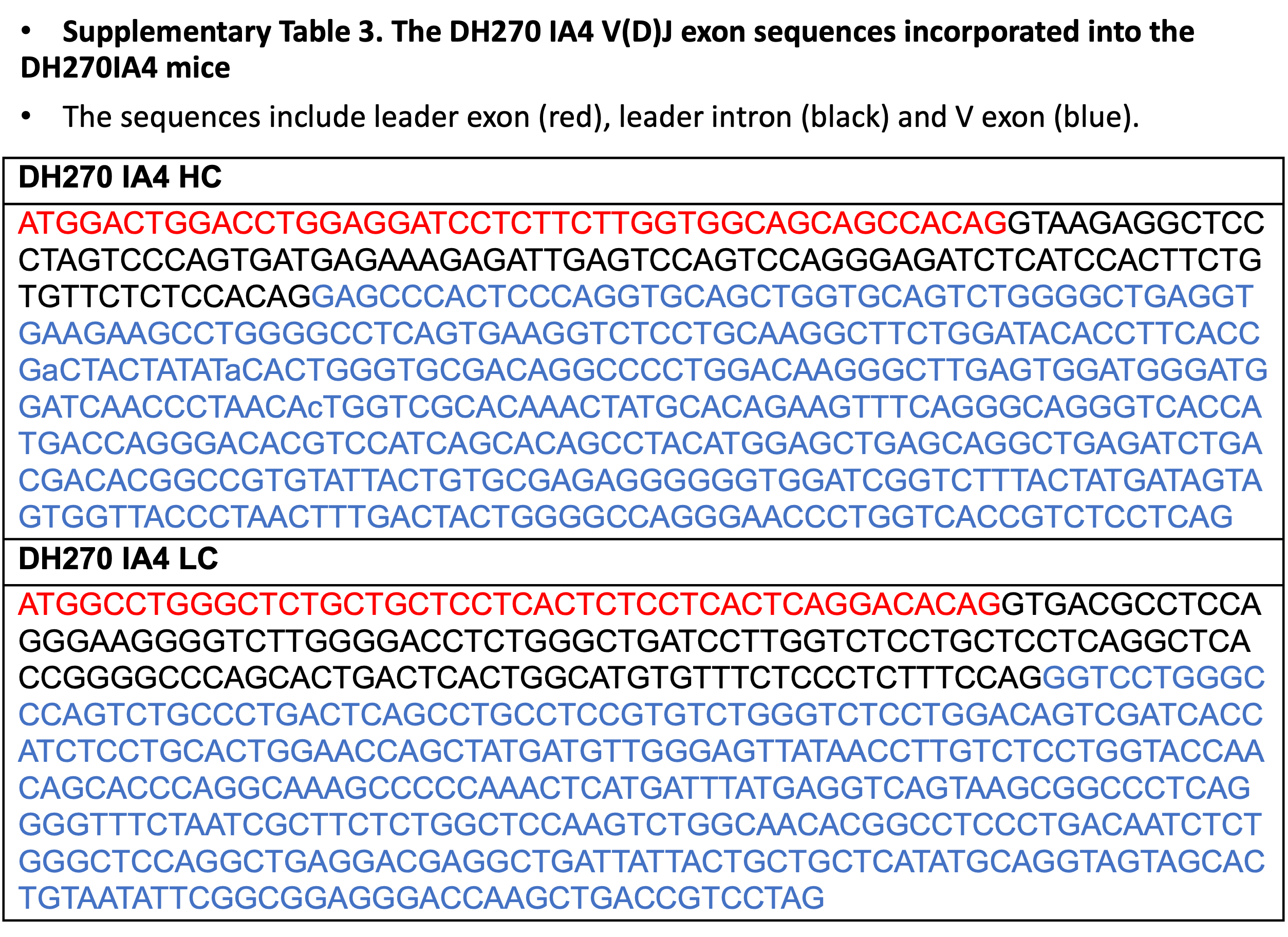

Supplemental Table 3

**Supplemental Figure 1. A)** Cartoon representation of the truncated, closed state gp120 domain overlayed with fifty representative states (ribbon; sheets, helices, and loops colors are orange, blue and cyan/green, respectively). **B)** (left) Plot of the RMSD for all atoms or sheet and helix atoms only for each final state in the five-hundred ~one-microsecond simulation set compared to the initial state. (right) Table of statistics for the RMSD data. **C)** Ribbon representation for all aligned final states.

**Supplemental Figure 2. A)** The 400 state DH270.6 structural ensemble (grey) relative to the gp120T produced using molecular dynamics simulation. The Fv HCDR3 (blue) and N332-glycan (translucent purple spheres) highlight the relative orientation of the antibody. **B)** (left) The N332-glycan D-arm contacts with the DH270 heavy and light chain residues. (right) Histogram of initiation state N332-glycan contacts.

**Supplemental Figure 3.** Initial states for each of one thousand independent simulations from uncoupled interactive pairs (gp120-DH270.6 Fv). The gp120 (light blue) domain is colored according to V1/V2 (green), V3 (red), and V4 (orange). The N332-glycan is represented as translucent grey spheres. Each initial state Fv is represented as a centroid bead (cyan) and an arrow (purple) representing the vector connecting the centroid with a residue in HCDR3. Each panel represents a different viewing orientation.

**Supplemental Figure 4. A)** Counts histogram for DH270.6 simulations in the projected TICA space (dimensions 1 and 2). **B)** Implied timescales plot for the clustered simulation set. **C)** Chapman-Kolmogorov plots for the coarse-grained Markov state model.

**Supplemental Figure 5. A)** Free energy surface calculated from the MSM plotted along TICA dimensions 1 and 2 with (right) and without (left) K-means cluster centers (colors according to macro-states). **B)** Free energy surface calculated from the MSM plotted along the DH270.6 Fv VH R57 to gp120 I322 minimum distance and the RMSD relative to the bound state with (right) and without (left) K-means cluster centers (colors according to macro-states). **C)** Table indicating statistics from the model for each state, including the RMSD to the bound state.

**Supplemental Figure 6. A)** Committer distribution for the states in the MSM. Inset table provides key rate-related statistics from the model. **B)** Path statistics describing the connectivity, flux, and contributions of differing paths to the bound state.

**Supplemental Figure 7. A)** (left) DH270.6 ribbon ensembles depicting fifty representative states from each MSM macro-state. The HCDR3 loop is highlighted in bright green. Elements highlighted as surfaces in gp120 (yellow; V1/V2=green, V3=red) include the V1 loop (magenta), V4 (purple), and the GDIK motif (grey). The N332-glycan is represented depicted as red sticks. (right) View of gp120 depicting fifty representative states of the V1 loop (ribbon) from the MSM. **B)** Overlay of the prebound State 4 and bound state Fv ensembles. **C)** Overlay of the prebound State 4 and bound state V1 ensembles. **D)** View of State 2 showing the position of the bound state (cartoon Fv), the Fv ensemble (ribbon), and the N442-glycan ensemble (purple sticks).

**Supplemental Figure 8. A)** Representative SPR curves for binding of the DH270 UCA to the parent and combinations of DH270 clonal lineage targeting Env SOSIP gp140 designs. **B)** Representative SPR curves for binding of the DH270 I5 intermediate to the parent and combinations of DH270 clonal lineage targeting Env SOSIP gp140 designs.

**Supplemental Figure 9. A)** Representative SPR curves for binding of the DH270 I3 intermediate to the parent and combinations of DH270 clonal lineage targeting Env SOSIP gp140 designs. **B)** Representative SPR curves for binding of the DH270 I2 intermediate to the parent and combinations of DH270 clonal lineage targeting Env SOSIP gp140 designs.

**Supplemental Figure 10.** Representative SPR curves for binding of the mature DH270.6 bnAb to the parent and combinations of DH270 clonal lineage targeting Env SOSIP gp140 designs.

**Supplemental Figure 11.** Glycan 442 is associated with resistance to DH270 IA4 (or I5.6) and IA2 (or I3.6) intermediates. By analyzing Env signatures associated with breadth gain in the DH270 lineage (Wagh, Korber et al. submitted), we identified 442 glycan is associated with resistance to IA4 and IA2 intermediates. (A) Neutralization data for DH270.6 and intermediates IA4, IA2 and IA1 (similar to I5.6, I3.6 and I2.6, respectively) tested against heterologous viruses(*1*). A panel of 205 heterologous pseudoviruses was tested against these antibodies, of which 113 viruses (55%) were neutralized by DH270.6; data and analyses for only these viruses are shown. IC50 titers are shown with red to yellow to green color scale indicating more to less potent neutralization, and black cells indicate resistant viruses (IC50 > 50µg/ml). The sequence at HXB2 position 442 is shown for each virus; “NxST” indicates Asn at 442 in a potential N-linked glycan sequon (PNGS), while “N” indicates Asn is not in the context of the sequon. (B) Percentage of N442 glycan containing viruses in IA4 sensitive (IC_50_ < 50µg/ml) and resistant (IC_50_ > 50µg/ml) viruses is shown in blue. Percentage of viruses not containing N442 glycan shown in black. Statistical difference was calculated using Fisher’s exact test comparing the number of viruses with and without N442 glycan in IA4 sensitive, and p-value is shown on the top. (C) Same as panel (B), except sensitive and resistant viruses to IA2 are used. (D) Difference in IA2 IC_50_ titers against viruses with N442 glycan (blue) and without N442 glycan (black). Thick green lines indicate medians, and thin green lines indicate 25^th^ and 75^th^ percentiles. Statistical difference was calculated using Wilcoxon rank sum test and the p-value is indicated. These results indicate that IA4 has significantly limited ability to neutralize N442 glycan containing heterologous viruses, but IA2 does not suffer from this roadblock. However, IA2 is still significantly less potent against N442 glycan containing heterologous viruses.

**Supplemental Figure 12.** *In vitro selection of N442D substitution in HIV-1 Env that improves DH270 UCA association rate for envelope.* **A)** (left) Flow cytometry dot plot of DH270 UCA binding to cells expressing an envelope (Env) variant library derived from CH848.D0949.10.17 N133D/ N138T/ E169K gp140. The envelope library contained single mutations at each position within gp120 and were enriched with four rounds of sorting with DH270 UCA. (right) Flow cytometry dot plot of binding to DH270 UCA binding to cells expressing unmutated CH848.D0949.10.17 N133D/ N138T/ E169K gp140. **B)** PacBio sequencing results of enriched DH270 UCA-selected envelope variant expressing cells shown in A. The red and blue bars show the envelope mutations chosen to construct a 2nd generation combinatorial mutation envelope library. **C)** Fluorescence activated cell sorting (FACS) dot plot of cells expressing the 2nd generation Env library that bound to DH270 UCA. **D)** Envelope mutation enrichment after FACS of DH270 UCA-reactive cells from the 2nd generation library. Change in frequency is represented by the frequency of sequences with the mutation after DH270 UCA sorting subtracted by the frequency of the sequence in the library prior to sorting. This difference in frequency was then divided by the initial frequency of the mutation in the library prior to sorting. Enrichment determined from Sanger sequencing of random colonies (Left) and MiSeq sequencing of bulk sorted cells (Right) from the 2nd generation combinatorial mutagenesis library. Blue bar shows N442D increased in frequency after DH270 UCA sorting. **E)** DH270 UCA binding kinetics to CH848.D0949.10.17 N133D/ N138T/ E169K /K327 gp140 with (K327/N442D) or without N442D (K327) determined with Bio-Layer Interferometry. **F)** Comparison of the DH270 UCA association rate with CH848.D0949.10.17 N133D/ N138T/ E169K /K327 gp140 with

**Supplemental Figure 13. A)** (top) Representative cryo-EM micrograph (middle) 2D-class averages for the DE3 design bound to the DH270.6 Fab. (bottom) FSC plot for the DE3 design bound to the DH270.6 Fab map reconstruction **B)** Local resolution map. **C)** (left) Representative map-to-model fit in the region of the gp41 3-helix bundle. (right) Representative map-to-model fit in the region of the GDIK motif and the N442A mutation site. **D)** Alignment between cryo-EM structures of the parent CH8483-d949 Env SOSIP (green) and DE3 design (purple) bound to the DH270.6 Fab.

**Supplemental Figure 14. A)** (top) Representative cryo-EM micrograph (middle) 2D-class averages for the DE3 design bound to the VRC01 Fab. (bottom) FSC plot for the DE3 design bound to the VRC01 Fab map reconstruction **B)** Local resolution map. **C)** (left) Representative map-to-model fit in the region of the gp41 3-helix bundle. (right) Representative map-to-model fit in the region of the N332-glycan and the N442A mutation site. **D)** Alignment of the unmutated CH848.10.17 Env SOSIP, VRC01 bound structure with the DE3 design bound to the VRC01 highlighting the DH270.6 epitope. **E)** Difference map (green) for the VRC01 bound parent vs. the VRC01 bound design overlayed with the VRC01 bound design map.

**Supplemental Figure 15. A)** Panel of pseudo-virus IC50 neutralization titers for sera from IA4 knock in mice immunized with either an N-glycan hole-filled CH848-d949 SOSIP trimer (N=12) or the DE3 design(N=13). Statistically significant differences are indicated in cyan (Unpaired Mann-Whitney test). **B)** Mutation frequencies for the IA4 key improbable heavy chain mutation G110Y and light chain L48Y. Bars indicate the mean frequency. C) Sequence alignments for R98T and/or L48Y containing DH270.IA4 sequences from B cells isolated from immunized mice.

**Supplemental Figure 16. A)** Initial CH505 gp120_T_ vs. CH235.12 Fv structural state used for simulation of the association process. Man5 glycans are shown as multi-color translucent spheres. **B)** Distribution of CH235.12 Fv positions relative to the gp120_T_ after 200 independent, 250 ns simulations. **C)** RMSD trajectories (dark blue) of three representative transitions from encounter states to the bound state. Distances between CH235.12 heavy chain R56 and gp120 I371 (solid black) and light chain W94 and gp120T K282 α-carbons show the evolution of heavy and light chain positions relative to the gp120T prior to reaching the pre-bound state. **D)** Representative encounter to pre-bound transition showing rotation of the CH235.12 Fv (fade from red to white to blue) about the CD4bs loop (cyan). **E)** Frames from the representative simulation in a CD4bs loop interactive encounter transition (top) and the final, VL to loop D proximal pre-bound state.

**Supplemental Figure 17. A)** Counts histogram for CH235.12 vs. CH505 gp120_T_ simulations in the projected TICA space (dimensions 1 and 2). **B)** Implied timescales plot for the clustered simulation set. **C)** Chapman-Kolmogorov plots for the coarse-grained Markov state model.

**Supplemental Figure 18. A)** Free energy surface calculated from the CH235.12 vs. CH505 gp120_T_ MSM plotted along TICA dimensions 1 and 2 with (right) and without (left) K-means cluster centers (colors according to macro-states). **B)** Free energy surface calculated from the MSM plotted along CH235.12 Fv VH R56 to gp120 I371 minimum distance and the RMSD relative to the bound state with (right) and without (left) K-means cluster centers (colors according to macro-states). **C) Table** indicating statistics from the model for each state, including the RMSD to the bound state.

**Supplemental Figure 19. A)** Committer distribution for the states in the CH235.12 vs. CH505 gp120_T_ MSM. The Inset table provides key rate-related statistics from the model. **B)** Path statistics describing the connectivity, flux, and contributions of differing paths to the bound state.

**Supplemental Figure 20. A)** Structural ensembles of states for the CH235.12 association model relative to an HIV-1 Env trimer structure (PDB ID 6UDA). The simulation gp120 domain was aligned with the trimer structure to show the proximity of the ensemble to the adjacent protomer gp120.

**Supplemental Figure 21. A)** Ensemble of CH235.12 states for State 3 of the model positioned relative to the initial simulation gp120 conformation. The HCDR3 loop is highlighted in green. The gp120 CD4bs loop, V5 loop, and D loop are represented as backbone atom surfaces. The N386 and N197 glycans are shown in stick representation **B)** A representative state from the State 3 ensemble depicted as in (A). **C)** The State 3 ensemble as in (A) with the V5 loop backbone ensemble shown as black sticks overlayed on the initial simulation V5 conformation translucent surface.

**Supplemental Figure 22. A)** (top) Representative cryo-EM micrograph (middle) 2D-class averages for the CH505.M5 SOSIP bound to the CH235.I60 Fab. (bottom) FSC plot for the CH505.M5 SOSIP bound to the CH235.I60 Fab map reconstruction **B)** Local resolution map for CH505.M5 SOSIP bound to the CH235.I60. **C)** (left) Representative map-to-model fit in the region of the gp41 3-helix bundle. (right) Representative map-to-model fit in the loop D region. **D)** (top) Representative cryo-EM micrograph (middle) 2D-class averages for the CH505.M5.G458Y SOSIP bound to the CH235.12 Fab. (bottom) FSC plot for the CH505.M5.G458Y SOSIP bound to the CH235.12 Fab map reconstruction **E)** Local resolution map for CH505.M5.G458Y SOSIP bound to the CH235.12. **F)** (left) Representative map-to-model fit in the region of the gp41 3-helix bundle. (right) Representative map-to-model fit in the loop D region.

**Supplemental Figure 23. A)** Representative SPR curves for binding of the CH235 UCA to the parent and combinations of CH235 clonal lineage targeting Env SOSIP gp140 designs. **B)** Representative SPR curves for binding of the CH235.I60 intermediate to the parent and combinations of CH235 clonal lineage targeting Env SOSIP gp140 designs.

**Supplemental Figure 24. A)** Representative SPR curves for binding of the CH235.I59 intermediate to the parent and combinations of CH235 clonal lineage targeting Env SOSIP gp140 designs. **B)** Representative SPR curves for binding of the CH235.I39 intermediate to the parent and combinations of CH235 clonal lineage targeting Env SOSIP gp140 designs.

**Supplemental Figure 25. A)** Representative SPR curves for binding of the CH235.I35 intermediate to the parent and combinations of CH235 clonal lineage targeting Env SOSIP gp140 designs. **B)** Representative SPR curves for binding of the mature CH235.12 to the parent and combinations of CH235 clonal lineage targeting Env SOSIP gp140 designs.

**Supplemental Figure 26. A)** (left) Association rates for the CH505.M5.G458Y N197D construct interacting with CH235 clonal lineage members. (right) Dissociation rates for the CH505.M5.G458Y N197D construct interacting with CH235 clonal lineage members. **B)** (left) Association rates for the CE0 and CE1 designs interacting with CH235 clonal lineage members. (right) Dissociation rates for the CE0 and CE1 designs interacting with CH235 clonal lineage members. **C)** Association rates for the CE2 V5-disulfide stapled design interacting with CH235 clonal lineage members.

**Supplemental Figure 27. A)** (top) Representative cryo-EM micrograph (middle) 2D-class averages for the unliganded CH505.M5.G458Y V5 disulfide stapled design. (bottom) FSC plot for the unliganded CH505.M5.G458Y.N197D design map reconstruction **B)** Local resolution map. **C)** (left) Representative map-to-model fit in the region of the gp41 3-helix bundle. (right) Representative map-to-model fit for the V5 loop highlighting the position of the introduced C547-C465 disulfide.

**Supplemental Figure 28. A)** The map to model fit for the CH235.I60 bound CH505.M5 trimer depicting the conformation for the (top) D loop and (bottom) V5 loop. **B)** The map to model fit for the CH235.12 bound CH505.M5 trimer depicting the conformation for the (top) D loop and (bottom) V5 loop. **C)** The map to model fit for the CH235.12 bound CH505.M5.G458Y trimer depicting the conformation for the (top) D loop and (bottom) V5 loop. **D)** The map to model fit for the unliganded CH505.M5.G458Y V5 disulfide stapled design depicting the conformation for the (left) D loop and (right) V5 loop. **E)** Alignment of each structure’s gp120 domain highlighting differences in the D loop and V5 loop conformations.

**Supplemental Figure 29.** Heavy chain mutation frequencies for key CH235 lineage mutations in the immunized CH235.UCA knock-in mice.

**Supplemental Figure 30.** Light chain mutation frequencies for key CH235 lineage mutations in the immunized CH235.UCA knock-in mice.

1.	M. Bonsignori *et al.*, Staged induction of HIV-1 glycan-dependent broadly neutralizing antibodies. *Sci Transl Med* **9**, (2017).
